# Resident Cardiac Macrophages Mediate Adaptive Myocardial Remodeling

**DOI:** 10.1101/2021.01.28.428724

**Authors:** Nicole R. Wong, Jay Mohan, Benjamin J Kopecky, Shuchi Guo, Lixia Du, Jamison Leid, Oleksandr Dmytrenko, Hannah Luehmann, Geetika Bajpai, Laura Ewald, Lauren Bell, Nikhil Patel, Inessa Lokshina, Andrea Bredemeyer, Carla J. Weinheimer, Jessica M. Nigro, Attila Kovacs, Sachio Morimoto, Peter O. Bayguinov, Max. R. Fisher, James A.J. Fitzpatrick, Slava Epelman, Daniel Kreisel, Rajan Sah, Yongjian Liu, Hongzhen Hu, Kory J. Lavine

## Abstract

Cardiac macrophages represent a heterogeneous cell population with distinct origins, dynamics, and functions. Recent studies have revealed that C-C Chemokine Receptor 2 positive (CCR2+) macrophages derived from infiltrating monocytes regulate myocardial inflammation and heart failure pathogenesis. Comparatively little is known about the functions of tissue resident (CCR2−) macrophages. Herein, we identify an essential role for CCR2− macrophages in the chronically failing heart. Depletion of CCR2− macrophages in mice with dilated cardiomyopathy accelerated mortality and impaired ventricular remodeling and coronary angiogenesis, adaptive changes necessary to maintain cardiac output in the setting of reduced cardiac contractility. Mechanistically, CCR2− macrophages interacted with neighboring cardiomyocytes via focal adhesion complexes and were activated in response to mechanical stretch through a transient receptor potential vanilloid 4 (TRPV4) dependent pathway that controlled growth factor expression. These findings establish a role for tissue resident macrophages in adaptive cardiac remodeling and introduce a new mechanism of cardiac macrophage activation.

## Introduction

Paradigm shifting studies have demonstrated surprising heterogeneity among macrophage populations. It is now widely recognized that macrophages arise from distinct developmental origins including extraembryonic (yolk sac) and definitive hematopoietic progenitors and are maintained through differing mechanisms (Davies et al., 2013; Epelman et al., 2014b; Hashimoto et al., 2013; Hettinger et al., 2013; Hoeffel et al., 2015; Wynn et al., 2013; Yona et al., 2013). For example, microglia found within the brain are derived from extraembryonic hematopoiesis and maintained through local proliferation independent of monocyte input (Ginhoux et al., 2010), while macrophages that reside within the intestine are derived from definitive hematopoietic progenitors and are continually replenished by recruited monocytes (Bain et al., 2013). Most organs including the heart, lung, liver, and skin contain admixtures of distinct macrophage subsets with differing development origins, morphologies, tissue localizations, and population dynamics (Epelman et al., 2014a; Guilliams et al., 2013; Hoeffel et al., 2012; Theret et al., 2019). These findings have raised the possibility that individual macrophage subsets may execute unique and context specific functions.

Beyond regulating inflammatory signaling, macrophages contribute important functions to shaping and remodeling tissues throughout development and adulthood.

Macrophages are essential for the development and maturation of the nervous and vascular systems, contribute to bone and tooth morphogenesis, and clear remnants of cells that undergo programmed cell death within the embryo (Fantin et al., 2010; Munoz-Espin et al., 2013; Parkhurst et al., 2013; Storer et al., 2013; Theret et al., 2019). Many of these macrophage populations originate from embryonic progenitors, reside within tissues for prolonged periods of time, and are referred to as tissue resident macrophages. Tissue resident macrophages also orchestrate regeneration of cardiac and appendage tissue following amputation or other forms of injury (Aurora et al., 2014; Godwin et al., 2017; Godwin et al., 2013; Lavine et al., 2014; Petrie et al., 2014). In the adult organism, tissue resident macrophages play key roles in maintaining organ homeostasis and physiology including iron metabolism and transport, regulation of hematopoiesis, clearance of airway debris and surfactant, facilitation of electrical impulses through cardiac conduction tissue (Chow et al., 2011; Hashimoto et al., 2013; Hulsmans et al., 2017; Soares and Hamza, 2016). However, little is understood regarding the functions of tissue resident macrophages in the context of chronic disease.

Cardiac tissue remodeling is a widely recognized response to reductions in contractility, hemodynamic loading, or pathological insults to the heart. In these contexts, the heart undergoes robust geometric changes characterized by concentric thickening and dilation of the left ventricle (LV). While initially beneficial through reductions in LV wall stress, progressive myocardial hypertrophy and enlargement contributes to the development and progression of heart failure through cardiomyocyte cell death, further reduction in contractile function, and interstitial fibrosis. This process is referred to as adverse/pathological LV remodeling and is commonly observed across numerous cardiac pathologies such as myocardial infarction, viral myocarditis, and nonischemic cardiomyopathies (Burchfield et al., 2013; Xie et al., 2013).

It is important to note that not all forms of cardiac remodeling are maladaptive. Adaptive changes such as LV chamber enlargement and eccentric hypertrophy represent physiological adaptations to exercise conditioning. This physiological form of hypertrophy is associated with coronary angiogenesis, cardiomyocyte lengthening, and is distinct from adverse remodeling at the transcriptional level. Interstitial fibrosis and contractile dysfunction are typically not present in physiological hypertrophy (Nakamura and Sadoshima, 2018). At present, the precise cellular and molecular mechanisms that orchestrate adaptive cardiac remodeling are incompletely understood. Intriguingly, features of adverse and adaptive LV remodeling coexist in patients with chronic heart failure highlighting the clinical relevance of understanding each form of cardiac tissue remodeling (Cohn et al., 2000; Patel et al., 2017).

Given their functions during heart development, cardiac tissue resident macrophages represent an attractive cell type that may govern remodeling of myocardial tissue in response to hemodynamic perturbations and/or chronic disease. Under homeostatic conditions, the adult heart contains a heterogeneous population of tissue resident macrophages that can be divided into two functionally distinct subsets based on the cell surface expression of C-C chemokine receptor 2 (CCR2) (Epelman et al., 2014a). CCR2+ macrophages are derived from definitive hematopoietic progenitors, replenished by monocyte recruitment and subsequent proliferation, and function to initiate inflammatory cascades. In response to cardiomyocyte death, CCR2+ macrophages produce inflammatory cytokines, orchestrate the recruitment of neutrophils and monocytes, generate damaging oxidative productions, and consequently, contribute to the progression of heart failure through collateral myocardial injury and adverse cardiac remodeling (Bajpai et al., 2019; Lavine et al., 2014; Li et al., 2016; Patel et al., 2018). Clinically, CCR2+ macrophage abundance is predictive of and associated with adverse LV remodeling in advanced heart failure patients (Bajpai et al., 2018), and thus, represent a target for future immunomodulatory therapies. CCR2− macrophages are largely derived from embryonic (yolk sac and fetal liver) hematopoietic progenitors and are maintained independent of monocyte recruitment through local proliferation. CCR2− macrophages orchestrate the maturation of the developing coronary vasculature and neonatal heart regeneration (Lavine et al., 2014; Leid et al., 2016). In the adult heart, CCR2− macrophages appear to suppress inflammatory responses following acute myocardial injury (Bajpai et al., 2019). Whether CCR2− macrophages have similar reparative functions in the context of chronic heart failure is unknown.

Herein, we test the hypothesis that tissue resident CCR2− macrophages are involved in adaptive remodeling of the chronically failing heart. By employing a mouse model of dilated cardiomyopathy harboring a causative human mutation, we define the composition and dynamics of macrophages residing within the chronically failing heart. Through selective cell depletion studies, we reveal an essential role for CCR2− macrophages in adaptive LV remodeling, coronary angiogenesis, maintenance of cardiac output, and survival of mice with dilated cardiomyopathy. Furthermore, we provide evidence that mechanical sensing through a transient receptor potential vanilloid 4 (TRPV4) dependent pathway constitutes a novel mechanism controlling growth factor expression in tissue resident cardiac macrophages.

## Results

### Cardiac Macrophage Heterogeneity in Dilated Cardiomyopathy

To investigate cardiac macrophage composition and function in chronic heart failure, we chose to focus on a mouse model of human dilated cardiomyopathy. Previously, knock-in mice were generated that harbor a causative mutation (ΔK210) in the endogenous Troponin T2 (Tnnt2) locus (Du et al., 2007). The Tnnt2^ΔK210^ mutation has been identified in numerous cohorts of familial and sporadic adult and pediatric dilated cardiomyopathy patients and is considered clinically as a pathogenic variant (McNally and Mestroni, 2017). Incorporation of the Tnnt2^ΔK210^ mutant protein into sarcomeres leads to reduced thin filament calcium sensitivity and cardiomyocyte contractility (Clippinger et al., 2019; Morimoto et al., 2002).

Consistent with previous reports, homozygous mice (Tnnt2^ΔK210/ΔK210^) develop a dilated cardiomyopathy with profound LV remodeling (dilation and hypertrophy), reduced LV function (ejection fraction), and early mortality (**Fig. 1A-B and Fig. S1A**). Serial echocardiography revealed that Tnnt2^ΔK210/ΔK210^ mice are born with reduced LV ejection fraction reflective of intrinsic impairment in cardiomyocyte contractility. LV remodeling (dilation and eccentric hypertrophy) was not evident until 2 weeks of age and increased progressively over time (**Fig. S1B**). These findings suggest that early LV remodeling may represent a compensatory response to reduced LV contractility.

**Figure 1.**
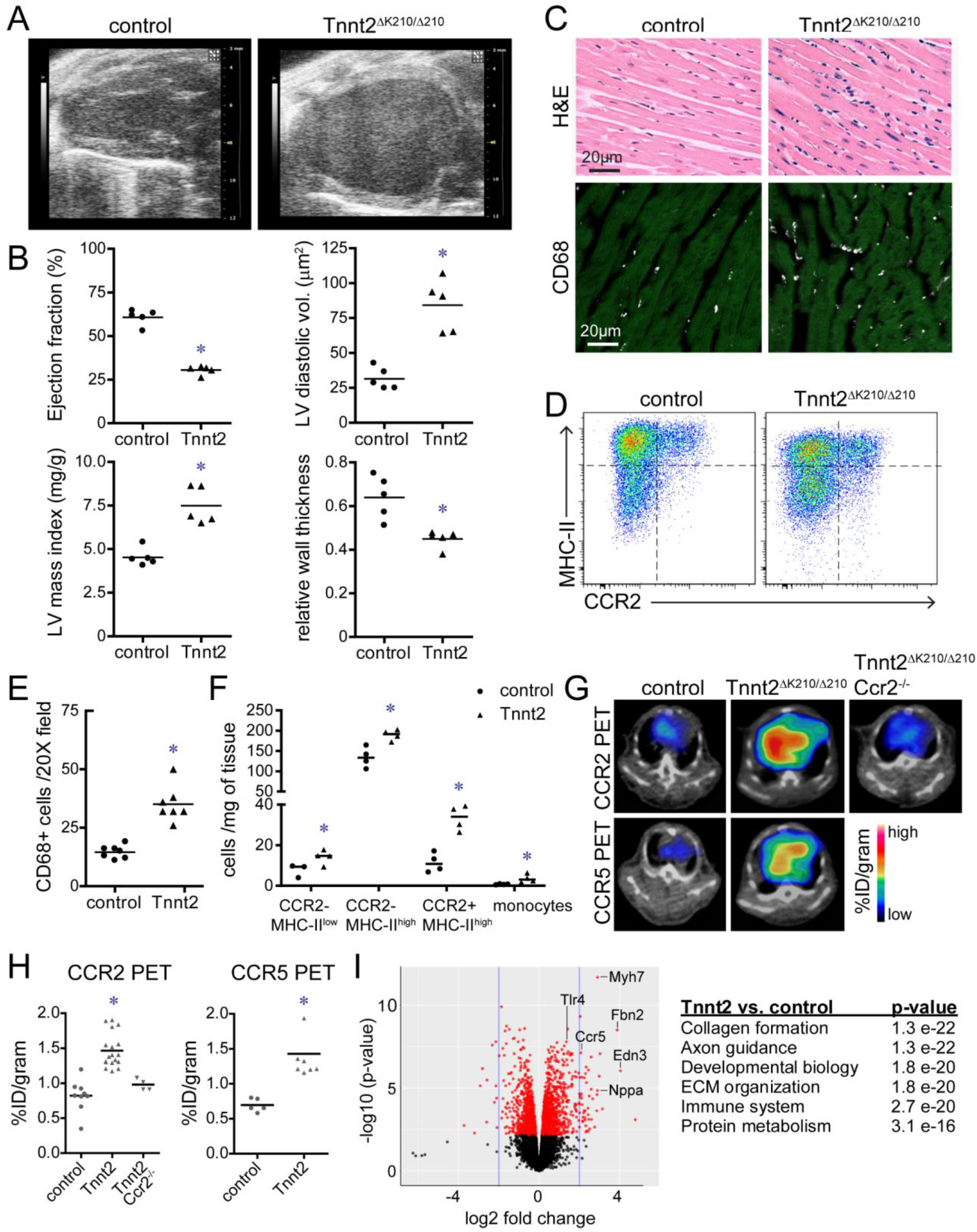
Expansion of cardiac macrophages in a mouse model of dilated cardiomyopathy. **A**, End-diastolic echocardiographic images of 8-week-old control and Tnnt2^ΔK210/ΔK210^ mice. **B**, Quantification of ejection fraction, LV diastolic dimension, LV mass index, and relative wall thickness. **C**, H&E (top) and immunostaining images (bottom, CD68-white, cardiac actin-green) of 8-week-old control and Tnnt2^ΔK210/ΔK210^ mice revealing expansion of cardiac macrophages in Tnnt2^ΔK210/ΔK210^ mice compared to controls. Representative images from n=7 mice per experimental group. **D**, Representative flow cytometry plots of CD45^+^Ly6G^-^ CD64^+^ macrophages showing increased abundance of CCR2−and CCR2+ macrophages in Tnnt2^ΔK210/ΔK210^ mice compared to controls at 8 weeks of age. n=4 per experimental group. **E-F**, Quantification of CD68 immunostaining and flow cytometry. **G**, PET/CT images of control and Tnnt2^ΔK210/ΔK210^ mice (6-10 weeks of age) using CCR2 and CCR5 tracers showing tracer uptake in the hearts of Tnnt2^ΔK210/ΔK210^ mice. CCR2 signal is absent from Tnnt2^ΔK210/ΔK210^ Ccr2^-/-^ mice compared to controls. n=4-17 per experimental group. PET: positron emission tomography, CT: computed tomography. **H**, Quantification of CCR2 and CCR5 tracer uptake within the heart. **I**, MA plot and pathway analysis of RNA sequencing data comparing control to Tnnt2^ΔK210/ΔK210^ hearts highlighting upregulated expression of transcripts associated with innate immunity in Tnnt2^ΔK210/ΔK210^ hearts. n=5-6 per experimental group. For all panels, each data point denotes individual animals. * denotes p<0.05 (Mann-Whitney test) compared to controls.

To investigate the influence of chronic heart failure on cardiac macrophage abundance and composition, we examined histological sections obtained from control and Tnnt2^ΔK210/ΔK210^ hearts. CD68 immunostaining revealed increased macrophage abundance at both 1 week and 8 weeks of age (**Fig. 1C, E and Fig. S1C-D**). Flow cytometry demonstrated significant shifts in cardiac macrophage composition over time. At 1 week of age, Tnnt2^ΔK210/ΔK210^ hearts displayed distributions of CCR2− macrophages, CCR2+ macrophages, and monocytes that were indistinguishable from controls. At 8 weeks of age, Tnnt2^ΔK210/ΔK210^ hearts contained increased abundance of CCR2+ macrophages, CCR2−MHC-II^low^ macrophages, CCR2−MHC-II^high^ macrophages, and monocytes compared to controls. At 12 weeks of age, there was a progressive increase in the percentage of CCR2+ macrophages and marked increase in monocyte abundance (**Fig. 1D, F, Fig. S1E, Fig. S2**).

To non-invasively assess cardiac macrophage composition in intact mice, we took advantage of a positron emission tomography (PET) based molecular imaging strategy that detects CCR2+ macrophages (CCR2 PET) and total macrophages (CCR5 PET) previously established by our group (Heo et al., 2019; Luehmann et al., 2014). We observed robust CCR2 PET signal in the hearts of Tnnt2^ΔK210/ΔK210^ mice compared to controls. Inclusion of Tnnt2^ΔK210/ΔK210^ Ccr2^-/-^ mice provided evidence of radiotracer specificity and ruled out the possibility that increased CCR2 PET signal was a result of expanded blood pool size. Consistent with greater numbers of total cardiac macrophages in Tnnt2^ΔK210/ΔK210^ hearts, we observed increased CCR5 PET signal in the hearts of Tnnt2^ΔK210/ΔK210^ mice compared to controls (**Fig. 1G-H**).

Analysis of RNA sequencing data comparing control and Tnnt2^ΔK210/ΔK210^ hearts showed marked differences in gene expression that included pathways associated with collagen deposition, extracellular matrix organization, cell migration, and immune functions (**Fig. 1I**). Numerous differentially expressed genes have been implicated in macrophage activation and function (CD44, Mrc2, Nr4a1, Tlr4, Lbp, Csf2ra, Jun, Fos, Irf6, Socs2, Chil1, Ctgf, Gdf15, Ifngr1, Maff) (**Fig. S3**), suggesting a potential role for macrophages in the chronically failing heart.

### Origins and Dynamics of Cardiac Macrophages in Chronic Heart Failure

To delineate the contribution of monocytes to each cardiac macrophage population in the chronically failing heart, we bred Tnnt2^ΔK210/ΔK210^ mice to CCR2^gfp^ knock-in mice to generate the following experimental groups: CCR2^gfp/+^ (control), CCR2^gfp/gfp^ (CCR2 KO), Tnnt2^ΔK210/ΔK210^ CCR2^gfp/+^ (heart failure), Tnnt2^ΔK210/ΔK210^ CCR2^gfp/gfp^ (heart failure, CCR2 KO). CCR2^gfp^ knock-in mice allow visualization of CCR2+ cells by immunostaining or flow cytometry in the absence of CCR2 protein expression or signaling (Satpathy et al., 2013). Immunostaining at 8 weeks of age demonstrated increased abundance of both CCR2− macrophages and CCR2+ macrophages in Tnnt2^ΔK210/ΔK210^ hearts compared to controls. Deletion of CCR2 did not impact the abundance of CCR2− macrophages in non-failing or Tnnt2^ΔK210/ΔK210^ hearts. Conversely, CCR2 deletion led to significant reductions in the abundance of CCR2+ macrophages in non-failing and Tnnt2^ΔK210/ΔK210^ hearts (**Fig. S4A-B**). Flow cytometric analysis at 8 weeks of age confirmed selective reduction in CCR2+ macrophages in Tnnt2^ΔK210/ΔK210^ CCR2^gfp/gfp^ hearts compared to Tnnt2^ΔK210/ΔK210^ CCR2^gfp/+^ hearts (**Fig. S4C**). These data indicate that during this stage of chronic heart failure, CCR2− macrophages are maintained in the absence of monocyte input, whereas, CCR2+ macrophages require ongoing monocyte recruitment. Cell proliferation as assessed by Ki67 staining was exclusively increased in CCR2− macrophages in Tnnt2^ΔK210/ΔK210^ hearts compared to controls (**Fig. S4D-E**).

To delineate the developmental origin of cardiac macrophages in the chronically failing heart, we crossed Tnnt2^ΔK210/ΔK210^ mice to Flt3-Cre Rosa26-tdTomato mice. Flt3-Cre is selectively active in definitive hematopoietic stem cells, and thus labels monocytes and macrophages derived from definitive hematopoiesis (Boyer et al., 2011). This strategy has extensively been used to distinguish macrophages derived from extraembryonic hematopoiesis from macrophages derived from definitive hematopoiesis (Epelman et al., 2014a; Lavine et al., 2014; Leid et al., 2016). Consistent with previous reports, >90% of CCR2+ macrophages in control hearts were tdTomato+ at both 1 and 12 weeks of age. Conversely, <40% of CCR2− macrophages in control hearts were tdTomato+ highlighting the significant contribution of extraembryonic hematopoiesis to this macrophage subset. The frequency of tdTomato+ positivity did not significantly differ between control and Tnnt2^ΔK210/ΔK210^ hearts (**Fig. S4F-G**). Collectively, these findings indicate that in the context of chronic heart failure, CCR2− macrophages represent a mixed population of cells with contributions from extraembryonic and definitive hematopoiesis and are maintained by local proliferation in the absence of monocyte input. CCR2+ macrophages are exclusively derived from definitive hematopoiesis, long-lived, and maintained through gradual monocyte input.

### Tissue Resident CCR2− and CCR2+ Macrophages Represent Functionally Distinct Populations in the Chronically Failing Heart

To gain insights into functional differences between macrophage populations that reside within the chronically failing heart, we performed gene expression profiling using a high sensitivity microarray platform. RNA was harvested from the following cell populations isolated from Tnnt2^ΔK210/ΔK210^ Flt3-Cre Rosa26-tdTomato hearts using flow cytometry based cell sorting: CCR2+ macrophages (CCR2+MHC-II^high^tdTomato+), CCR2−MHC-II^low^tdTomato-macrophages, CCR2−MHC-II^high^tdTomato-macrophages, CCR2−MHC-II^low^tdTomato+ macrophages, CCR2−MHC-II^high^tdTomato+ macrophages, and Ly6C^high^ monocytes (Ly6C^high^MHC-II^low^CCR2+). Hierarchical clustering and principal component analysis revealed that the largest differences existed between CCR2− macrophages, CCR2+ macrophages, and Ly6C^high^ monocytes. CCR2+ macrophages clustered close to monocytes, which is consistent with ontological relationship between cell types. Importantly, each subset of CCR2− macrophages clustered together suggesting a high degree of similarity in gene expression amongst those populations (**Fig. 2A-B**).

**Figure 2.**
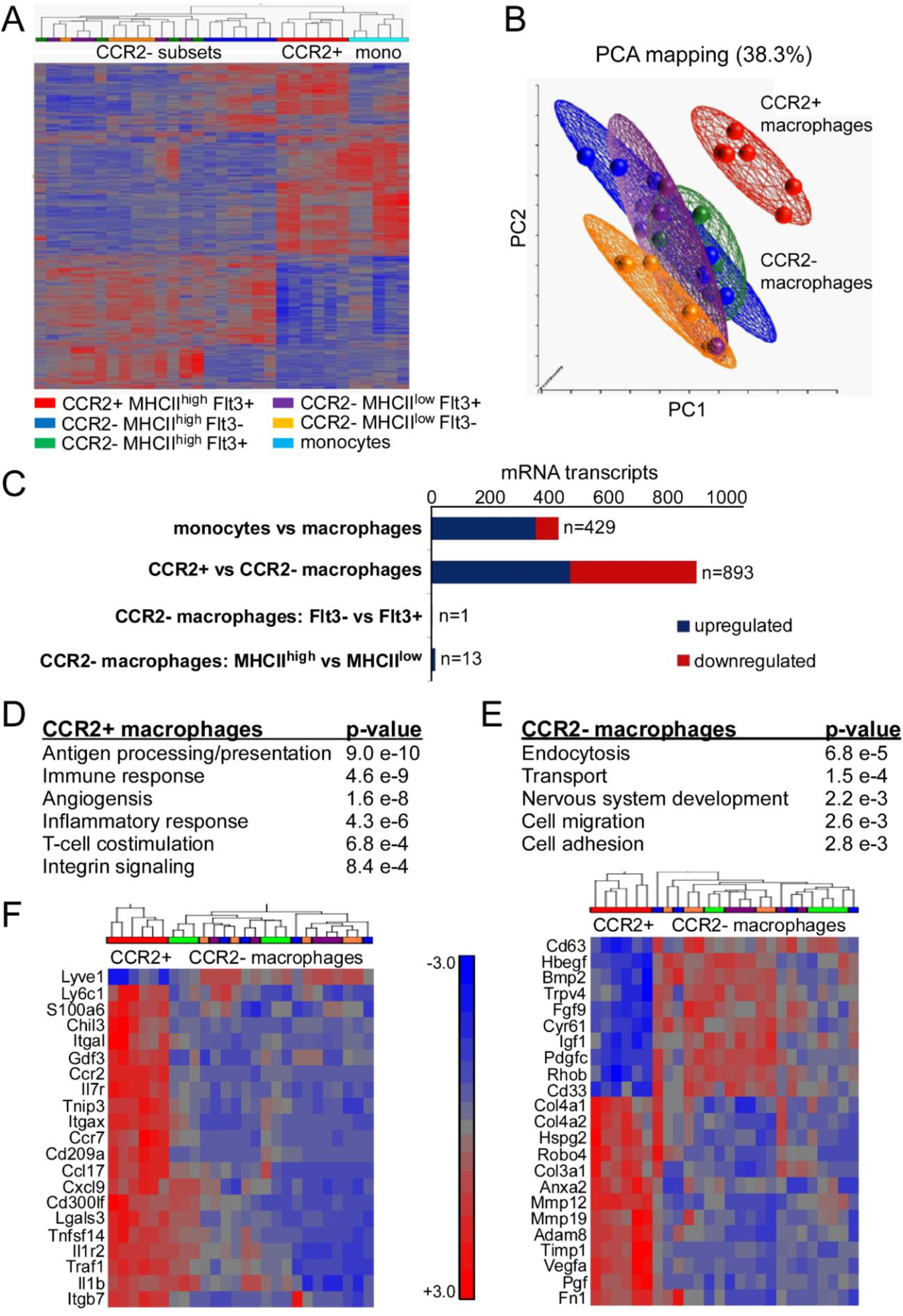
CCR2−and CCR2+ macrophages have distinct gene expression profiles in dilated cardiomyopathy. **A**, Hierarchical Clustering of CCR2+MHCII^high^ Flt3+, CCR2−MHCII^low^ Flt3-, CCR2−MHCII^high^ Flt3-, CCR2−MHCII^low^ Flt3+, CCR2−MHCII^high^ Flt3+, and CCR2+Ly6C^high^ monocytes FACS sorted from 8-week-old Tnnt2^ΔK210/ΔK210^ mice. n=5-7 per experimental group. **B**, Principal component analysis (PCA) highlighting that CCR2+MHCII^high^ Flt3+ macrophages cluster independently from each of the CCR2− macrophage populations. Each data point represents biologically independent samples. **C**, Bar graph showing the number of differentially expressed genes (FDR p<0.05, fold change>1.5) for each of the listed comparisons. Blue: upregulated in Tnnt2^ΔK210/ΔK210^ mice. Red: down regulated in Tnnt2^ΔK210/ΔK210^ mice. **D**, Pathways enriched in CCR2+ and CCR2− (all subgroups combined) macrophages. **F**, Heat maps listing individual genes differentially expressed in CCR2+ and CCR2− macrophages. Scale bar denotes fold change.

Differential gene expression analysis demonstrated that 893 genes were differentially expressed between CCR2− macrophages and CCR2+ macrophages and 429 genes were differentially expressed between monocytes and macrophages using a threshold value of 1.5-fold and FDR adjusted p-value<0.05, highlighting the marked divergence between CCR2− macrophages and CCR2+ macrophages. Few differences were observed between individual CCR2− macrophage populations. Comparisons between CCR2− macrophages derived from definitive and extra-embryonic hematopoiesis revealed a single differentially expressed gene. Only 13 genes were differentially expressed between CCR2−MHC-II^high^ and CCR2−MHC-II^low^ populations, many of which were MHC-II alleles (**Fig. 2C**). Pathway analysis of genes differentially expressed between CCR2− macrophages and CCR2+ macrophages demonstrated that CCR2+ macrophages expressed genes associated with antigen presentation, immune/inflammatory response, T-cell co-stimulation, integrin signaling, and angiogenesis (**Fig. 2D-E**). Examples of genes upregulated in CCR2+ macrophages included Il1β, Gdf3, Lgals3, Ccl17, Cxcl19, Itgax, Itgb7, Itgax, Traf1, Tnip3, Tnfsf14, Timp1, Mmp12, Mmp19, Vegfa, Pgf, Col4a1, Col3a1, and Fn1. In contrast, CCR2− macrophages showed enrichment of pathways associated with endocytosis/transport, nervous system development, cell adhesion, and migration in CCR2− macrophages. CCR2− macrophages expressed a paucity of inflammatory mediators and instead differentially expressed growth factors and genes associated with sensing mechanical stimuli including Igf1, Hbegf, Bmp2, Cyr61, Pdgfc, Fgf9, Trpv4, CD33, and Rhob (**Fig. 2F**).

### CCR2− macrophages are Required for Survival, Adaptive Tissue Remodeling, and Maintenance of Cardiac Output in the Chronically Failing Heart

We chose to focus on delineating the functions of CCR2− macrophages in the chronically failing heart, given their absolute abundance and unique gene expression signatures. We utilized CD169-DTR mice to selectively deplete CCR2− macrophages from the heart. We generated the following experimental groups: control, CD169-DTR, Tnnt2^ΔK210/ΔK210^, and Tnnt2^ΔK210/ΔK210^ CD169-DTR mice. Consistent with our previous findings (Bajpai et al., 2019), daily intraperitoneal administration of diphtheria toxin (DT) to CD169-DTR and Tnnt2^ΔK210/ΔK210^ CD169-DTR mice led to marked reduction in cardiac macrophage density and selective elimination of CCR2− macrophages (**Fig. 3A-D**). Neutrophil, monocyte, and CCR2+ macrophage abundance was not impacted by CCR2− macrophages depletion. CCR2− macrophage depletion did not increase CCR2+ macrophage chemokine or cytokine expression or result in alteration in serum chemistries or cytokines (**Fig. S5-6**).

**Figure 3.**
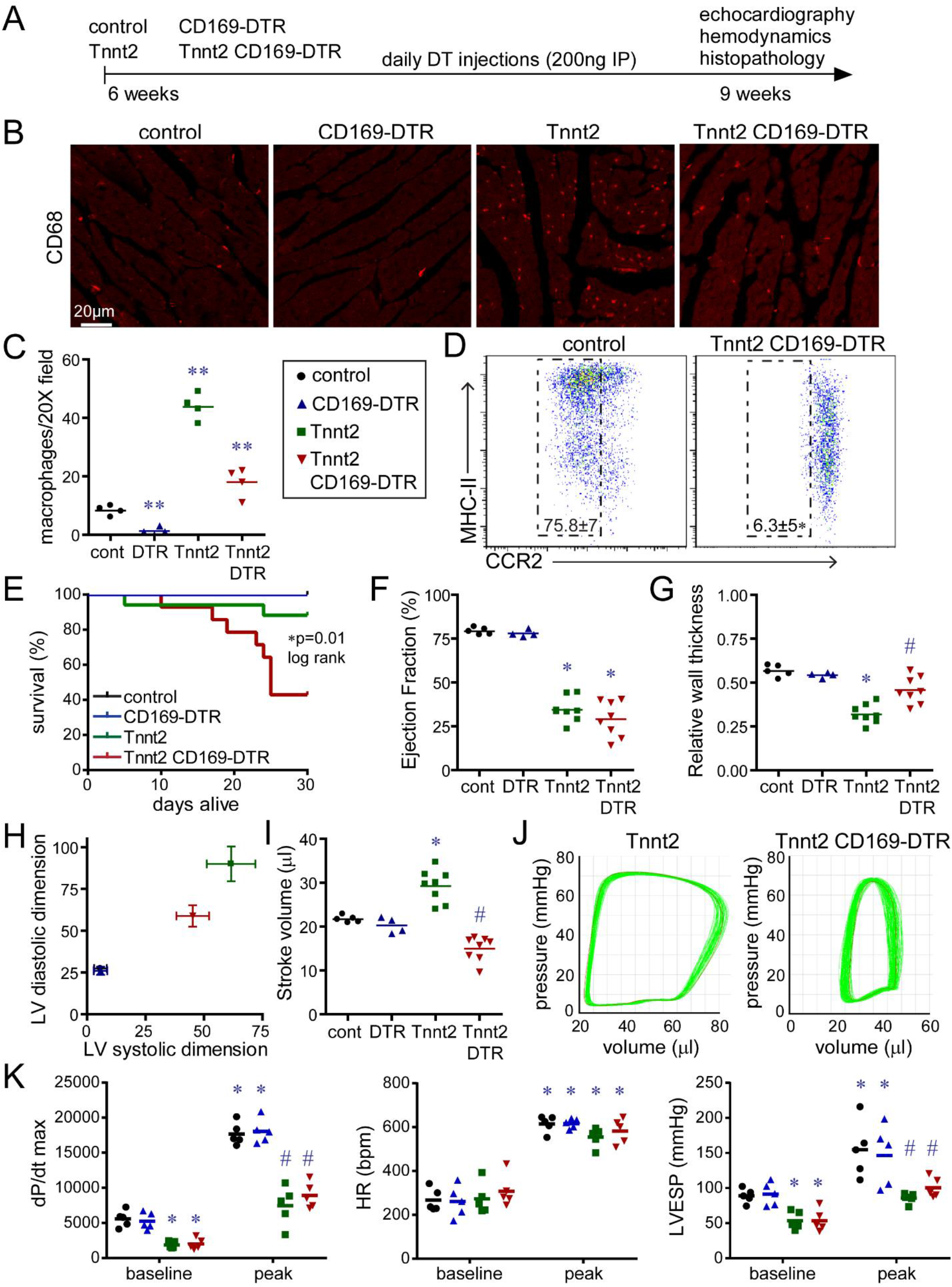
CCR2− macrophages influence survival and LV remodeling in dilated cardiomyopathy. **A**, Schematic outlining experimental groups, CCR2− macrophage depletion strategy, and analyzed endpoints. CCR2− macrophages were depleted by daily injection of diphtheria toxin (DT) into mice that expressed the diphtheria toxin receptor (DTR) under the control of the CD169 promoter. **B**, CD68 immunostaining of control, CD169-DTR, Tnnt2^ΔK210/ΔK210^, and Tnnt2^ΔK210/ΔK210^ CD169-DTR hearts after 3 weeks of DT treatment. N=4 per experimental group. **C**, Quantification of CD68 immunostaining. **D**, Flow cytometry plot of CD45^+^Ly6G^-^CD64^+^ macrophages showing specific depletion of CCR2− macrophages in Tnnt2^ΔK210/ΔK210^ CD169-DTR hearts compared to controls. N=4 per experimental group. **E**, Kaplan-Meier analysis demonstrating reduced survival of Tnnt2^ΔK210/ΔK210^ CD169-DTR compared to Tnnt2^ΔK210/ΔK210^ mice. No mortality was evident in control and CD169-DTR mice over the analyzed time period. n=12-15 per experimental group. **F-I**, Echocardiographic assessment of LV ejection fraction, relative wall thickness, LV volumes (μl), and stroke volume in control, CD169-DTR, Tnnt2^ΔK210/ΔK210^, and Tnnt2^ΔK210/ΔK210^ CD169-DTR mice 3 weeks after DT treatment. n=4-8 per experimental group. **J**, Representative pressure volume loops showing reduced stroke volume in Tnnt2^ΔK210/ΔK210^ CD169-DTR compared to Tnnt2^ΔK210/ΔK210^ mice. n=4 per experimental group. **K**, Invasive hemodynamic measurements of LV dP/dt max, heart rate (HR), and LV end systolic pressure (LVESP) at baseline and during peak infusion of dobutamine (64 ng/min) in control, CD169-DTR, Tnnt2^ΔK210/ΔK210^, and Tnnt2^ΔK210/ΔK210^ CD169-DTR mice 3 weeks after DT treatment. n=5 per experimental group. Each data point denotes independent animals. Error bars denote standard deviation. * denotes p<0.05 (ANOVA, Post-hoc Tukey) compared to controls. # denotes p<0.05 compared to Tnnt2^ΔK210/ΔK210^ mice.

To assess whether CCR2− macrophages influence survival, cardiac function, and LV remodeling in the context of chronic heart failure, we treated control, CD169-DTR, Tnnt2^ΔK210/ΔK210^, and Tnnt2^ΔK210/ΔK210^ CD169-DTR mice with DT beginning at 6 weeks of age. The primary endpoints included a Kaplan-Meier survival analysis and echocardiographic assessment of LV function and remodeling performed at 9 weeks of age (3-weeks of DT treatment). Kaplan-Meier analysis revealed reduced survival of Tnnt2^ΔK210/ΔK210^ CD169-DTR mice compared Tnnt2^ΔK210/ΔK210^ mice. No mortality was observed in control or CD169-DTR mice over the treatment period (**Fig. 3E**). Echocardiography demonstrated that similar reductions in LV ejection fraction between Tnnt2^ΔK210/ΔK210^ mice and Tnnt2^ΔK210/ΔK210^ CD169-DTR mice compared to controls **(Fig. 3F**). Tnnt2^ΔK210/ΔK210^ mice displayed significantly greater LV remodeling (LV dilation and reduced relative LV wall thickness) compared to Tnnt2^ΔK210/ΔK210^ CD169-DTR mice (**Fig. 3G-H**). Tnnt2^ΔK210/ΔK210^ mice displayed increased LV stroke volumes compared to controls, whereas Tnnt2^ΔK210/ΔK210^ CD169-DTR mice had lower LV stroke volumes compared to both control and Tnnt2^ΔK210/ΔK210^ mice. (**Fig. 3I**). No differences were observed between control and CD169-DTR mice for all echocardiographic variables examined. Simultaneous measurement of LV pressure and volume confirmed reductions in LV end diastolic volume and stroke volume (measures of dilation) in Tnnt2^ΔK210/ΔK210^ CD169-DTR compared to Tnnt2^ΔK210/ΔK210^ mice (**Fig. 3J**). LV catheterization and dobutamine infusion revealed reduced LV end systolic pressure, myocardial contractility, and contractile reserve in Tnnt2^ΔK210/ΔK210^ mice compared to controls. CCR2− macrophage depletion did not impact measurements of myocardial contractility in either wild type or Tnnt2^ΔK210/ΔK210^ backgrounds (**Fig. 3K, Fig. S7**). Collectively, these findings indicate that CCR2− macrophage depletion blunts LV chamber remodeling in Tnnt2^ΔK210/ΔK210^ mice without impacting myocardial contractility.

### CCR2− macrophages are Required for Myocardial Tissue Remodeling and Coronary Angiogenesis

We performed histological analysis following 3 weeks of DT treatment to examine whether structural differences within the myocardium explain reductions in LV dilation and remodeling in Tnnt2^ΔK210/ΔK210^ CD169-DTR hearts. Compared to controls, Tnnt2^ΔK210/ΔK210^ hearts displayed evidence of myocardial reorganization consisting of circumferential enlargement of the LV, loss of trabecular myocardial tissue, and expansion of compact myocardial tissue. Tnnt2^ΔK210/ΔK210^ CD169-DTR hearts displayed reduced circumferential LV enlargement, persistent trabecular myocardial tissue, and failed to expand the compact myocardium. CD169-DTR mice displayed no significant changes compared to controls (**Fig. 4A-C**). To verify these structural alterations by a second method, we performed X-Ray microscopy (XRM), a variant of micro-computed tomography (μCT). This technique provides full volume datasets enabling three-dimensional reconstruction of cardiac anatomy and virtual histology analysis. Surface projection and virtual histology images of the LV chamber revealed the presence of smooth appearing walls and precise alignment of muscle fibers in Tnnt2^ΔK210/ΔK210^ hearts. In contrast, the LV walls of Tnnt2^ΔK210/ΔK210^ CD169-DTR hearts displayed a more ribbon-like appearance and impaired muscle fiber alignment (**Fig. 4D**). Collectively, these findings indicate that CCR2− macrophages influence LV remodeling through alterations in myocardial tissue organization.

**Figure 4.**
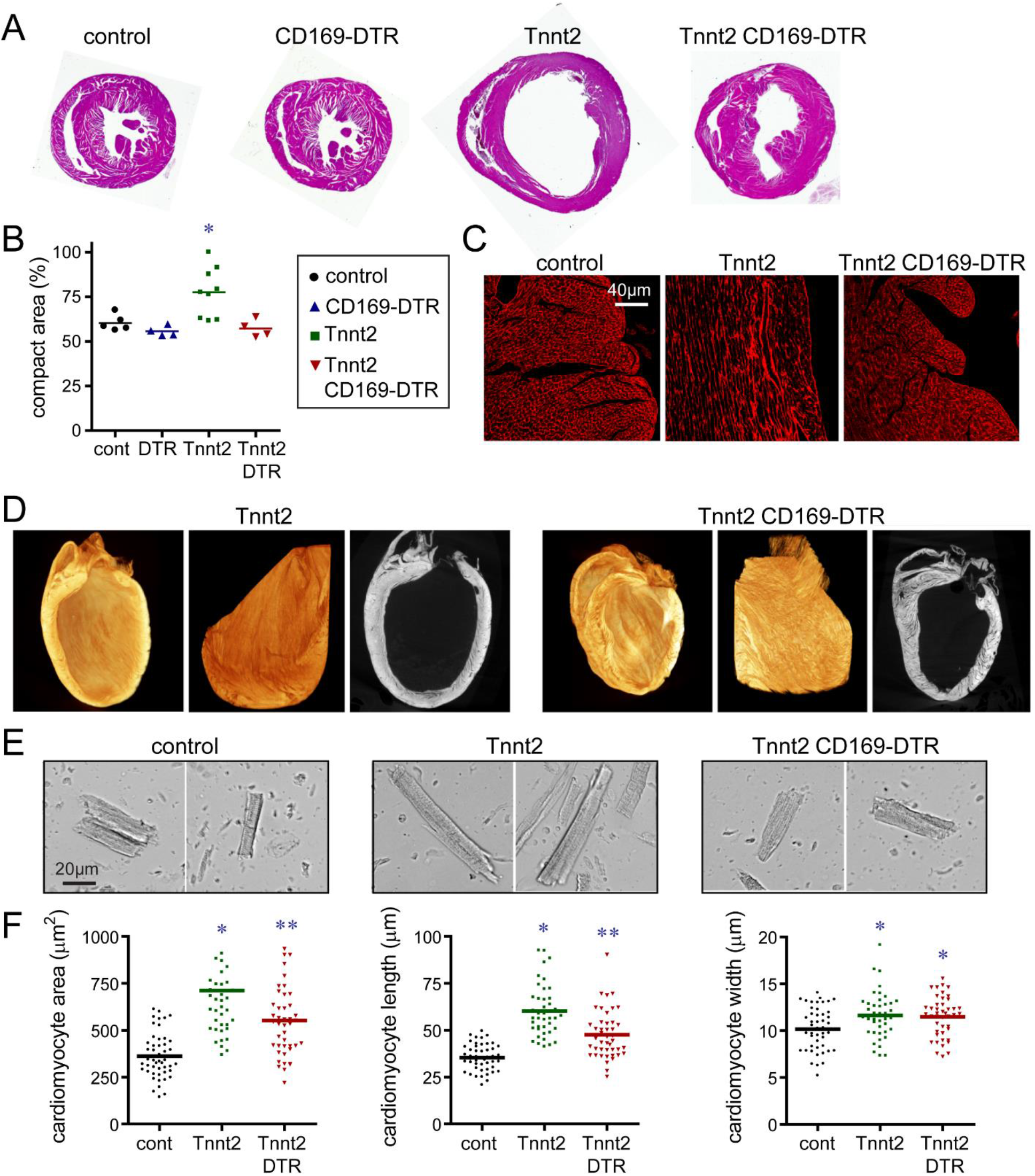
CCR2− macrophages orchestrate myocardial tissue adaptations in dilated cardiomyopathy. **A**, Low magnification H&E images of control, CD169-DTR, Ţnnt2^ΔK210/ΔK210^, and Tnnt2^ΔK210/ΔK210^ CD169-DTR hearts (LV in cross-section) after 3 weeks of DT treatment. n=4-9 per experimental group. **B**, Quantification of the ratio of compact to trabecular myocardium in control, CD169-DTR, Tnnt2^ΔK210/ΔK210^, and Tnnt2^ΔK210/ΔK210^ CD169-DTR hearts. Each data point represents an individual animal. **C**, Wheat germ agglutinin (WGA, red) staining showing alterations in the alignment of trabecular cardiomyocytes in Tnnt2^ΔK210/ΔK210^ compared to control hearts. The alignment of trabecular cardiomyocytes inTnnt2^ΔK210/ΔK210^ CD169-DTR hearts is indistinguishable from controls. n=4-9 per experimental group. **D**, Reconstructed X-ray microscopy images and virtual histology comparing the myocardial architecture of Tnnt2^ΔK210/ΔK210^ and Tnnt2^ΔK210/ΔK210^ CD169-DTR hearts following 3 weeks of DT injection. n=4 per experimental group. The endocardial surface of Tnnt2^ΔK210/ΔK210^ hearts is smooth and compacted whereas the endocardial surface of Tnnt2^ΔK210/ΔK210^ CD169-DTR hearts has a rougher and meshwork-like appearance. **E**, Images of cardiomyocytes digested from control, Tnnt2^ΔK210/ΔK210^, and Tnnt2^ΔK210/ΔK210^ CD169-DTR hearts. Hearts were relaxed in potassium prior to fixation. **F**, Quantification of cardiomyocyte area, length, and width. Each data point represents individual cardiomyocytes analyzed from 4 independent animals per experimental group. * denotes p<0.05 (ANOVA, Post-hoc Tukey) compared to controls. ** denotes p<0.05 compared to control and Tnnt2^ΔK210/ΔK210^ mice.

To examine whether CCR2− macrophages also affect cardiomyocyte size, we performed a morphometric analysis of cardiomyocytes isolated from control, Tnnt2^ΔK210/ΔK210^, and Tnnt2^ΔK210/ΔK210^ CD169-DTR hearts. Tnnt2^ΔK210/ΔK210^ and Tnnt2^ΔK210/ΔK210^ CD169-DTR cardiomyocytes demonstrated increased 2-dimensional area compared to cardiomyocytes isolated from control hearts. However, the extent of cell enlargement was greater in Tnnt2^ΔK210/ΔK210^ cardiomyocytes compared to Tnnt2^ΔK210/ΔK210^ CD169-DTR cardiomyocytes. Tnnt2^ΔK210/ΔK210^ cardiomyocytes displayed increases in both cell width and length compared to control hearts. Interesting, Tnnt2^ΔK210/ΔK210^ CD169-DTR cardiomyocytes displayed similar increases in cell width, but less extensive cell lengthening (**Fig. 4E-F**). These data suggest that modulation of cardiomyocyte length might also be involved in CCR2− macrophage dependent LV remodeling.

To evaluate whether CCR2− macrophages influence adverse/pathological LV remodeling, we examined cardiomyocyte cross-sectional area, interstitial fibrosis, and mRNA expression of established marker genes. Examination of cardiomyocyte cross-sectional area *in situ* using wheat germ agglutinin stained sections demonstrated increased cardiomyocyte area in both Tnnt2^ΔK210/ΔK210^ and Tnnt2^ΔK210/ΔK210^ CD169-DTR hearts compared to controls. Of note, minimal evidence of interstitial fibrosis was evident in this model at the stages examined (**Fig. S8A-D**). Nppa, Nppb, and Myh7 mRNA expression was increased to a similar degree in Tnnt2^ΔK210/ΔK210^ and Tnnt2^ΔK210/ΔK210^ CD169-DTR hearts compared to controls (**Fig. S8E-G**). Previous studies have implicated matrix metalloproteinase (MMP) activity in LV dilation and remodeling in the context of myocardial infarction and injury models (Ducharme et al., 2000; Heymans et al., 1999). We observed negligible MMP9 activity in the hearts of control and Tnnt2^ΔK210/ΔK210^ mice. Robust MMP9 activity was present in a mouse model of DT-mediated cardiomyocyte ablation that mimics myocardial infarction (**Fig. S8H**).

Given previous findings that CCR2− macrophages regulate coronary angiogenesis in the embryonic and neonatal heart (Lavine et al., 2014; Lavine et al., 2013; Leid et al., 2016), we evaluated whether CCR2− macrophages also modulate coronary angiogenesis in the context of chronic heart failure. Visualization of the coronary arterial vasculature using Microfil casting demonstrated marked increases in epicardial coronary arterial vasculature in Tnnt2^ΔK210/ΔK210^ hearts compared to controls. Strikingly, we observed expansion of the epicardial coronary arterial vasculature was markedly attenuated in Tnnt2^ΔK210/ΔK210^ CD169-DTR hearts (**Fig. 5A**). Measurement of coronary microvascular density similarly revealed robust increases in Tnnt2^ΔK210/ΔK210^ hearts compared to controls and Tnnt2^ΔK210/ΔK210^ CD169-DTR hearts (**Fig. 5B-C**). To explore potential mechanisms by which CCR2− macrophages promote coronary angiogenesis, we measured the expression pro-angiogenic growth factors expressed in CCR2− macrophages. Consistent with such a mechanism, Igf1, Pdgfc, Cyr61, and Hbegf mRNA expression was increased in CCR2− macrophages from Tnnt2^ΔK210/ΔK210^ hearts compared to controls (**Fig. 5D**). Immunostaining analysis further revealed that macrophage IGF1 and CYR61 expression was increased in Tnnt2^ΔK210/ΔK210^ hearts compared to controls. Increased macrophage IGF1 and CYR61 expression was not evident in Tnnt2^ΔK210/ΔK210^ CD169-DTR hearts, presumably due to the absence of CCR2− macrophages (**Fig. 5E-F**). These observations indicate that CCR2− macrophages promote coronary angiogenesis in the context of chronic heart failure.

**Figure 5.**
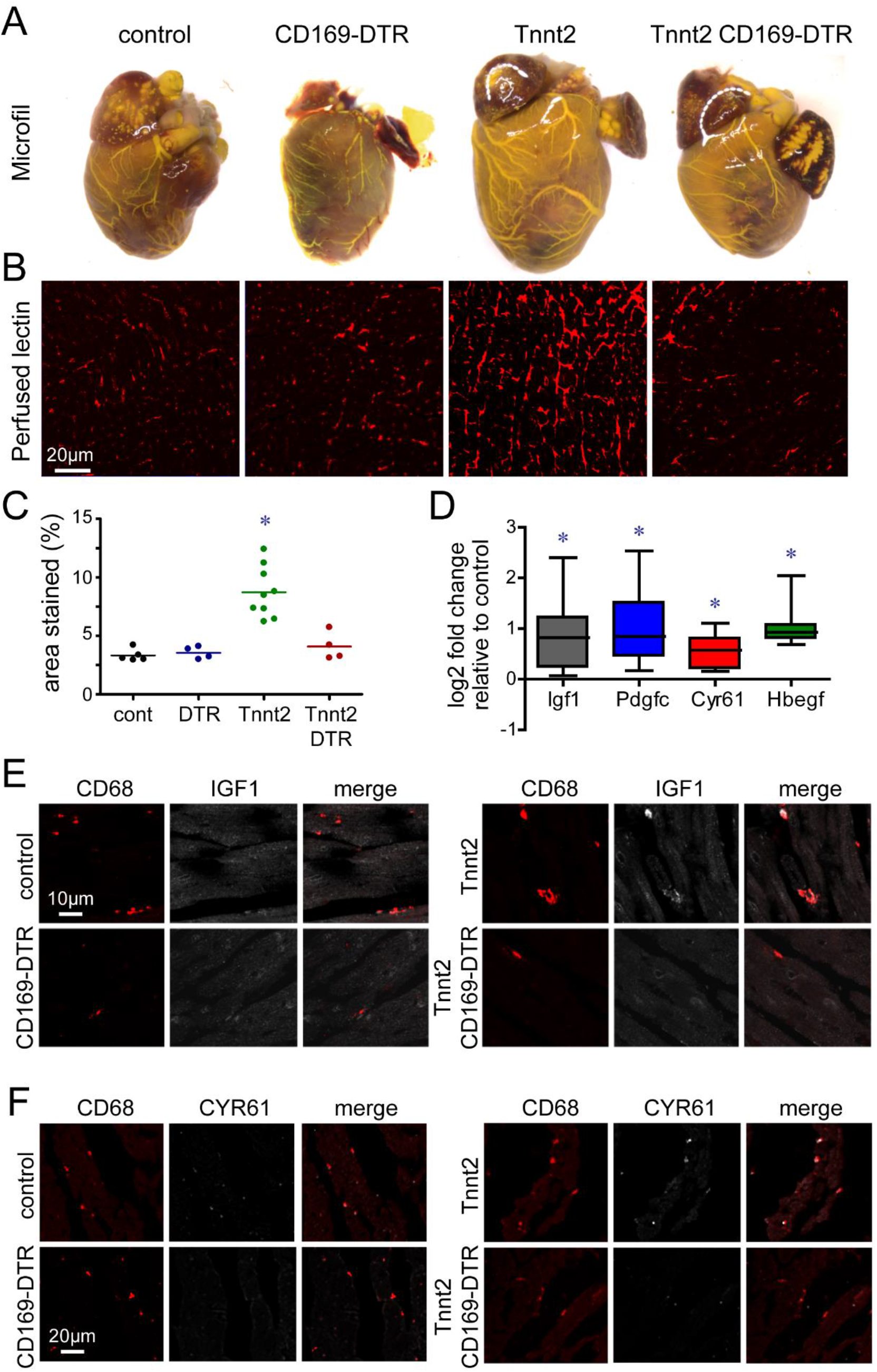
CCR2− macrophages are essential for coronary angiogenesis in dilated cardiomyopathy. **A**, Microfil vascular casting of control, CD169-DTR, Tnnt2^ΔK210/ΔK210^, and Tnnt2^ΔK210/ΔK210^ CD169-DTR hearts after 3 weeks of DT treatment. Tnnt2^ΔK210/ΔK210^ hearts display marked expansion in the coronary macrovasculature that is diminished in Tnnt2^ΔK210/ΔK210^ CD169-DTR hearts. n=4 per experimental group. **B**, Microvascular imaging (perfused lectin) of control, CD169-DTR, Tnnt2^ΔK210/ΔK210^, and Tnnt2^ΔK210/ΔK210^ CD169-DTR hearts after 3 weeks of DT treatment. Tnnt2^ΔK210/ΔK210^ hearts display marked expansion in the coronary microvasculature that is diminished in Tnnt2^ΔK210/ΔK210^ CD169-DTR hearts. n=4-9 per experimental group. **C**, Quantification of the coronary microvascular in control, CD169-DTR, Tnnt2^ΔK210/ΔK210^, and Tnnt2^ΔK210/ΔK210^ CD169-DTR hearts. Each data point represents an individual animal. * denotes p<0.05 (ANOVA, Post-hoc Tukey) compared to controls. **D**, CCR2− macrophages from Tnnt2^ΔK210/ΔK210^ hearts display increased Igf1, Pdgfc, Cyr61, and Hbegf mRNA expression compared to CCR2− macrophages isolated from control mice. n=5 per experimental group. * denotes p<0.05 (Mann-Whitney test) compared to controls. **E-F**, Immunostaining of control, CD169-DTR, Tnnt2^ΔK210/ΔK210^, and Tnnt2^ΔK210/ΔK210^ CD169-DTR hearts after 3 weeks of DT treatment showing that macrophages within the LV myocardium of Tnnt2^ΔK210/ΔK210^ mice display increased expression of IGF1 (E, white) and CYR61 (F, white). Red: CD68. n=5 per experimental group.

Previous work has suggested that cardiac macrophages have the potential to influence propagation of electrical signals through the atrioventricular node (Hulsmans et al., 2017). To assess whether alterations in electoral conduction occurred following depletion of CCR2− macrophages from control and Tnnt2^ΔK210/ΔK210^ mice, we analyzed surface electrocardiograms (ECGs) obtained from anesthetized (isoflurane) control, CD169-DTR, Tnnt2^ΔK210/ΔK210^, and Tnnt2^ΔK210/ΔK210^ CD169-DTR mice treated with DT for 3 weeks. We did not observe significant differences in RR (heart rate), PR (atrioventricular node conduction), or QRS (intraventricular conduction) intervals between experimental groups. Tnnt2^ΔK210/ΔK210^ and Tnnt2^ΔK210/ΔK210^ CD169-DTR mice displayed prolongation of the QT (ventricular repolarization) interval compared to control and CD169-DTR mice. No significant differences were observed between Tnnt2^ΔK210/ΔK210^ and Tnnt2^ΔK210/ΔK210^ CD169-DTR mice for any examined parameter (**Fig. S9**). While these results indicate that defects in electrical propagation are unlikely to account for increased mortality observed in Tnnt2^ΔK210/ΔK210^ CD169-DTR mice, they do not rule out the possibility that CCR2− macrophages contribute to optimal cardiac conduction. Intriguingly, we found that both CCR2−and CCR2+ macrophages are located within the AV node potentially accounting for the lack of an overt electrical phenotype (**Fig. S10**).

A recent study suggested that tissue resident cardiac macrophages regulate myocardial metabolism and function through effects on mitochondrial homeostasis (Nicolas-Avila et al., 2020). To examine whether this might contribute to the phenotype of Tnnt2^ΔK210/ΔK210^ CD169-DTR mice, we isolated mitochondria from control, CD169-DTR, Tnnt2^ΔK210/ΔK210^, and Tnnt2^ΔK210/ΔK210^ CD169-DTR hearts following 3 weeks of DT treatment. We did not detect any differences in mitochondrial respiration across experimental groups (**Fig. S11)**, indicating that alterations in mitochondrial function are unlikely responsible for the cardiac phenotype of Tnnt2^ΔK210/ΔK210^ CD169-DTR mice.

### CCR2− macrophages Physically Interact with Neighboring Cardiomyocytes

To gain insights into how resident cardiac macrophages might influence myocardial remodeling and angiogenesis, we examined CCR2−and CCR2+ macrophage localization and structure in control and Tnnt2^ΔK210/ΔK210^ hearts using three-dimensional confocal microscopy. Under baseline and heart failure conditions, CCR2− macrophages were observed within close proximity to cardiomyocytes and appeared to extend processes that contacted adjacent cardiomyocytes. CCR2+ macrophages were also found within the myocardium and extended processes within the interstitial space (**Fig. 6A-B**). Electron microscopy of Tnnt2^ΔK210/ΔK210^ CCR2^gfp/+^ hearts stained with anti-CD68 and anti-GFP antibodies confirmed that CCR2− macrophages were in close apposition to neighboring cardiomyocytes and revealed the presence of physical contacts between these two cell types (**Fig. 6C**). CCR2+ macrophages were also found within the myocardium and extended processes into the interstitial space but did not directly contact cardiomyocytes (**Fig. 6D**). The projection length of CCR2− macrophages was greater in Tnnt2^ΔK210/ΔK210^ hearts compared to controls. The projection length of CCR2+ macrophages was indistinguishable between control and Tnnt2^ΔK210/ΔK210^ CCR2^gfp/+^ hearts (**Fig. 6E**).

**Figure 6.**
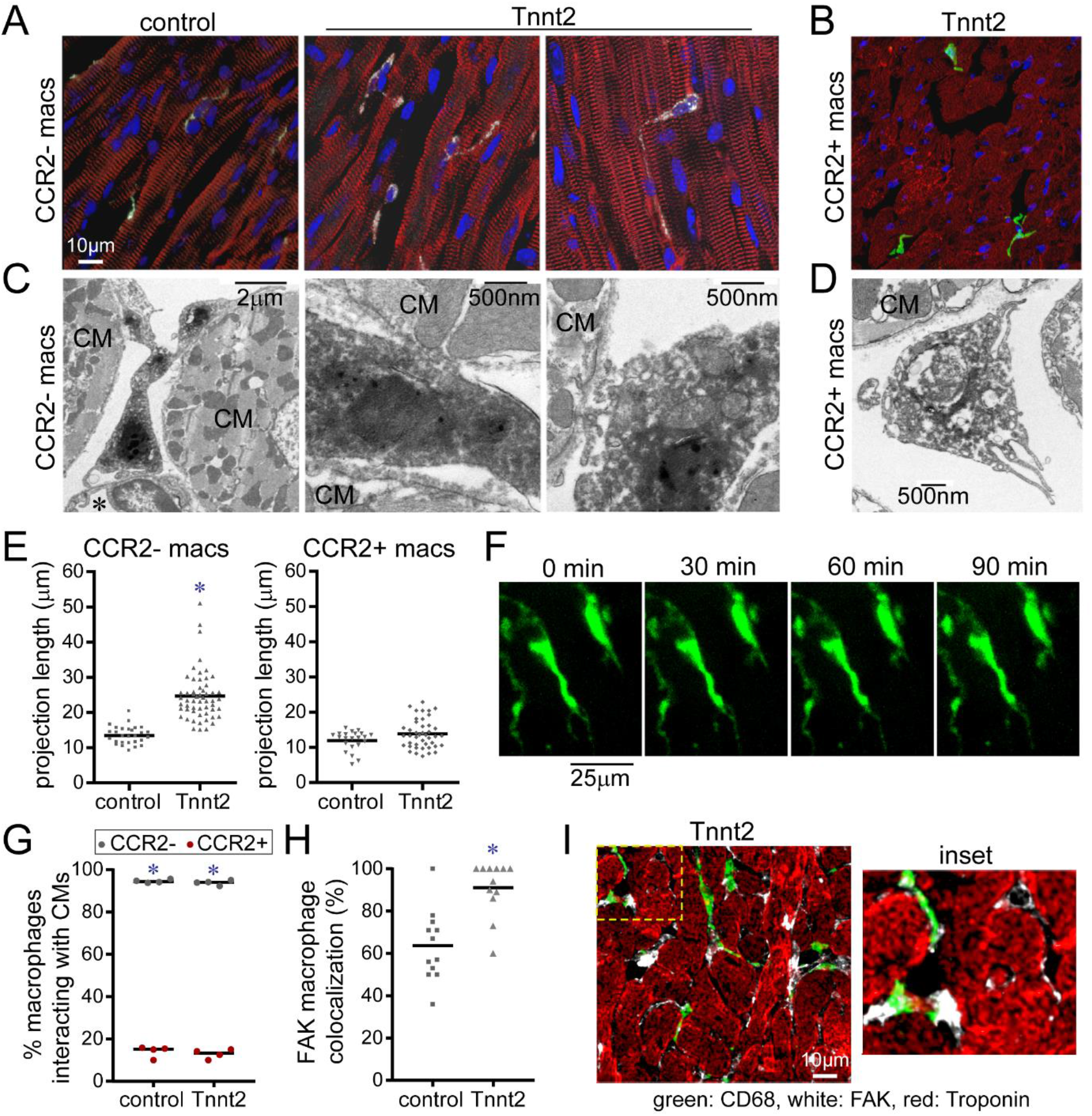
CCR2− macrophages interact with neighboring cardiomyocytes through focal adhesion complexes. **A-B**, Compressed Z-stack images showing distinct morphologies of CCR2−(A) and CCR2+ (B) cardiac macrophages in the LV myocardium of a 8-week-old control CCR2^GFP/+^ and Tnnt2^ΔK210/ΔK210^ CCR2^GFP/+^ hearts. CCR2−cardiac macrophage projections are closely associated with neighboring cardiomyocytes whereas CCR2+ macrophage projections are shorter and remain localized to the interstitial space. n=4-6 per experimental group. CD68: white, GFP: green, α-actinin, red, DAPI: blue. **C-D**, Electron microscopy of CCR2− (C) and CCR2+ (D) cardiac macrophages in Tnnt2^ΔK210/ΔK210^ CCR2^GFP/+^ hearts. CCR2− macrophage are found adjacent to endothelial cells (*) and make contact with cardiomyocytes (CM). CCR2+ macrophages remain within the interstitial space between cardiomyocytes. n=4 per experimental group. **E**, Measurement of projection length in CCR2−and CCR2+ macrophages found within the myocardium of 8-week-old control CCR2^GFP/+^ and Tnnt2^ΔK210/ΔK210^ CCR2^GFP/+^ hearts. * denotes p<0.05 (Mann-Whitney test) compared to controls. **F**, 2-photon microscopy of live CX3CR1^GFP/+^ CCR2^RFP/+^ papillary muscle cell preparations (n=4) showing that projections emanating from CCR2− macrophages (green) remain stable over 90 minutes. **G**, Quantification of the percent of CCR2− (black data points) and CCR2+ (red data points) macrophages interacting with cardiomyocytes in control and Tnnt2^ΔK210/ΔK210^ hearts. Each data point represents an individual animal (n=4). * denotes p<0.05 (Mann-Whitney test) comparing CCR2−to CCR2+ macrophages. **H-I**, Immunostaining for CD68 (green), FAK (white), and troponin (red) reveals evidence of focal adhesion complexes at sites of macrophage-cardiomyocyte interaction. Each data point represents an individual animal (n=12 per experimental group). * denotes p<0.05 (Mann-Whitney test) compared to controls.

To characterize the temporal dynamics of how CCR2− macrophages interact with adjacent cardiomyocytes *in situ*, we performed two-photon microscopy on isolated mouse papillary muscle preparations. Papillary muscles were isolated from Cx_3_cr1^GFP/+^ Ccr2^RFP/+^ mice and imaged for 1-2 hours in a temperature-controlled imaging chamber containing oxygenated media. Cellular processes were observed that extended from CCR2− macrophages and formed stable contacts with neighboring cardiomyocytes. These processes did not further extend or retract over the 90-minute time course of imaging (**Fig. 6F, supplemental movie**). Predominately CCR2− macrophages interacted with cardiomyocytes (**Fig. 6G**).

We did not observe electron densities indicative of desmosomes, adherens junctions, or tight junctions between CCR2− macrophages and cardiomyocytes. Immunostaining for α-cadherin (adherens conjunctions), desmoplakin (desmosomes), claudin (tight conjunctions), and Cx43 (gap junctions) revealed infrequent co-localization of these markers at sites of macrophage-cardiomyocyte interactions. Instead, we found that FAK and Paxillin (markers of focal adhesion complexes) were frequently present between CCR2− macrophages and cardiomyocytes (**Fig. 6H-I, Fig. S12**).

We then utilized an *in vitro* system to delineate whether focal adhesion complexes were responsible for macrophage-cardiomyocyte interactions. We found that HL-1 cardiomyocytes and bone marrow-derived macrophages formed spontaneous interactions when co-cultured. Electron microscopy revealed evidence of physical interaction between HL1-cells and cardiomyocytes. Electron densities consistent with desmosomes, adherens, tight, or gap junctions were not evident. Immunostaining showed presence of Paxillin staining at sites of interactions between HL-1 cells and bone marrow-derived macrophages (**Fig. S13**).

As previous studies have established that β-integrins are essential for formation of focal adhesion complexes (Parsons et al., 2010), we focused our attention on inhibiting β-integrin binding. Addition of RDG peptides or antibodies that blocked either β1-integrin or β2-integrin binding was sufficient to disrupt interactions between HL-1 cells and bone marrow-derived macrophages. Application of β1-integrin neutralizing antibodies also resulted in loss of macrophage Paxillin staining at sites of cardiomyocyte-macrophage interaction (**Fig. S14**). Collectively, these findings indicate that CCR2− macrophages physically interact with cardiomyocytes and form focal adhesion complexes at sites of cell-cell contacts.

### TRPV4 Regulates Growth Factor Expression in Macrophages

Based on their morphology and physical interaction with neighboring cardiomyocytes, we considered the possibility that CCR2− macrophages may be activated by mechanical cues in the context of heart failure. Specifically, we proposed that elevated LV chamber pressure and resultant increased LV myocardial wall stress may be sensed by CCR2− macrophages through their interactions with neighboring cardiomyocytes. To explore this concept, we assayed the expression of known mechanoresponsive factors and found that TRPV4 mRNA was abundantly expressed in CCR2− macrophages (**Fig. 7A, Fig. S15**). Ratiometric calcium assays demonstrated that the TRPV4 channel was active in cardiac macrophages. Treatment of CCR2−and CCR2+ macrophages isolated from the heart by flow cytometry with a highly specific TRPV channel activator (GSK101) or TRPV4 channel inhibitor (GSK219) confirmed functional expression of TRPV4 protein in both CCR2−and CCR2+ macrophages within the ventricular myocardium (**Fig. 7B, Fig. S15**). Immunostaining of TRPV4-GFP BAC transgenic mice provided further evidence the TRPV4 was predominately expressed in macrophages located within the LV myocardium (**Fig. 7C**). Flow cytometry demonstrated expression of TRPV4 in cardiac macrophages and neutrophils (**Fig. 7D, Fig. S16**). We additionally detected TRPV4 activity in cardiac macrophages *in situ* using CX3CR1-ertCre Rosa26-GCaMP6s/tdtomato reporter mice (Madisen et al., 2015), which allows visualization of macrophage cytoplasmic calcium by restricting GCaMP6s expression to macrophages. 2-photon imaging of papillary muscles isolated from CX3CR1-ertCre Rosa26-GCaMP6/tdTomato hearts placed under axial tension revealed GCaMP signal in cardiac macrophages. Application of a TRPV4 inhibitor (GSK219) suppressed GCaMP signal, indicating that TRPV4 channel activity is responsible for the observed rise in macrophage cytoplasmic calcium (**Fig. 7E-F, Fig. S17**).

**Figure 7.**
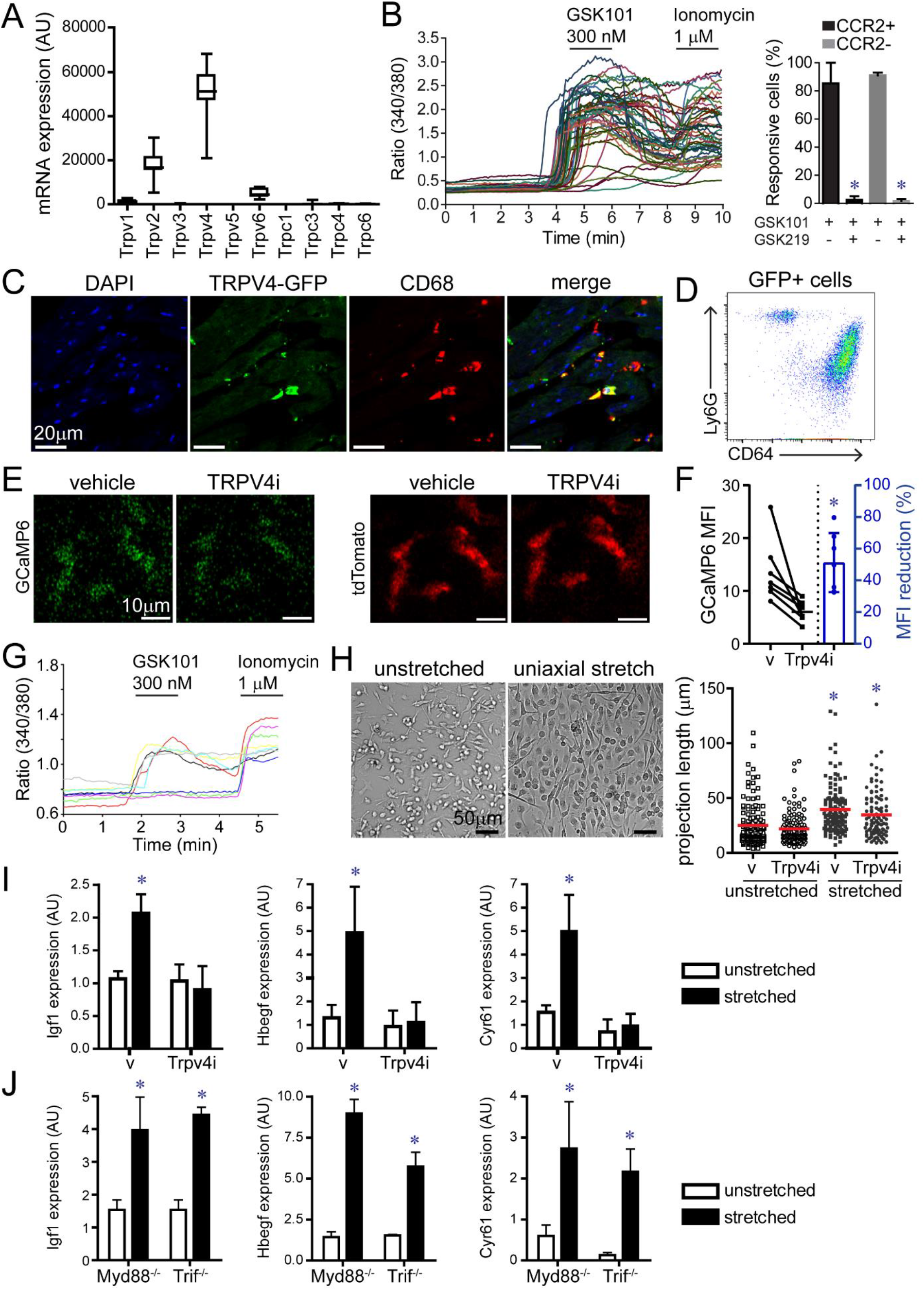
The mechanoresponsive TRPV4 channel regulates growth factor expression in macrophages. **A**, mRNA expression of TRP channels in CCR2− macrophages showing robust expression of Trpv4. Data generated from n=20 samples. **B**, Ratiometric calcium assay demonstrating that cardiac macrophages have active TRVP4 channels. GSK101: TRPV4 agonist, GSK219: TRPV4 antagonist. * denotes p<0.05 comparing GSK101 treated cells with GSK101 and GSK219 treated cells. **C**, Immunostaining of TRPV4-GFP BAC transgenic mice showing GFP (green) expression in CD68+ macrophages (red) within the LV myocardium. **D**, Flow cytometry of cardiac CD45+GFP+ leukocytes isolated from TRPV4-GFP heart revealing that macrophages and neutrophils express TRPV4. Repetitive images from n=4 mice. **E**, 2-photon imaging of GFP (green) and tdTomato (red) in papillary muscle preparations harvested from CX3CR1-ertCre; Rosa26-GCaMP6/tdTomato mice treated with either vehicle or the TRPV4 inhibitor GSK219 (TRPV4i). **F**, Quantification of GCaMP6 signal. Each data point represent mean data from an individual experiment (n=6). * denotes p<0.05 compared to vehicle. **G**, Ratiometric calcium assay showing that bone marrow derived macrophages express active TRPV4 channels. GSK101: TRPV4 agonist, Ionomycin: calcium ionophore. **H**, Cyclic uniaxial stretch (1 Hz, 10% deformation, 24 hours) promotes elongation of bone marrow derived macrophages independent of TRPV4 channel activity. n=4 independent experiments. **I**, Quantitative RT-PCR assays demonstrating that cyclic uniaxial stretch promotes increased Igf1, Hbegf, and Cyr61 mRNA expression in bone marrow-derived macrophages. Upregulation of Igf1, Hbegf, and Cyr61 mRNA expression by uniaxial cyclic stretch is dependent on TRPV4 channel activity. n=4 independent experiments. **J**, Quantitative RT-PCR assays demonstrating that Igf1, Hbegf, and Cyr61 mRNA expression by uniaxial cyclic stretch is independent of MYD88 and TRIF signaling pathways. n=4 independent experiments. * denotes p<0.05 compared to vehicle treated unstretched cells (ANOVA post-hoc Tukey) (F-H). Error bars denote standard deviation (G-H).

To examine the possibility that TRPV4 mediates the activation of macrophages by mechanical cues, we first utilized bone marrow-derived macrophages. Ratiometric calcium assays confirmed that bone marrow-derived macrophages express functional TRPV4 channels (**Fig. 7G**). Immunostaining of bone marrow-derived macrophages co-cultured with HL1 cardiomyocytes revealed expression of TRPV4 expression at sites of macrophage and cardiomyocyte interaction (**Fig. S18**). We then subjected bone marrow-derived macrophages to cyclic mechanical stretch. Cells were cultured on silicone membranes coated with collagen and fibronectin. Membranes were then stretched (10% deformation, 1 Hz) for 24 hours in the presence of vehicle control or GSK219 (TRPV inhibitor). Following 24 hours of cyclic uniaxial stretch, both vehicle and TRPV4 inhibitor treated macrophages elongated and aligned along the axis of membrane deformation (**Fig. 7H**). Quantitative RT-PCR assays revealed that bone marrow-derived macrophages expressed increased levels of Igf1, Hbegf, and Cyr61 mRNA in response to mechanical stretch. Application of the TRPV4 channel inhibitor blocked this response (**Fig. 7I**).

To determine whether canonical pathways involved in macrophage activation affected the ability of mechanical stretch to induce macrophage growth factor expression, we subjected control, Myd88^-/-^ and Trif^-/-^ bone marrow derived-macrophages to uniaxial cyclic stretch. Quantitative RT-PCR assays demonstrated that deletion of MYD88 or TRIF had no impact on the expression of Igf1, Hbegf, or Cyr61 (Fig. 7**J**). Further, activators of MYD88 and TRIF signaling (LPS, PolyIC) were unable to increase the expression of Igf1, Hbegf, or Cyr61 and mechanical stretch did not induce the expression of inflammatory cytokines (**Fig. S19**). Collectively, these observations indicate that mechanical stretch promotes growth factor expression from macrophages through a TRPV4 dependent mechanism that is independent of MYD88 and TRIF signaling.

### TRPV4 regulates IGF1 expression in CCR2− macrophages and is required for coronary angiogenesis

To assess the functional relevance of TRPV4 *in vivo*, we treated control and Tnnt2^ΔK210/ΔK210^ mice with either vehicle control, TRPV4 inhibitor, or TRPV4 agonist beginning at 6 weeks of age. Immunostaining for CD68 and IGF1 after 2 days of treatment revealed that TRPV4 activity modulates cardiac macrophage IGF1 expression. Compared to controls, cardiac macrophages in Tnnt2^ΔK210/ΔK210^ hearts expressed IGF1 at higher frequency and increased mean levels. Treatment of Tnnt2^ΔK210/ΔK210^ mice with the TRPV4 inhibitor was sufficient to reduce IGF1 expression (frequency and mean levels) in cardiac macrophages. Conversely, mice treated with the TRPV4 agonist displayed increased IGF1 expression in cardiac macrophages (**Fig. 8A-B, Fig. S20**). These data indicate that TRPV4 regulates cardiac macrophage IGF1 expression *in vivo*.

**Figure 8.**
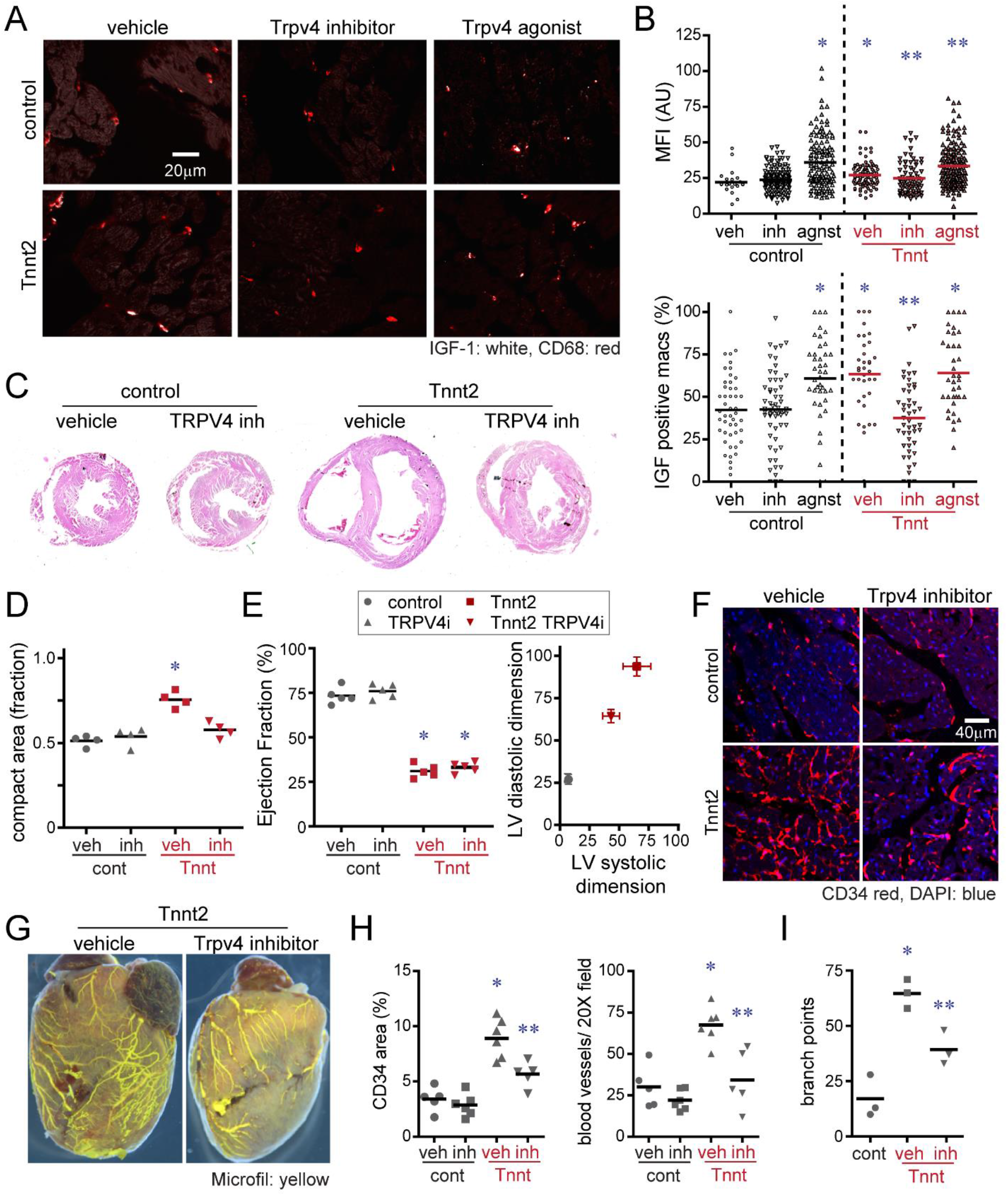
TRPV4 channel activity regulates IGF1 expression in CCR2− macrophages and is required for coronary angiogenesis. **A**, Immunostaining for IGF1 (white) and CD68 (red) in the LV myocardium of control and Tnnt2^ΔK210/ΔK210^ mice treated with either vehicle, TRPV4 inhibitor, or TRPV4 agonist demonstrating that TRPV4 channel activity regulates macrophage IGF1 protein expression *in vivo*. **B**, Quantification of IGF1 protein expression (% IGF1 + macrophages, IGF1 MFI) in control and Tnnt2^ΔK210/ΔK210^ hearts treated with either vehicle, TRPV4 inhibitor, or TRPV4 agonist. MFI: mean florescent intensity. Each data point represents an analyzed 20X field. n=5 animals per experimental group (A-B). * denotes p<0.05 compared to vehicle treated control hearts. ** denotes p<0.05 compared to vehicle treated Tnnt2^ΔK210/ΔK210^ hearts. (ANOVA, post-hoc Tukey). **C**, Low magnification H&E images of control and Tnnt2^ΔK210/ΔK210^ mice treated with either vehicle or TRPV4 inhibitor for 2 weeks beginning at 6 weeks of age. LV in cross-section (n=4 per experimental group). **D**, Quantification of the ratio of compact to trabecular myocardium in control and Tnnt2^ΔK210/ΔK210^ mice treated with either vehicle or TRPV4 inhibitor. Each data point represents an individual animal. * denotes p<0.05 compared to vehicle treated control hearts (ANOVA, post-hoc Tukey). **E**, Echocardiographic assessment of LV ejection fraction and LV chamber dimensions in control and Tnnt2^ΔK210/ΔK210^ mice treated with either vehicle or TRPV4 inhibitor for 2 weeks beginning at 6 weeks of age. n=5 per experimental group. * denotes p<0.05 compared to vehicle treated control hearts (ANOVA, post-hoc Tukey). **F**, Immunostaining for CD34 (red) in the LV myocardium of control and Tnnt2^ΔK210/ΔK210^ mice treated with either vehicle or TRPV4 inhibitor showing that TRPV4 channel activity contributes to coronary microvascular expansion in Tnnt2^ΔK210/ΔK210^ hearts. **G**, Microfil vascular casting showing that TRPV4 channel activity is necessary for expansion of coronary microvasculature in Tnnt2^ΔK210/ΔK210^ hearts. **H-I**, Quantification of coronary microvasculature (G) and coronary microvasculature (H) in the designated experimental groups. Each data point represents an individual animal (n=5 per experimental group). * denotes p<0.05 compared to vehicle treated control. ** denotes p<0.05 compared to all other groups (ANOVA, post-hoc Tukey).

To determine whether suppression of TRPV4 activity impairs reparative responses that are dependent on CCR2− macrophages, we treated control and Tnnt2^ΔK210/ΔK210^ mice with a TRPV4 inhibitor daily for 2 weeks beginning at 6 weeks of age. Histological evaluation demonstrated persistence of trabecular myocardium in Tnnt2^ΔK210/ΔK210^ hearts treated with the TRPV4 inhibitor, a phenotype that is reminiscent of depleting CCR2− macrophages (**Fig. 8C-D**). While TRPV4 inhibition had no impact on ejection fraction, we observed attenuated LV dilation in Tnnt2^ΔK210/ΔK210^ mice treated with the TRPV4 inhibitor (**Fig. 8E**). Examination of the coronary vasculature revealed that vehicle treated Tnnt2^ΔK210/ΔK210^ mice displayed evidence of coronary angiogenesis at the microvascular and macrovascular levels compared to vehicle treated control mice. Importantly, Tnnt2^ΔK210/ΔK210^ mice treated with the TRPV4 inhibitor displayed marked reductions in CD34+ blood vessel density within the LV myocardium and reduced large coronary artery complexity and branching compared to vehicle treated Tnnt2^ΔK210/ΔK210^ hearts. In contrast, treatment with the TRPV4 inhibitor had no impact on coronary microvascular density in control mice (**Fig. 8F-I**). Together, these observations demonstrate that TRPV4 channel activity is necessary for adaptive LV remodeling and coronary angiogenesis in the context of dilated cardiomyopathy and suggest a novel mechanism by which tissue resident cardiac macrophages contribute to the survival of the failing heart.

## Discussion

Inflammation has long been associated with heart failure development, progression, and prognosis. Strong clinical associations and mechanistic studies in model organisms have established that monocytes and macrophages contribute to adverse LV remodeling and heart failure pathogenesis. Unfortunately, clinical studies exploring the use of corticosteroids and tumor necrosis factor (TNF) antagonists in heart failure and myocardial infarction failed to show clinical efficacy dampening enthusiasm for the development of immunomodulatory therapies (Chung et al., 2003; Mann, 2015; Murphy et al., 2020; Parrillo et al., 1989). In fact, each of these treatment strategies was associated with potential harm. One explanation for these disappointing results is that distinct components of the innate immune system differentially contribute to disease pathogenesis, tissue homeostasis and repair. If this holds true, strategies that broadly target the innate immune system may have competing effects of not only limiting myocardial inflammation, but also, suppressing beneficial innate immune functions such as cardiac tissue repair.

Indeed, previous studies have strongly supported the division of labor concept in regards to cardiac macrophages, which are the most abundant innate immune cell population within the mouse and human heart (Bajpai et al., 2018; Epelman et al., 2014a; Pinto et al., 2016; Pinto et al., 2012). In this manuscript, we provide evidence that tissue resident CCR2−cardiac macrophages represent a protective population that mediates adaptive remodeling and survival of the chronically failing heart. By employing a mouse model of dilated cardiomyopathy harboring a causative human mutation, we demonstrate that CCR2− macrophages were essential to maintain adequate cardiac output in the setting of reduced cardiac contractility by promoting LV enlargement and expansion of the coronary system at the macrovascular and microvascular levels. Intriguingly, we revealed a novel mechanism of cardiac macrophage activation. Through formation of stable focal adhesion complexes with neighboring cardiomyocytes, CCR2− macrophages may sense mechanical stretch in response to elevated loading conditions (*i.e*., LV end diastolic pressure). The contribution of focal adhesion complexes to mechanosensing is well established (Geiger et al., 2009). Consistent with this notion, inhibition of the mechanosensitive channel, TRPV4, drastically reduced CCR2− macrophage pro-angiogenic growth factor expression and prevented coronary angiogenesis and myocardial tissue remodeling in our mouse model of dilated cardiomyopathy. Collectively, these findings establish an unanticipated role for cardiac macrophages in adaptive remodeling of the chronically failing heart and introduce a new mechanism of cardiac macrophage activation through sensing of myocardial stretch.

These findings have several important implications for the cardiovascular field. First, it is widely recognized that LV dilation is one of the strongest predictors of heart failure outcomes including mortality (Merlo et al., 2011). Whether this represents an associative or causative relationship is not immediately apparent and ultimately may depend on the underlying pathology and clinical context. Our observations indicate that LV dilation may be adaptive in some scenarios as it preserved cardiac output through augmentation of stroke volume. Second, previous studies have demonstrated that CCR2− macrophages remain abundant within the myocardium of patients with chronic heart failure (Bajpai et al., 2018). However, their function within this context was unknown. Using a mouse model of genetic dilated cardiomyopathy, we reveal an indispensable role for CCR2− macrophages. Depletion of CCR2− macrophages blunted myocardial tissue reorganization, cardiomyocyte lengthening, LV chamber enlargement, and coronary angiogenesis. Markers of pathological hypertrophy including increased cardiomyocyte cross-sectional area and fetal gene expression were not affected. These data further substantiate the division of labor between CCR2−and CCR2+ cardiac macrophage populations and highlight dichotomous contributions to disease pathogenesis and protective adaptations, respectively (Bajpai et al., 2019; Dick et al., 2019; Epelman et al., 2014a; Hulsmans et al., 2018; Lavine et al., 2014; Leid et al., 2016). Therapeutically, these findings indicate the need to develop strategies that preserve or enhance the function of CCR2− macrophages. Such an approach may enhance coronary angiogenesis and favor adaptive forms of LV remodeling, thus providing additive benefit to established medications for heart failure (ACE inhibitors, ARNIs, beta blockers, aldosterone antagonists), which target a separate mechanism (adverse remodeling).

Exciting work has recently implicated cardiac macrophages in facilitating electrical conduction, particularly through the atrioventricular node (Hulsmans et al., 2017). Using surface electrocardiography, we did not observe arrhythmias or clear alterations in PR, RR, or QT intervals following depletion of CCR2− macrophages. These data do not exclude the possibility that CCR2−and/or CCR2+ macrophages participate in aspects of cardiac pacemaker function, electrical propagation, or arrhythmia susceptibility in our dilated cardiomyopathy model. Additional depletion models and dedicated electrophysiology studies including appropriate provocative maneuvers will be required to address these important questions.

This study also provides new insights into the properties and functions of tissue resident macrophages. Specifically, we found that CCR2− macrophages within the LV myocardium display a stereotyped morphology where they interact with neighboring cardiomyocytes through the formation of focal adhesion complexes. It is not yet clear whether macrophages interact with the cardiomyocyte basement membrane or directly with cardiomyocytes themselves. Regardless, these structures are stable over time and have the potential to serve as sensors of mechanical deformations such as increased wall tension that may occur in the context of elevated preload or afterload. Consistent with this concept, we found that mechanical stretch serves as a stimulus for pro-angiogenic growth factor expression, a process that was dependent on the mechanoresponsive TRPV4 channel. This observation suggests that a key function of CCR2− macrophages may be to sense hemodynamic alterations and promote adaptive tissue remodeling. Future work will clarify the breadth of hemodynamic stimuli that might activate CCR2− macrophages, identify the exact signaling pathways triggered by TRPV4 channel activity, and determine whether this mechanism might be active in regions of the heart other than the LV.

As TRPV4 has been implicated in alveolar and intestinal macrophages (Hamanaka et al., 2010; Luo et al., 2018; Pairet et al., 2018), sensing of mechanical tissue deformation may constitute a conserved role of tissue resident macrophages throughout the body. The ability to directly interact with parenchymal cells and sensing mechanical inputs may reflect an additional division of labor between tissue resident and infiltrating monocyte-derived macrophages. TRPV4 may also play an essential role in phagocytosis-induced inflammation a mechanism that is involved in clearance of cellular debris following tissue injury (Dutta et al., 2020; Goswami et al., 2019; Mannaa et al., 2018; Scheraga et al., 2016). Interestingly, we found that activation of TRPV4 by cyclic mechanical stretch was not dependent on MYD88 or TRIF signaling. A recent manuscript has suggested that Piezo1 activation may trigger TRPV4 channel opening providing a more direct link to mechanical stimulation (Swain et al., 2020).

Our study is not without limitations. We primarily focused on a genetic mouse model of dilated cardiomyopathy. It remains to be shown whether CCR2− macrophages function in a similar manner in other non-ischemic and ischemic heart failure models. Based on available literature (Bajpai et al., 2019), we chose to employ CD169-DTR mice to deplete CCR2− macrophages. This line depletes other macrophage populations outside of the heart and we cannot rule out the possibility that extra-cardiac macrophage populations contribute to some of the observed phenotypes. Finally, while CCR2− macrophages represent the most abundant cell type that express active TRPV4 channels, we cannot exclude the possibility that TRPV4 may also influence the function of other cell types within the heart. TRPV4 was expressed in CCR2+ macrophages, neutrophils, and in a small population of cardiac endothelial cells. The impact of TRPV4 in these cell types remains to be elucidated.

In conclusion, our findings establish a role for tissue resident CCR2− macrophages in adaptive cardiac tissue remodeling and survival of the chronically failing heart. Furthermore, we provide initial evidence of a novel mechanism of cardiac macrophage activation, whereby CCR2− macrophages sense myocardial stretch through a TRPV4 dependent pathway.

## Acknowledgments

This project was made possible by funding provided from the Children’s Discovery Institute of Washington University and St. Louis Children’s Hospital (CH-II-2015-462, CH-II-2017-628), Foundation of Barnes-Jewish Hospital (8038-88), and the NHLBI (R01 HL138466, R01 HL139714). K.J.L. is supported by Burroughs Welcome Fund (1014782). Histology was performed in the DDRCC advanced imaging and tissue analysis core supported by Grant #P30 DK52574. Imaging was performed in the Washington University Center for Cellular Imaging (WUCCI) which is funded, in part by the Children’s Discovery Institute of Washington University and St. Louis Children’s Hospital (CDI-CORE-2015-505 and CDI-CORE-2019-813) and the Foundation for Barnes-Jewish Hospital (3770). D.K. is supported by NIH P01AI116501 and R01 HL094601, Veterans Administration Merit Review grant 1I01BX002730 and The Foundation for Barnes-Jewish Hospital. We acknowledge the McDonnel Genome Institute for their assistance in designing and performing the micro array and RNA sequencing analysis. We recognize the mouse cardiovascular phenotyping core for performing echocardiography and invasive hemodynamics studies. B.J. Kopecky was supported by the Principles in Cardiovascular Research Training Grant (T32 HL007081) and the Washington University Physician Scientist Training Program. H.H. was supported by grants from the NIH, R01DK103901, R01AR077183, and R01AA027065, the Department of Anesthesiology at Washington University School of Medicine. Y.L. is supported by NIH R35HL145212 and R01HL131908. XRM data was generated on a Zeiss Xradia Versa 520 3D X-Ray microscope which was purchased with support from the Office of Research Infrastructure Programs (ORIP), a part of the NIH Office of the Director under grant OD021694.

## Author Contributions

N.W., J.M., B.K., and S.G. performed the immunostaining, blood vessel casting, histology, RNA sequencing, and cell culture experiments. G.B, A.B., and O.D. performed the flow cytometry experiments. S.G. and B.K. performed the 2-photon imaging studies. H.L. and Y.L. performed the PET imaging studies and processed the data. L.E. and L.B. assisted in the cull culture experiments. I.K. and J.L. assisted in the macrophage depletion and performed the mitochondrial respiration studies. N.P. assisted in analysis of the ECG studies. S.M. provided Tnnt2^ΔK210^ mice. M.R.F, P.O.B. and J.A.J.F. performed the x-ray microscopy studies. L.D. and H.H. performed the ratiometric calcium imaging studies. C.M, A.K., J.M.N. performed the cardiac catheterization experiments. S.E., D.K., and R.S. assisted with experimental design and critical review of the manuscript. K.L. is responsible for all aspects of this manuscript including experimental design, data analysis, and manuscript production.

