## Supplemental data and methods for "Resident Cardiac Macrophages Mediate Adaptive Myocardial Remodeling"

### Materials and Methods

#### Animal Models

Mice were bred and maintained at the Washington University School of Medicine and all experimental procedures were done in accordance with the animal use oversight committee. Mouse strains utilized included Tnnt2$^{\Delta K210}$ (Du et al., 2007), Rosa26-td (Madisen et al., 2010), Rosa26-GCaMP6 (Madisen et al., 2015), Flt3-Cre (Boyer et al., 2011), CCR2$^{GFP}$ (Satpathy et al., 2013), CD169-DTR (Miyake et al., 2007), Tnnt2-DTR (Bajpai et al., 2019), CX3CR1$^{GFP}$CCR2$^{RFP}$ (Jung et al., 2000; Saederup et al., 2010), MYD88$^{-/-}$ (Hou et al., 2008), TRIF/TICAM1$^{-/-}$ (Hoebe et al., 2003), and TRPV4-GFP (Mutant Mouse Regional Resource Centers, MMRRC). All mice were on the C57/B6 background and genotyped according to established protocols. Equal numbers of male and female mice were included in all experiments. CCR2- macrophage depletion was induced by administering 100ng diphtheria toxin (Sigma) via intraperitoneal injection daily to CD169-DTR mice beginning at 6 weeks of age. GSK2193874 (10mg/kg) and GSK1016790A (0.1mg/kg) were administered to mice via IP injection daily for 2 weeks starting at 6 weeks of age.

#### Echocardiography, Invasive Hemodynamics, and Electrocardiography

Mouse echocardiography was performed in the Washington University Mouse Cardiovascular Phenotyping Core facility using the VisualSonics 770 Echocardiography System. Avertin (0.005 ml/g) was used for sedation based on previously established methods of infarct quantification (Kanno et al., 2002). 2D and M-mode images were obtained in the long and short axis views. Ejection fraction (EF) and LV dimensions were calculated using edge detection software and standard techniques. Measurements were performed on 3 independently acquired images per animal, by investigators who were blinded to experimental group. Each experimental group included at least 5 animals.

For invasive hemodynamic measurements, mice were anesthetized with 1% isoflurane and a 1.2 French pressure-volume catheter was position into the LV (ADVantage PV system, Scisense). Pressure and volume data were recorded using a Scisense 404-16 Bit Four Channel Recorder (Scisense) and analyzed with LabScribe2 Software (Scisense). Each experimental group included at least 5 animals.

For dobutamine cardiac catheterization studies, mice were anesthetized with a mixture of xylazine (10 mg/kg) and ketamine (100 mg/kg) administered i.p. Mice were then ventilated using a Harvard MicroVent. Open chest cardiac catheterization was performed by opening the chest wall just above the diaphragm and visualizing the apex of the heart. A small hole was formed by puncture with a 28 gauge needle and a 1.4 Fr Scisense catheter was inserted and advanced into the left ventricle of the mouse. The catheter was secured in the ventricle by a loop of suture on the chest wall. Hemodynamic measurements were then recorded. After acquisition of stable baseline data, dobutamine was serially infused at rates of 2, 4, 8, 16, 32, and 64 ng · g $BW^{-1}$ · $min^{-1}$ Data was analyzed using the Lab Scribe software for comparison of contractile performance between groups at each infusion rate.

Surface electrocardiography was performed using a MouseMonitor system (INDUS, Instruments). Recordings were obtained for 1 hour under inhaled anesthesia (1% isoflurane). Temperature was monitored and maintained using a heating pad. For data analysis, .csv files were imported into MATLAB with channels for time and voltage outputs for leads I, II, and III. The remainder of the analysis was performed using data from lead II. The signal was first filtered using Savitzky-Golay filtering based on previously established methods for ECG signal de-noising (Samann and Schanze, 2019). This filtering was performed twice so that both high frequency and low frequency noise could be filtered. The wavelet transform was then computed using the maximal overlap discrete wavelet transform from the MATLAB wavelet toolbox. The peaks of the QRS complexes were identified from the square of this transformed signal. For each QRS complex, the locations of the peak and minima on either side were identified. The largest peak above a specified threshold and within a specified range before each QRS complex was identified as the p-wave. A similar procedure was applied to capture the peak of the t-wave in the range following a QRS complex. Local minima were captured for signals determined by inspection to have inverted t-waves. After the P, Q, R, S, and T waves were identified, the RR, PR, and QT intervals as well as the QRS width were computed. Each experimental group included at least 4 animals.

**Micro-PET/CT**

The CCR2 (ECL1i: LGTFLKC) and CCR5 (DAPTA: D-$A_1$STTTNYT) targeting peptides were synthesized from D-form amino acids by CPC Scientific (Sunnyvale, CA). Maleimido-mono-amide-DOTA was purchased from Macrocyclics, Inc (Dallas, TX). DOTA-ECL1i and DOTA-DAPTA was synthesized as reported (Heo et al., 2019; Liu et al., 2017; Luehmann et al., 2014). Probes were radiolabeling with $^{64}$Copper ($^{64}$Cu). The radiolabeled compound was analyzed using radio-HPLC to ensure more than 95% radiochemical purity prior to animal studies. Mice were anesthetized with isoflurane and injected with 3.7 MBq of $^{64}$Cu-DOTA-ECL1i or 3.7 MBq $^{64}$Cu-DOTA-DAPTA in 100 µL of saline *via* the tail vein. Small animal PET scans (0 to 60 min dynamic scan) were performed on either microPET Focus 220 (Siemens, Malvern, PA) or Inveon PET/CT system (Siemens, Malvern, PA). The microPET images were corrected for attenuation, scatter, normalization, and camera dead time and co-registered with microCT images. All of the PET scanners were cross-calibrated periodically. The microPET images were reconstructed with the maximum a posteriori (MAP) algorithm and analyzed by Inveon Research Workplace. The uptake was calculated as the percent injected dose per gram (%ID/gram) of tissue in three-dimensional regions of interest (ROIs) without the correction for partial volume effect.

**Histology, Immunostaining, Picrosirius Red, and Wheat Germ Agglutinin Staining**. For histological analyses, tissues were fixed in 2% PFA overnight at 4°C, dehydrated in 70% ethyl alcohol, and embedded in paraffin. 4-μm sections were cut and stained with Picrosirius red using standard techniques. Picrosirius red staining was quantified using Image J software. To quantify cardiomyocyte cross-sectional area paraffin sections were stained with rhodamine conjugated WGA (Vector labs), visualized on a Zeiss confocal microscope, and measurements preformed using Zeiss Axiovision software.  For all immunostaining assays, tissues were fixed in 2% PFA overnight at 4°C, embedded in OCT, infiltrated with 30% sucrose, frozen, and 12-μm cryosections cut. Primary antibodies used were: CD68 clone FA-11 1:400 (Biolegend Cat# 137001), GFP (Abcam Cat# ab13970), RFP (Abcam Cat# ab62341), cardiac actin clone AC1-20.4.2 (Sigma Cat# A9357), $\alpha$-actinin clone BM-75.2 (Sigma Cat# A5044), Ki67 (Abcam Cat# ab15580), CD34 clone MEC14.7 (Abcam Cat# ab8158), Cx43 (Cell Signaling Cat# 3512), Paxillin clone Y113 (Abcam Cat# ab32084), Pan-Cadherin (Cell Signaling Cat# 4068), Claudin I (Cell Signaling cat# 13255), Desmoplakin I/II clone DP2.15 (Abcam Cat# ab16434), IGF1 (R&D systems Cat# AF791), CYR61 (R&D systems Cat# AF4055), Ly6G clone 1A8 (BD Cat# 551459), HCN4 (Fisher Cat# PA5-111878), FAK (Abcam, Cat# ab76496), and TRPV4 clone 1B2.6 (Millipore Cat# MABS466). Immunofluorescence was visualized using appropriate secondary antibodies on a Zeiss confocal microscopy system. For all experiments, at least 4 sections from 4 independent samples were analyzed in blinded fashion. Antibody specificity was validated using appropriate no primary and isotype controls. Co-localization was assessed by scoring images for the percent of macrophages expressing each junction marker at sites of cardiomyocyte interaction. Scoring was performed by 2 independent evaluators blinded to sample designation.

**Cardiomyocyte Morphometry**

Mouse hearts were harvested, cut into 1-2 mm tissue blocks, and fixed with 2% PFA at overnight at 4 degrees C. Tissue blocks were washed in PBS and then digested in Collagenase B (1.8 mg/mL) and Collagenase D (2.4 mg/mL) at 37 degrees C overnight (Han et al., 2020). Isolated cells were washed in PBS, fixed in 2% PFA, and imaged using an invested bright field microscope. Cardiomyocyte dimensions were measured using Image J.

**Coronary Vascular Perfusion and Casting**

To visualize the coronary arterial tress, Microfil perfusion reagent (yellow, Flow tech) was perfused retrograde into the ascending aorta per the manufacturer's specifications. The hearts were then fixed and imaged on a Zeiss Discovery V.12 stereomicroscope. Branching points were quantified as a surrogate measure of vascular complexity. To visualize the coronary microvasulature, anesthetized mice were injected IV with biotinylated tomato lectin (Vector labs) 5 minutes prior to tissue harvest. The hearts were then fixed, embedded in OCT, and cryosections stained with streptavidin-FITC (Vector labs).

**Electron Microscopy**

200 micron thick vibratome sections were taken from the heart tissue obtained from $CX3CR1^{GFP/+}CCR2^{RFP/+}$ mice (VT1200S, Leica Biosystems, Vienna, Austria). Samples were immersion fixed in 2% paraformaldehyde overnight and immunolabeled with GFP (Abcam cat# ab13970) or RFP (Abcam cat# ab62341) antibodies. Antibody staining was visualized with DAB and tissues were then submerged in a mixture of 2.5% glutaraldehyde and 2% paraformaldehyde in 0.15 M cacodylate buffer (pH 7.4) containing 2 mM calcium chloride and incubated overnight at 4˚C. Sample were rinsed in 0.15 M cacodylate buffer 3 times for 10 minutes each, and subjected to a secondary fixation step for one hour in 1% osmium tetroxide containing 1.5% potassium ferrocyanide in cacodylate buffer on ice. Following fixation, samples were then washed in ultrapure water 3 times for 10 minutes each and *en bloc* stained for 1 hour

with 2% aqueous uranyl acetate. After staining was complete, samples were briefly washed in ultrapure water, dehydrated in a graded acetone series (50%, 70%, 90%, 100% x2) for 10 minutes in each step, infiltrated with microwave assistance (Pelco BioWave Pro, Redding, CA) into LX112 resin, embedded in silicone molds, and cured in an oven at 60˚C for 48 hours. Once the resin was cured, each block was trimmed and faced using a diamond trim tool. 70nm longitudinal sections were obtained and imaged on a FE-SEM (Zeiss Crossbeam 540, Oberkochen, Germany) using the aSTEM detector. The SEM was operated at 28 KeV with a probe current of 0.9 nA, and the STEM detector was operated with the annular rings inverted for additional sample contrast. Large sample areas were imaged at a resolution of 4096 x 3072 pixels with a pixel size of 5.582 nm. At least 40 macrophages from 4 independent samples were included in the analysis.

**X-Ray Microscopy**

Whole hearts were extracted from mice, immediately bisected and immersion fixed in 2.5% glutaraldehyde and 2% paraformaldehyde in 0.15 M cacodylate buffer (pH 7.4) containing 2 mM calcium chloride overnight at 4˚C. Post-fixation, each heart was stained by immersion into Lugol's Iodine solution for a period of 5 days. Post-staining, hearts were mounted in a support matrix of 2% low-melt agarose within an Eppendorf tube to mitigate any movement during scanning. The Eppendorf tube was subsequently fixed onto a stub which was then mounted in the rotation holder within the X-Ray Microscope (Versa 520 XRM, Zeiss Microscopy, Pleasanton, CA). The samples were then imaged in the XRM at 80kV with an exposure time of 4 seconds, a pixel pitch of 5.41 μm with 2301 individual images acquired through a 360 degree rotation. Tomographic reconstruction was performed using the Zeiss 3DXMViewer software and rendered using ORS Dragonfly (ORS, Montreal, Quebec. Canada).

**Papillary Muscle Preparations and Live Two-Photon Microscopy**

Papillary muscles were dissected from the hearts of $CX3CR1^{GFP/+}$ $CCR2^{RFP/+}$ or CX3CR1-ertCre; Rosa26-GCaMP6/tdTomato mice as previously described (Uhl et al., 2015). Briefly, mouse hearts were harvested and perfused in oxygenated Krebs-Henseleit (KH) buffer preparation solution on ice. Left ventricles were exposed and papillary muscles were dissected from the left ventricular free walls. Dissected papillary muscles were transported in KH buffer on ice, covered from light to the Washington University Center for Cellular Imaging.
For static imaging, papillary muscles were placed into a 60mm glass bottom dish (Cellvis D60-30-1.5-N) and held in place with a slice hold-down (Warner Instruments 64-1419). Mounted papillary muscles were continuously bathed in pre-warmed (32°C) KH buffer with 95% oxygen/5% carbon dioxide. Pre-warmed KH buffer was gravity fed into one edge of the imaging dish while being continuously vacuum suctioned from the other end of the dish. Time-lapse imaging was performed on a Zeiss 880 inverted two-photon microscope. GFP+ macrophages were imaged with a 920nm laser (12% laser power) and captured with an LD Plan-neofluar 20x/0.4 objective (2-3x zoom, 4-line average) and Airyscan detectors. Z-stacks of ~80-100µM thickness were collected every 30 seconds for the duration of the time-lapse (90 minutes). Post-image processing was performed with ZEN3.0 (Blue version).

To apply axial tension, we generated a custom apparatus using both commercially available supplies (Mitutoyo 0.01mm 0-25mm micrometer, Thor Lab [XRN25/M, VC1/M, XRN-XZ/M, DT12A] supplies) and custom-made inserts. After transport to the imaging core in ice cold KH buffer, papillary muscles were carefully snagged without tension on microtweezers. Tension was then applied and papillary muscles imaged. Time-lapse imaging was performed on a Zeiss 880 inverted two-photon microscope. GFP+ macrophages (GCaMP6) were imaged with a 488nm laser (2.7% laser power) and TdT+ macrophages were imaged with a 561nm laser (3% power) and captured with an LD Plan-neofluar 20x/0.4 objective (2-3x zoom, 4-line average) and Airyscan detectors. Z-stacks of ~80-100µM thickness were collected every 30 seconds for the duration of the time-lapse (30 minutes). Baseline images were obtained over 5 minutes, and then vehicle (DMSO) was placed directly in the imaging dish and in the KH buffer and imaged for 10 minutes. After baseline and vehicle imaging, the feedline was flushed and TrpV4 inhibitor (GSK219 300nM) was directly added to the dish and to a second aliquot of pre-warmed KH buffer and papillary muscles imaged for an additional 10 minutes. Post-acquisition processing was performed in ZEN 3.0 SR (black edition). At least 30 macrophages from 4 independent samples were included in the analysis. Regions of interest were identified based on tdTomato staining and relative intensities of GFP (GCaMP6) were normalized against background and recorded over the duration of the experiment. The GFP intensity of each macrophage was averaged over the duration of baseline/vehicle imaging and compared to the averaged GFP intensity of axially stretched macrophages after administration of TrpV4 inhibitor.

**Flow cytometry**

Single cell suspensions were generated from saline perfused hearts by finely mincing and digesting them in DMEM with Collagenase 1 (450 U/ml), Hyaluronidase (60 U/ml) and DNase I (60 U/ml) for 1 hour at 37°C. All enzymes were purchased from Sigma. To deactivate the enzymes samples were washed with HBSS that was supplemented with 2% FBS and 0.2% BSA and filtered through 40 µM cell strainers. Red blood cell lysis was performed with ACK lysis buffer (Thermo Fisher Scientific). Samples were washed with HBSS and resuspended in 100 µL of FACS buffer (DPBS with 2% FBS and 2 mM EDTA). Cells were stained with monoclonal antibodies at 4°C for 30 minutes in the dark. All the antibodies were obtained from Biolegend. A complete list of antibodies is provided below. Samples were washed twice, and final resuspension was made in 300 µL FACS buffer. DAPI or LIVE/DEADTM Aqua dyes were used for exclusion of dead cells. Immune cells were first gated as CD45+. Neutrophils were gated as Ly6G$^{high}$CD64-. Monocytes were gated as CD64$^{int}$Ly6C$^{high}$CCR2+MHCII$^{low}$. Macrophages were gated as Ly6G-CD64$^{high}$Ly6C$^{low}$ cells. Flow cytometric analysis and sorting were performed on BD LSRII, BD FACS ARIAIII, BD FACS Melody platforms.

CD45-PerCP/Cy5.5, clone 30-F11 (Biolegend cat# 103131)

CD64-APC, PE, PE/Cy7 clone X54-5/7.1 (Biolegend cat# 139305, 139303, 139313)

CCR2-BV421, clone: SA203G11 (Biolegend cat# 150605)

MHCII-APC/Cy7, clone M5/114.15.2 (Biolegend cat# 107627)

Ly6G-PE/Cy7, clone 1A8 (Biolegend cat# 127617)

Ly6C-APC and FITC, clone HK1.4 (Biolegend cat# 128015, 128005)

CD31-APC, clone 390 (Biolgend cat# 102409)

Anti-Feeder-PE, clone mEFSK4 (Miltenyi cat# 130-120-166)

**RT-PCR**

RNA was extracted from sorted cells or myocardial tissue using the RNeasy RNA mini kit and Tissue Lyser II (Qiagen). RNA concentration was measured using a nanodrop spectrophotometer (ThermoFisher Scientific). cDNA synthesis was performed using the High Capacity RNA to cDNA synthesis kit (Applied Biosystems). For sorted macrophages, cDNA was synthesized using the iScript™ Reverse Transcription Supermix (Bio-Rad) and pre-amplified using the Sso Advanced PreAmp Supermix kit (Bio-Rad). Quantitative real time PCR reactions were prepared with sequence-specific primers (IDT) with PowerUP™ Syber Green Master mix (ThermoFisher Scientific) in a 20 µL volume. Real time PCR was performed using QuantStudio 3 (ThermoFisher Scientific). mRNA expression was normalized to 36B4. All RT-PCR assays were performed using validated primer sets (IDT) with appropriate quality controls including melt curves and negative controls.

**RNA sequencing**

Total RNA was harvested from by disrupting samples in Trizol (Thermo) using the Qiagen TissueLyser homogenizer with stainless steel beads. Following phenol chloroform extraction, RNA was purified using the Ambion Purelink miniprep kit. RNA integrity was quantified on an Agilent Bioanalyzer and samples with RIN>8 utilized for RNA sequencing. Ribosomal RNAs were depleted with Ribo-Zero, cDNA libraries generated, and samples sequenced (1X50bp reads) on an Illumina HiSeq 3000 instrument. Sequence alignment, normalization, and differential expression analysis was carried out in the McDonnel Genome Institute at Washington University using Limma-Voom software. PCA and Hierarchical cluster analysis was performed in Partek Genomics. GO Pathway analysis was performed using DAVID. Transcripts with >10 reads in 50% of samples demonstrating a fold change >1.5 fold at an FDR<0.05 were included in the pathway analysis. Differential gene expression was performed using START (Nelson et al., 2017) and Partek Genomics analysis packages.

**Microarray**

To isolated RNA, macrophages were directly sorted into QLT buffer containing 2-mercaptoethanol and RNA isolated using the RNeay micro kit (Qiagen) per manufacturer's instructions. Gene expression profiling was performed using microarray analysis in collaboration with the McDonnell Genome Institute at Washington University. RNA was amplified using the WTA (Sigma) system and hybridized to Agilent 8X60 gene chips. Data analysis was performed using Partek genome suite software.

**MMP assay**

Protein was extracted from mouse hearts using EDTA-free RIPA buffer and the Tissue Lyser II (Qiagen). Equal concentrations of protein were loaded onto zymogram gels (Novex) and run at 125 V constant for 90 minutes on the XCell SureLock Mini-Cell. The gels were then developed and incubated in 1X Zymogram Renaturing Buffer and 1X Zymogram Developing Buffer (Novex) per the manufacturer's instructions. Gels were then stained with SimplyBlue Safestain (Invitrogen), imaged, and quantified using Image J software.

**Cell Culture**

To generate bone-marrow derived macrophages (BMDMs), isolated bone marrow cells from mouse femurs and tibiae were cultured for 7 days in DMEM supplemented with 10% FBS, 5% M-CSF, 5% horse serum, 1% streptomycin and 1% sodium pyruvate. On day 8 the media was replaced with M-CSF free media. For cell stretch experiments, BMDMs were plated on fibronectin and collagen coated silicone membranes and cultured overnight to facilitate adherence. Seeded membranes were then subjected to cyclic unaxial stretch (1 Hz, 10% deformation) for 24-48 hours. GSK2193874 was used at 1 μM. LPS and polyIC were used at 100 ng/ml and 20 μg/ml, respectively.

For cardiomyocyte-macrophage co-culture experiments, 6-well plates were coated with fibronectin and incubated at 37°C overnight. HL-1 cells were then cultured in Claycomb media (Sigma) supplemented with 2% L-glutamine for 24 hours. BMDMs were then co-cultured at a 1:6 ratio of BMDMs:HL-1 cells, at which point the co-culture was either treated with Itgb1/CD29 clone HM β1-1 (10 μM) or CD18 clone M18/2 (20 μM). Four hours after co-culture the cells were imaged at 20x and the number of BMDM-HL-1 interactions per 20x field were quantified. Cells were defined as interacting if a measurable BMDM projection made contact with an HL-1 cardiomyocyte.

**Ratiometric Calcium assays**

CCR2- and CCR2+ macrophages isolated from the heart by flow cytometry were resuspended in culture medium containing 10 % FBS, 100 U/mL penicillin, 100 μg/mL streptomycin, plated on

5 mm coverslips coated with poly-L-lysine (10 μg/mL) and cultured under a humidified atmosphere of 5 % CO2 / 95% air at 37°C for 2 h. Then, cultured cardiac macrophages were loaded with 4 μM Fura-2 AM (Invitrogen, Carlsbad, CA) in culture medium at 37°C for 60 min as previous described (Luo et al., 2018). Cells were washed three times and incubated in HBSS at room temperature for 30 min before use. Fluorescence at 340 and 380 nm excitation wavelengths was recorded on an inverted Nikon Ti-E microscope equipped with 340 and 380 nm excitation filter wheels using NIS Elements imaging software (Nikon). Fura-2 ratios (F340/F380) were used to reflect changes in intracellular Ca2+ upon stimulation. The threshold of cellular activation was defined as 20% above the baseline. GSK1016790A and GSK2193874 were used at 300 nM. Ionomycin (1 μM) was used as a positive control.

**Mitochondrial Isolation and Respiration Assays**

One day prior to mitochondrial isolation, XFe96 Sensor cartridges were hydrated by adding 180uL ultrapure H2O to each well and incubating the plate in a non CO2 incubator overnight. Mitochondria were isolated from mouse heart tissue as previously described (Frezza et al., 2007). Following hemodynamic study, mice were sacrificed through cervical dislocation. Using a surgical scalpel and scissors, hearts were removed rapidly and immersed in a small beaker containing 5 mL of ice-cold PBS supplemented with 10mM EDTA. Heart tissue was minced into small pieces using scissors, washed thrice with ice-cold PBS supplemented with 10mM EDTA over a 40um filter, spun down at 200g for 5 min, and the supernatant discarded. Heart tissue was then resuspended in 1mL IBm1 (Prepare 100 ml of IBm1 by mixing 6.7 ml of 1M sucrose, 5ml of 1M Tris/HCl, 5 ml of 1M KCl, 1 ml of 1M EDTA, and 2 ml of 10% BSA. Adjust pH to 7.4. Bring the volume to 100 ml with distilled water) and hand homogenized using a pre-cooled Teflon pestle ten times. The homogenate was then transferred to a 1.7mL microcentrifuge tube and centrifuged at 700g for 10 min at 4oC. Supernatant was then transferred to a new 1.7mL microcentrifuge tube and centrifuged at 8,000g for 10 min at 4oC. Remaining supernatant was removed, and the pellet was resuspended in 1mL IBm2 (Prepare 100 ml of IBm2 by mixing 25 ml of 1 M sucrose, 3 ml of 0.1 M EGTA/Tris, and 1 ml of 1 MTris/HCl. Adjust pH to 7.4. Bring the volume to 100 ml with distilled water). The suspension was centrifuged at 8,000g for 10 min at 4oC and the supernatant discarded. The residual supernatant was used to resuspend the pellet, and mitochondrial concentration was measured using BCA Protein Assay.

Mitochondrial Respiration was measured as follows.  Mitochondria were resuspended in MAS1 buffer (Prepare MAS1 by dissolving 220 mM of d-Mannitol, 70 mM of sucrose, 10 mM of KH2PO4, 5 mM of MgCl2, 2 mM of HEPES, 1 mM of EGTA, and 0.2% (w/v) of fatty acid-free BSA in ultrapure H2O and adjust the pH to 7.2 with KOH at 37 °C) containing either 10mM glutamate & 5mM malate (Complex I) or 5mM succinate & 2uM rotenone (Complex II). A volume of 50uL Mitochondria were plated on Seahorse XFe96 microculture plates at a concentration of 4ug mitochondria per well. Plates were centrifuged using a swinging bucket-microplate adaptor at 2000g for 20 min at 4oC. Seahorse sensor cartridges were loaded with 10X aliquots (ADP, oligomycin, FCCP, antimycin & rotenone) and calibrated during this step. Mitochondria were inspected by microscopy. Following centrifugation, 130uL MAS1 with substrates (glutamate/malate or succinate/rotenone) were added to each well for a total volume of 180uL. Plate was warmed at 37oC in a non-CO2 incubator for 5-10 min, after which it was placed into the Seahorse Bioanalyzer.

**Serum Chemistry**. Serum was collected by LV puncture prior to mouse harvest. Chemistries and electrolytes were measured in the Washington University School of Medicine Division of Comparative Medicine Research Animal Diagnostic Laboratory using a AMS Liasys 330 Clinical Chemistry System.

**Statistical Analysis.** Data were analyzed by using software (Prism, version 6.0-7.0; GraphPad, La Jolla, Calif). Differences between groups were compared by using Mann-Whitney U test. Multiple means were compared by using 1-way ANOVA (analysis of variance) with the post-hoc Tukey test. $p<0.05$ (two-sided) was indicative of a statistically significant difference. Bonferroni correction was performed when multiple hypotheses were tested. Data are presented as dot plots or box whisker plots generated in PRISM. The sample size used to calculate statistical significance is stated in the appropriate figure legend.

**Data Availability.** Source Data for all experiments have been provided. All other data are available from the corresponding author on reasonable request. Gene expression data will be deposited in GEO at the time of publication.

### KEY RESOURCES TABLE

| REAGENT or RESOURCE | SOURCE | IDENTIFIER |
|---|---|---|
| **Antibodies** | | |
| CD68 | Biolegend | Cat# 137001 |
| GFP | Abcam | Cat# ab13970 |
| RFP | Abcam | Cat# ab62341 |
| Cardiac actin | Sigma | Cat# A9357 |
| $\alpha$-actinin | Sigma | Cat# A5044 |
| Ki67 | Abcam | Cat# ab15580 |
| Paxillin | Abcam | Cat# ab32084 |
| Cx43 | Cell Signaling | Cat# 3512 |
| Pan-Cadherin | Cell Signaling | Cat# 4068 |
| Claudin I | Cell Signaling | Cat# 13255 |
| Desmoplakin I/II | Abcam | Cat# ab16434 |
| Itgb1/CD29 | BD | Cat# 562219 |
| Itgb2/CD18 | BD | Cat# 557437 |
| TRPV4 | Millipore | Cat# MABS466 |
| IGF1 | R&D systems | Cat# AF791 |
| CYR61 | R&D systems | Cat# AF4055 |
| CD34 | Abcam | Cat# ab8158 |
| MHCII-APC/Cy7 | Biolegend | Cat# 107627 |
| CCR2-BV421 | Biolegend | Cat# 150605 |
| CD45-PerCP/Cy5.5 | Biolegend | Cat# 103131 |
| CD64-PE/Cy7, -PE, -APC | Biolegend | Cat# 139305, 139303, 139313 |
| Ly6G-PE/Cy7 | Biolegend | Cat# 127617 |
| Ly6C-APC, -FITC | Biolegend | Cat# 128015, 128005 |
| CD31-APC | Biolegend | Cat# 102409 |
| MEFSK4-PE | Miltenyi | Cat# 130-120-166 |
| HCN4 | Fisher | Cat# PA5-111878 |
| Ly6G | BD | Cat# 551459 |
| FAK | Abcam | Cat# ab76496 |
| **Bacterial and Virus Strains** | | |
| none | | |
| **Biological Samples** | | |
| none | | |
| **Chemicals, Peptides, and Recombinant Proteins** | | |
| RGD peptides | Selleckchem | Cat# S8008 |
| GSK1016790A | Selleckchem | Cat# S8107 |
| GSK2193874 | Tocris | Cat# 5106 |
| Ionomycin | Sigma | Cat# I9657 |
| Microfil | Flow Tech Inc. | Cat# MV-122 |
| Biotinylated Tomato Lectin | Vector labs | Cat# B-1175 |
| Wheat Germ Agglutinin | Vector labs | Cat# RL-1022 |
| **Critical Commercial Assays** | | |
| none | | |
| **Deposited Data** | | |

| | | |
|---|---|---|
| Raw RNA sequencing data | deposited in GEO at time of publication | |
| Raw microarray data | deposited in GEO at time of publication | |
| **Experimental Models: Cell Lines** | | |
| HL1 cells | | |
| **Experimental Models: Organisms/Strains** | | |
| Mouse: B6;129-Tnnt2tm2Mmto | CARD | CARD: 1585 |
| Mouse: B6;129-Siglec1<tm1(HBEGF)Mtka> | RIKEN BRC | RBRC04395 |
| Mouse: B6.129-Tg(Flt3-cre)#Ccb/Ieg | EMMA | EMMA: 17790 |
| Mouse: B6.Cg-Gt(ROSA)26Sortm9(CAG-tdTomato)Hze/J | Jackson Laboratory | JAX: 007909 |
| Mouse: B6(C)-Ccr2tm1.1Cln/J | Jackson Laboratory | JAX: 027619 |
| Mouse: Tg(Trpv4-EGFP)MT43Gsat/Mmucd | Mutant Mouse Resource and Research Centers | MMRRC: 032771-UCD |
| Mouse: B6.129(Cg)-Cx3cr1tm1Litt Ccr2tm2.1Ifc/JernJ | Jackson Laboratory | JAX: 032127 |
| Mouse: B6.129P2(SJL)-Myd88tm1.1Defr/J | Jackson Laboratory | JAX: 009088 |
| Mouse: C57BL/6J-Ticam1Lps2/J | Jackson Laboratory | JAX: 005037 |
| Mouse: B6J.Cg-Gt(ROSA)26Sortm95.1(CAG-GCaMP6f)Hze/MwarJ | Jackson Laboratory | JAX: 028865 |
| **Oligonucleotides** | | |
| IGF1 | IDT | Mm.PT.58.5811533 |
| HB-EGF | IDT | Mm.PT.58.10448014 |
| CYR61 | IDT | Mm.PT.58.10039559 |
| IL-1β | IDT | Mm.PT.58.41616450 |
| TNF | IDT | Mm.PT.58.12575861 |
| IL6 | IDT | Mm.PT.58.10005566 |
| IP10 | IDT | Mm.PT.58.43575827 |
| MX2 | IDT | Mm.PT.58.29837402 |
| NPPA | IDT | Mm.PT.58.12973594 |
| NPPB | IDT | Mm.PT.58.8584045 |
| MYH7 | IDT | Mm.PT.58.17465550 |
| **Recombinant DNA** | | |
| none | | |
| **Software and Algorithms** | | |
| Partek Genomics | | |
| START | Nelson et al., 2017 | https://doi.org/10.1093/bioinformatics/btw624 |
| Graph Pad PRISM | | |
| ImageJ | Schneider et al., 2012 | https://imagej.nih.gov/ij/ |
| ZEN 3.0 SR (black edition) | | |
| Zeiss 3DXMViewer | | |
| ORS Dragonfly | | |

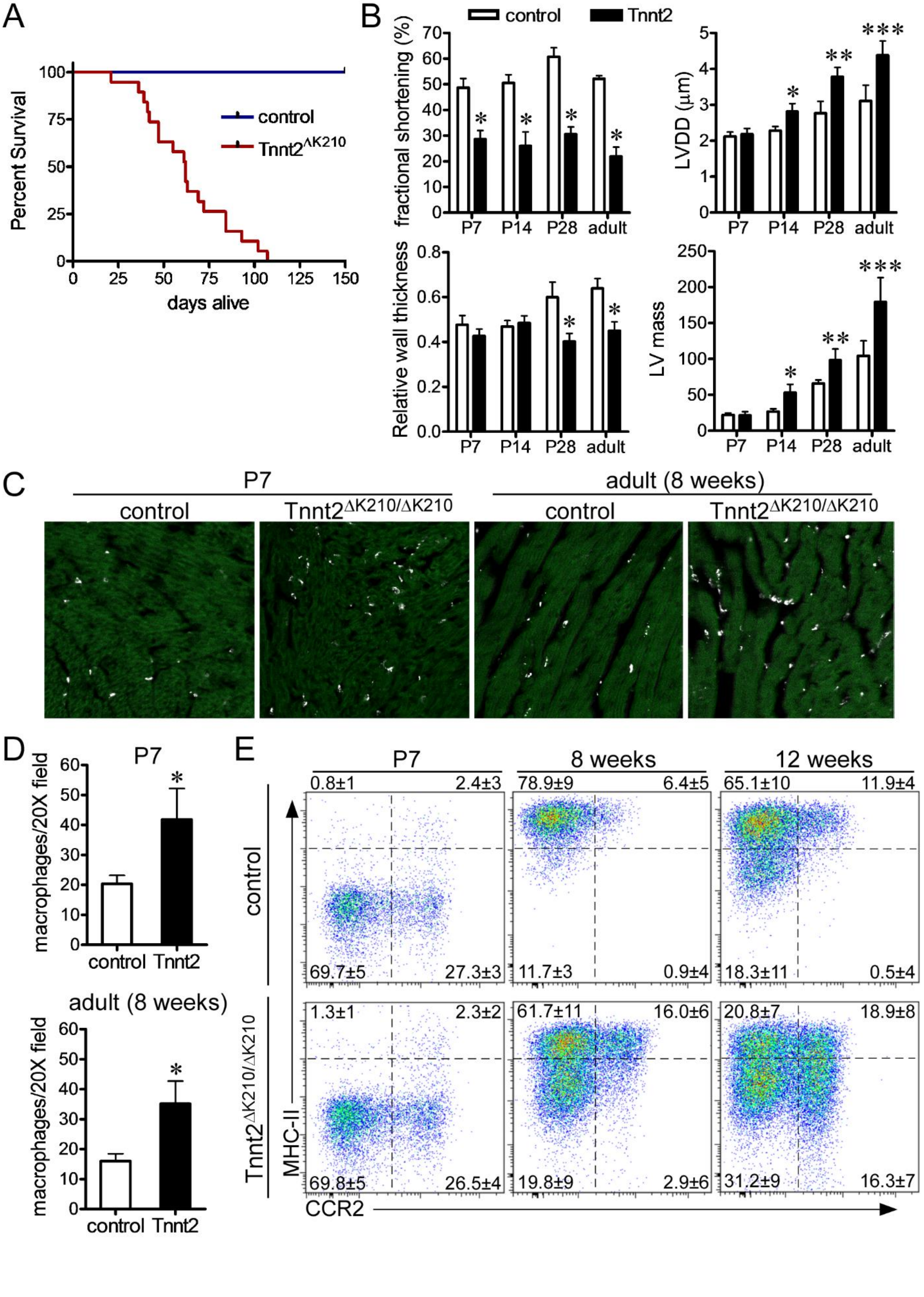
A
Percent Survival
days alive
control
Tnnt2ΔK210
B
control
Tnnt2
fractional shortening (%)
LVDD (μm)
Relative wall thickness
LV mass
P7
P14
P28
adult
C
P7
adult (8 weeks)
control
Tnnt2ΔK210/ΔK210
D
P7
adult (8 weeks)
macrophages/20X field
control
Tnnt2
E
P7
8 weeks
12 weeks
control
Tnnt2ΔK210/ΔK210
MHC-II
CCR2
0.8±1
2.4±3
69.7±5
27.3±3
78.9±9
6.4±5
11.7±3
0.9±4
65.1±10
11.9±4
18.3±11
0.5±4
1.3±1
2.3±2
69.8±5
26.5±4
61.7±11
16.0±6
19.8±9
2.9±6
20.8±7
18.9±8
31.2±9
16.3±7

**Figure S1** (related to Figure 1). **Cardiac Phenotype of Tnnt2$^{\Delta K210}$ mice across the spectrum of age.** **A**, Kaplan-Meier curve showing reduced survival of Tnnt2$^{\Delta K210/\Delta K210}$ mice compared to controls (n>20 per group). **B**, Echocardiographic analysis of control and Tnnt2$^{\Delta K210/\Delta K210}$ mice at postnatal days (P) 7, 14, 28, and adult time points (8 weeks of age). * denotes p<0.05 compared to control, ** denotes p<0.05 compared to other Tnnt2$^{\Delta K210/\Delta K210}$ groups, *** denotes p<0.05 compared to all other groups. (ANOVA, post-hoc Tukey). n=5-6 per experimental group. LVDD: LV end diastolic dimension. **C-D**, Immunostaining of control and Tnnt2$^{\Delta K210/\Delta K210}$ hearts showing increased numbers of CD68+ (white) macrophages in Tnnt2$^{\Delta K210/\Delta K210}$ hearts compared to controls at P7 and 8 weeks of age. Green: cardiac actin. * denotes p<0.05 compared to controls. Mann-Whitney test. n=6 per experimental group. **E**, Flow cytometry of CD45+CD64+Ly6G- macrophages showing shifts in macrophage composition in control and Tnnt2$^{\Delta K210/\Delta K210}$ hearts across the spectrum of age. n=6 per experimental group.

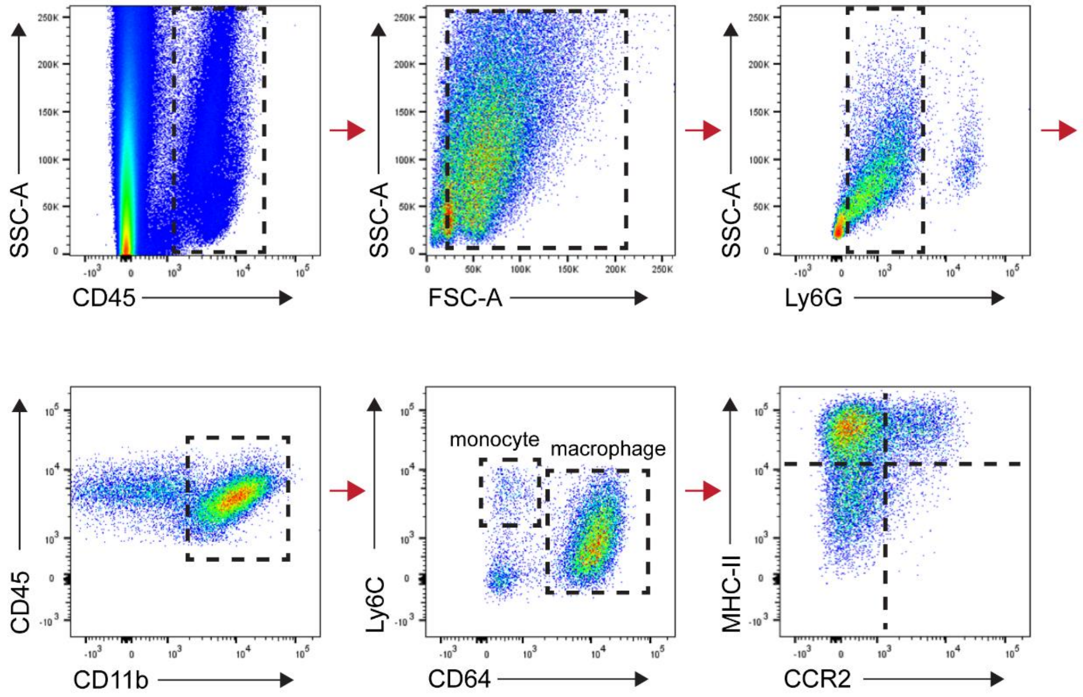


**Figure S2** (related to Figure 1). **Flow cytometry gating strategy.**

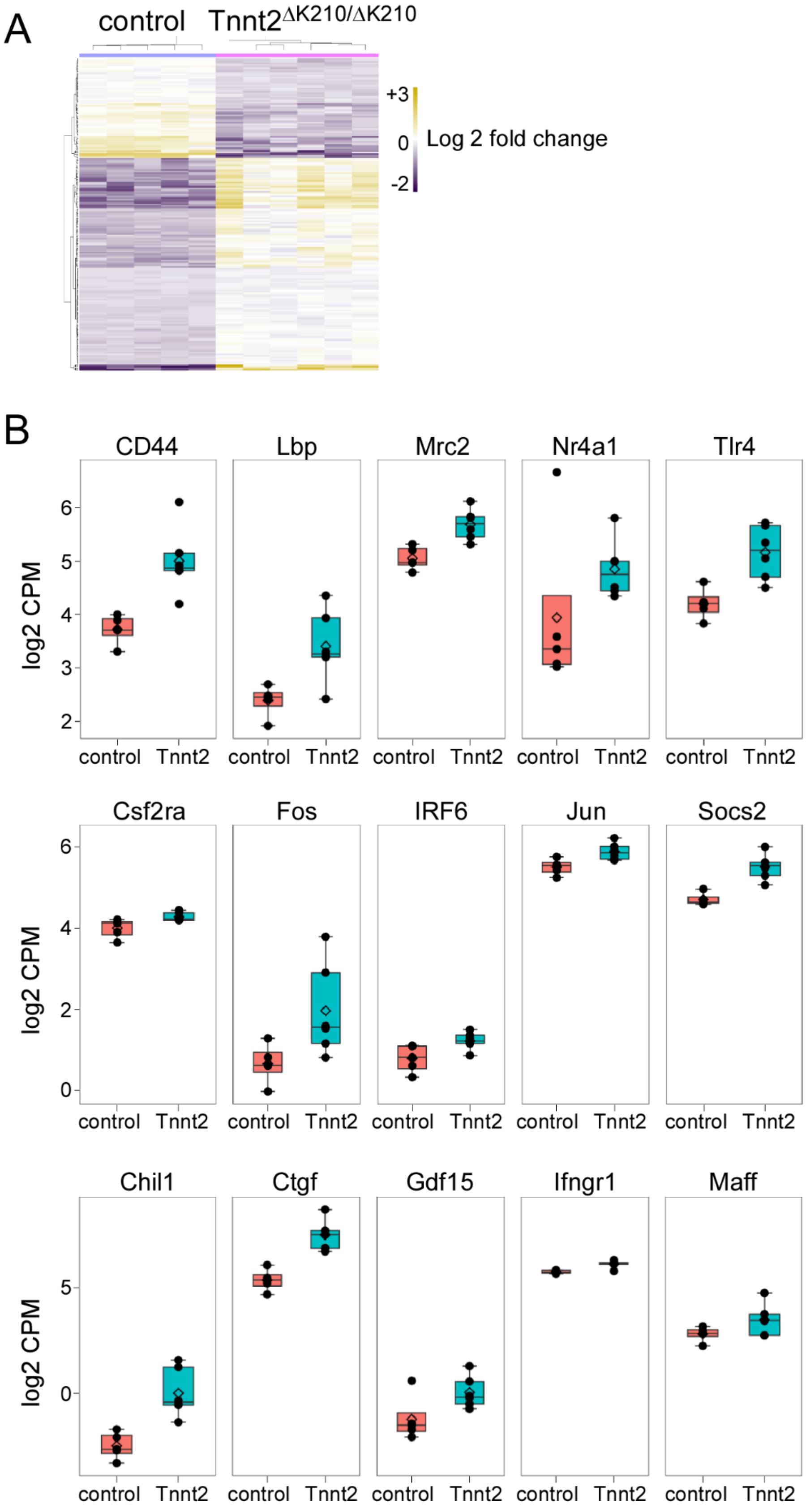
A
control
Tnnt2ΔK210/ΔK210
+3
0
-2
Log 2 fold change
B
CD44
Lbp
Mrc2
Nr4a1
Tlr4
Csf2ra
Fos
IRF6
Jun
Socs2
Chil1
Ctgf
Gdf15
Ifngr1
Maff
log2 CPM
control
Tnnt2

**Figure S3** (related to Figure 1). **RNA sequencing of control and Tnnt2$^{\Delta K210}$ mice.** **A**, Heat map showing genes differentially expressed in control and Tnnt2$^{\Delta K210/\Delta K210}$ hearts at 8 weeks of age (n=5-6 per experimental group). **B**, Box and whisker plots of genes related to macrophages and innate immunity that were differentially expressed in Tnnt2$^{\Delta K210/\Delta K210}$ hearts compared to controls. CPM: counts per million. FDR $p<0.05$ (control vs Tnnt2) for comparisons.

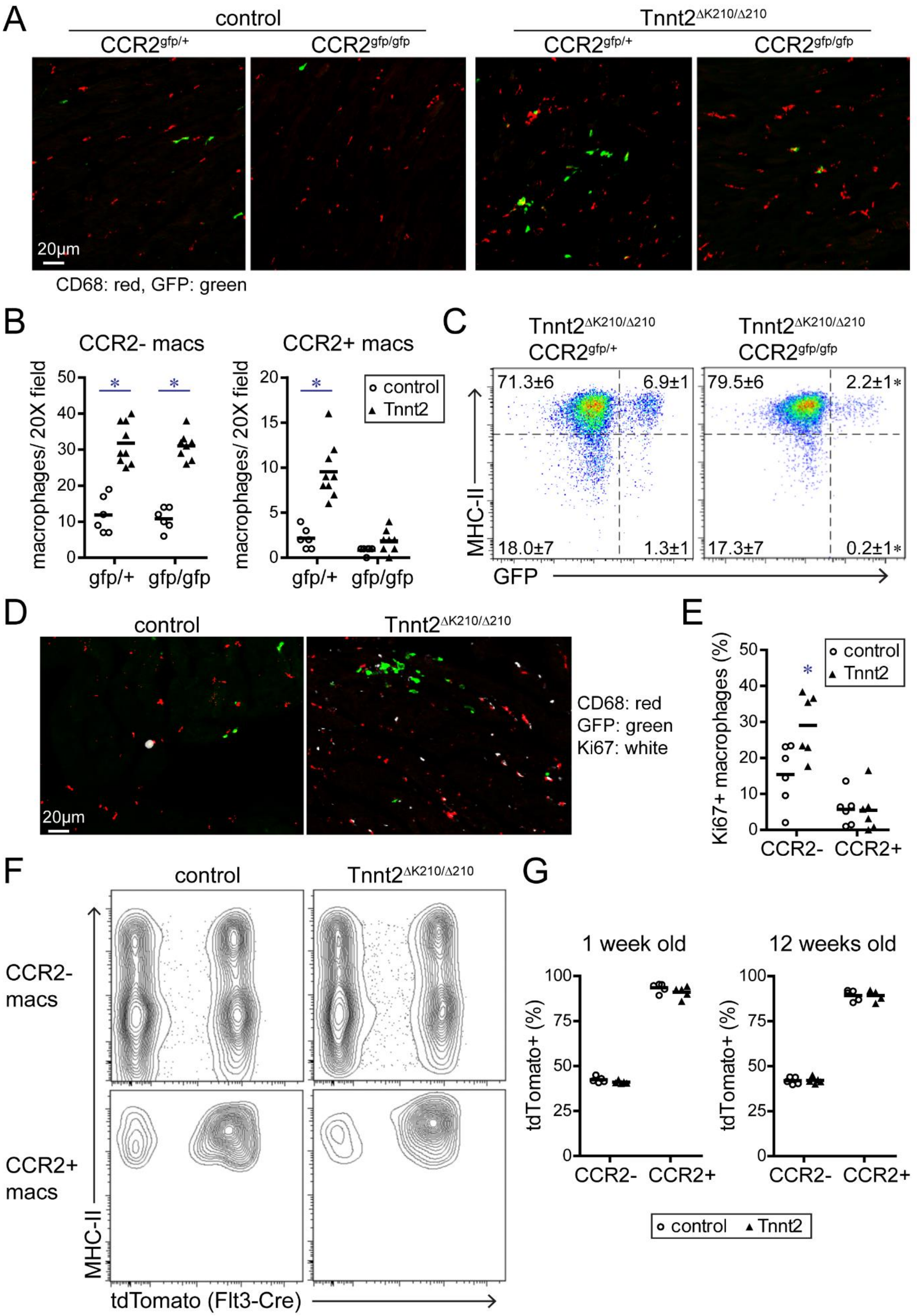

A
control
Tnnt2∆K210/∆210
CCR2gfp/+
CCR2gfp/gfp
CCR2gfp/+
CCR2gfp/gfp
20μm
CD68: red, GFP: green
B
CCR2- macs
CCR2+ macs
macrophages/ 20X field
gfp/+
gfp/gfp
control
Tnnt2
C
Tnnt2∆K210/∆210 CCR2gfp/+
Tnnt2∆K210/∆210 CCR2gfp/gfp
71.3±6
6.9±1
18.0±7
1.3±1
79.5±6
2.2±1*
17.3±7
0.2±1*
MHC-II
GFP
D
control
Tnnt2∆K210/∆210
20μm
CD68: red
GFP: green
Ki67: white
E
Ki67+ macrophages (%)
control
Tnnt2
CCR2-
CCR2+
F
control
Tnnt2∆K210/∆210
CCR2- macs
CCR2+ macs
MHC-II
tdTomato (Flt3-Cre)
G
1 week old
12 weeks old
tdTomato+ (%)
CCR2-
CCR2+
control
Tnnt2

**Figure S4** (related to Figure 2). **Recruitment dynamics and origins of cardiac macrophages in control and Tnnt2$^{\Delta K210/\Delta K210}$ mice. A**, Immunostaining for CD68 (red) and GFP (green) in control (CCR2$^{gfp/+}$), CCR2 knockout (CCR2$^{gfp/gfp}$), Tnnt2$^{\Delta K210/\Delta K210}$ CCR2$^{gfp/+}$, and Tnnt2$^{\Delta K210/\Delta K210}$ CCR2$^{gfp/gfp}$ hearts at 8 weeks of age. Reprentative images of n=6-8 per experimental group. **B**, Quantification of CCR2- and CCR2+ macrophages in the LV myocardium of CCR2$^{gfp/+}$, CCR2$^{gfp/gfp}$, Tnnt2$^{\Delta K210/\Delta K210}$ CCR2$^{gfp/+}$ ,and Tnnt2$^{\Delta K210/\Delta K210}$ CCR2$^{gfp/gfp}$ hearts. * denotes p<0.05 compared to control (Mann-Whitney test). n=6-8 per experimental group. **C**, Flow cytometry of CD45+CD64+Ly6G- macrophages in Tnnt2$^{\Delta K210/\Delta K210}$ CCR2$^{gfp/+}$ and Tnnt2$^{\Delta K210/\Delta K210}$ CCR2$^{gfp/gfp}$ hearts at 8 weeks of age showing specific reductions in only CCR2+ macrophages. * denotes p<0.05 compared to Tnnt2$^{\Delta K210/\Delta K210}$ CCR2$^{gfp/+}$ hearts (Mann-Whitney test). n=6 per experimental group. **D**, Immunostaining for CD68 (red), GFP (green), and Ki67 (white) in control and Tnnt2$^{\Delta K210/\Delta K210}$ hearts at 8 weeks of age showing increased proliferation of CCR2- macrophages in Tnnt2$^{\Delta K210/\Delta K210}$ hearts. n=6 per experimental group **E**, Quantification of the percent of CCR2- and CCR2+ macrophages that expressed Ki67. * denotes p<0.05 compared to controls (Mann-Whitney test). n=6-8 per experimental group. **F-G**, Flow cytometry of CD45+CD64+Ly6G- macrophages in control (Flt3-Cre Roas26-tdTomato) and Tnnt2$^{\Delta K210/\Delta K210}$ Flt3-Cre Rosa26-tdTomato hearts at 1 week and 12 weeks of age showing that the percent of CCR2- and CCR2+ macrophages derived from definitive hematopoietic progenitors (Flt3-Cre Rosa26-tdTomato+) remains unchanged between control and Tnnt2$^{\Delta K210/\Delta K210}$ hearts. B, E, G: each data point represents a biologically independent replicate (n=5-6 per experimental group).

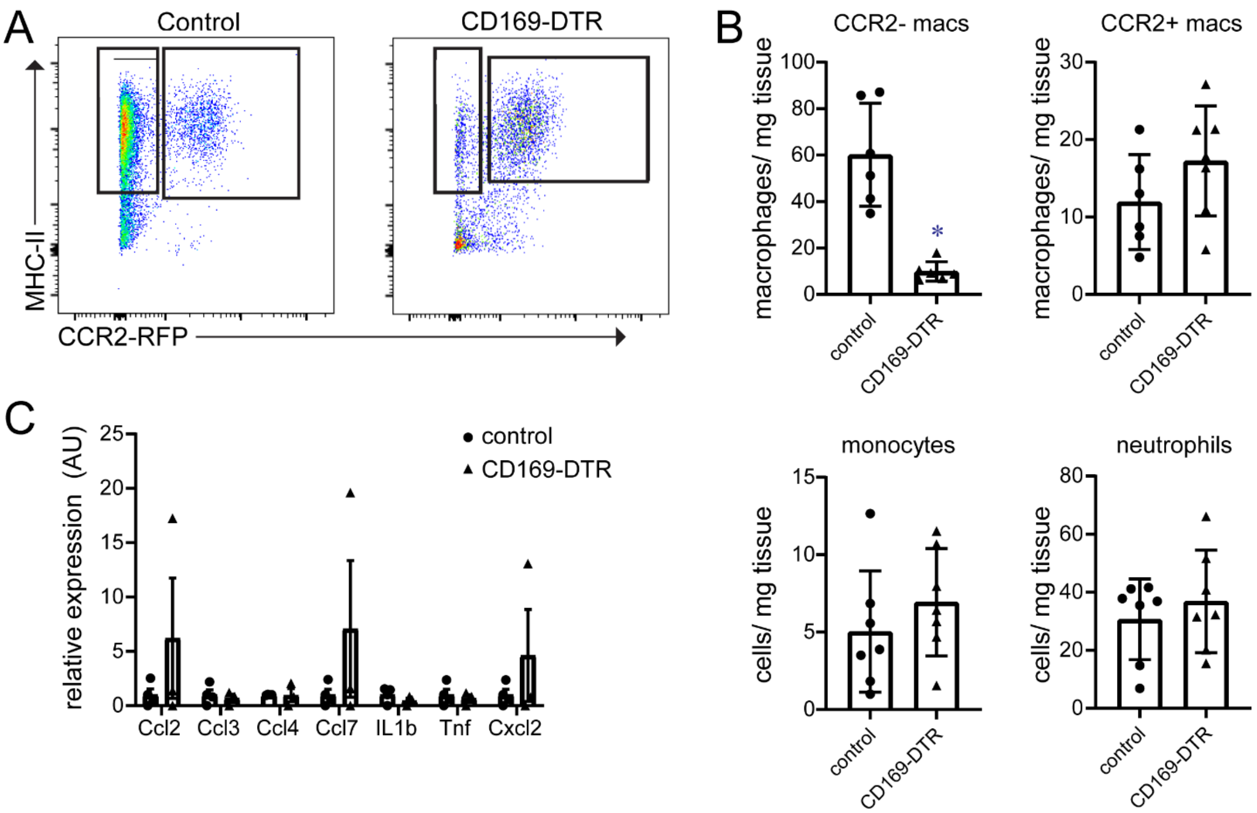


**Figure S5. CCR2- macrophage ablation** (related to Figure 3). **A**, Flow cytometry of macrophages (CD45+Ly6G- CD11b+ CD64+) isolated from the hearts of control (CX3CR1-GFP/+ CCR2-RFP/+) and CD169-DTR (CD169-DTR CX3CR1-GFP/+ CCR2-RFP/+) mice. n=6 per experimental group. **B**, Quantification of flow cytometry data showing depletion of CCR2- macrophages and no changes in the abundance of CCR2+ macrophages, monocytes, or neutrophils. Each data point denotes a biologically independent sample (n=6 per experimental group). * denotes $p<0.05$ compared to controls (Mann-Whitney test). **C**, Quantitative RT-PCR for chemokine and cytokine mRNA expression in CCR2+ macrophages isolated from control (CX3CR1-GFP/+ CCR2-RFP/+) and CD169-DTR (CD169-DTR CX3CR1-GFP/+ CCR2-RFP/+) hearts. Each data point denotes a biologically independent sample (n=4-6 per experimental group). No statistically significant differences were observed (Mann-Whitney test).

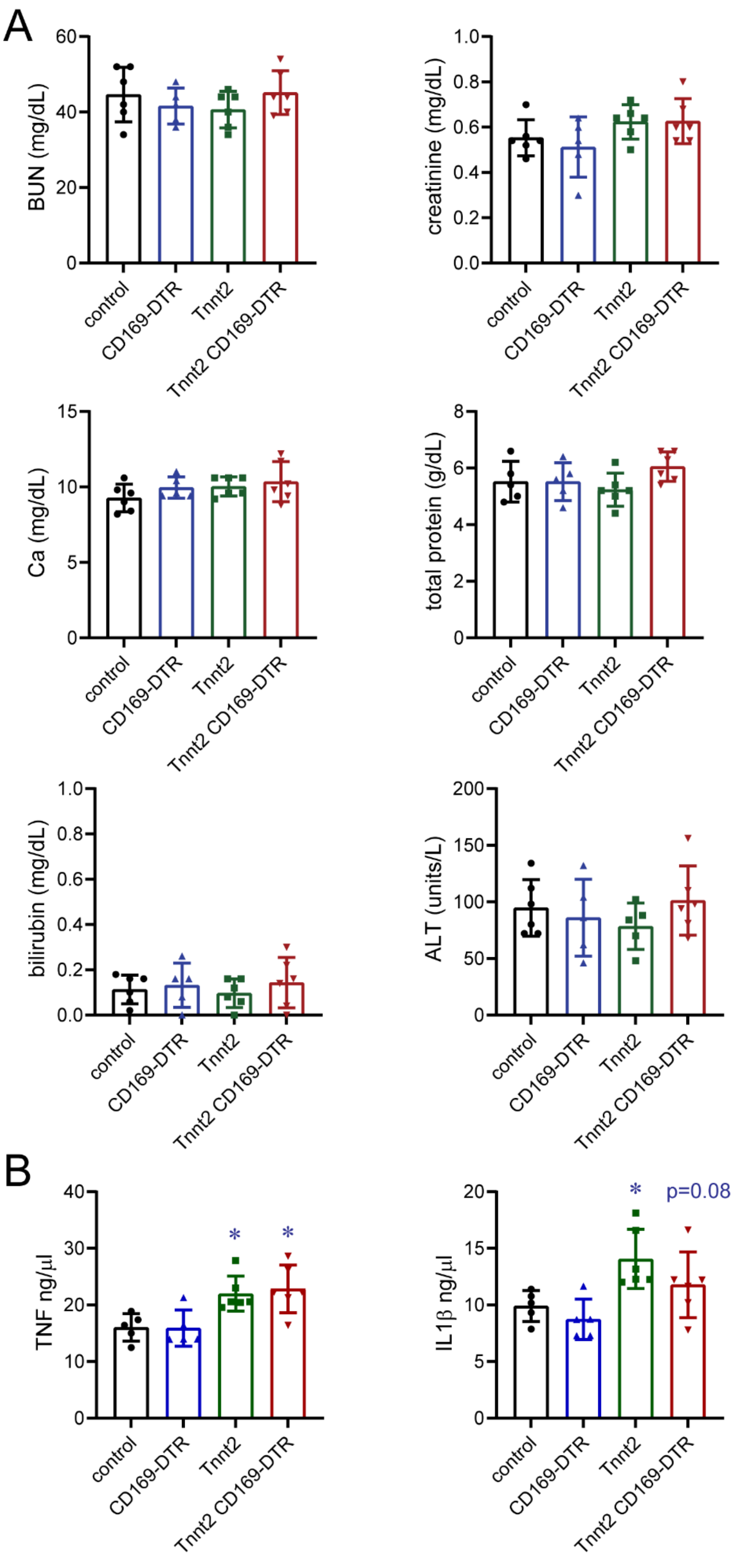
A
BUN (mg/dL)
creatinine (mg/dL)
Ca (mg/dL)
total protein (g/dL)
bilirubin (mg/dL)
ALT (units/L)
B
TNF ng/µl
IL1β ng/µl
*
p=0.08
control
CD169-DTR
Tnnt2
Tnnt2 CD169-DTR

**Figure S6** (related to Figure 3). **Serum chemistry and cytokine measurements**. **A**, Measurement of serum BUN, creatinine, calcium (Ca), total protein, bilirubin, and alanine aminotransferase (ALT) in control, CD169-DTR, Tnnt2$^{\Delta K210/\Delta K210}$, and Tnnt2$^{\Delta K210/\Delta K210}$ CD169-DTR hearts after 3 weeks of DT treatment. **B**, Measurement of serum IL1$\beta$ and TNF levels in control, CD169-DTR, Tnnt2$^{\Delta K210/\Delta K210}$, and Tnnt2$^{\Delta K210/\Delta K210}$ CD169-DTR mice after 3 weeks of DT treatment. * denotes $p<0.05$ compared to controls. $p=0.08$ for control vs Tnnt2 CD169-DTR. No statistically significant differences were observed between Tnnt2 and Tnnt2 CD169-DTR mice. Each data point represents a biologically independent replicate. n=5-6 per experimental group.

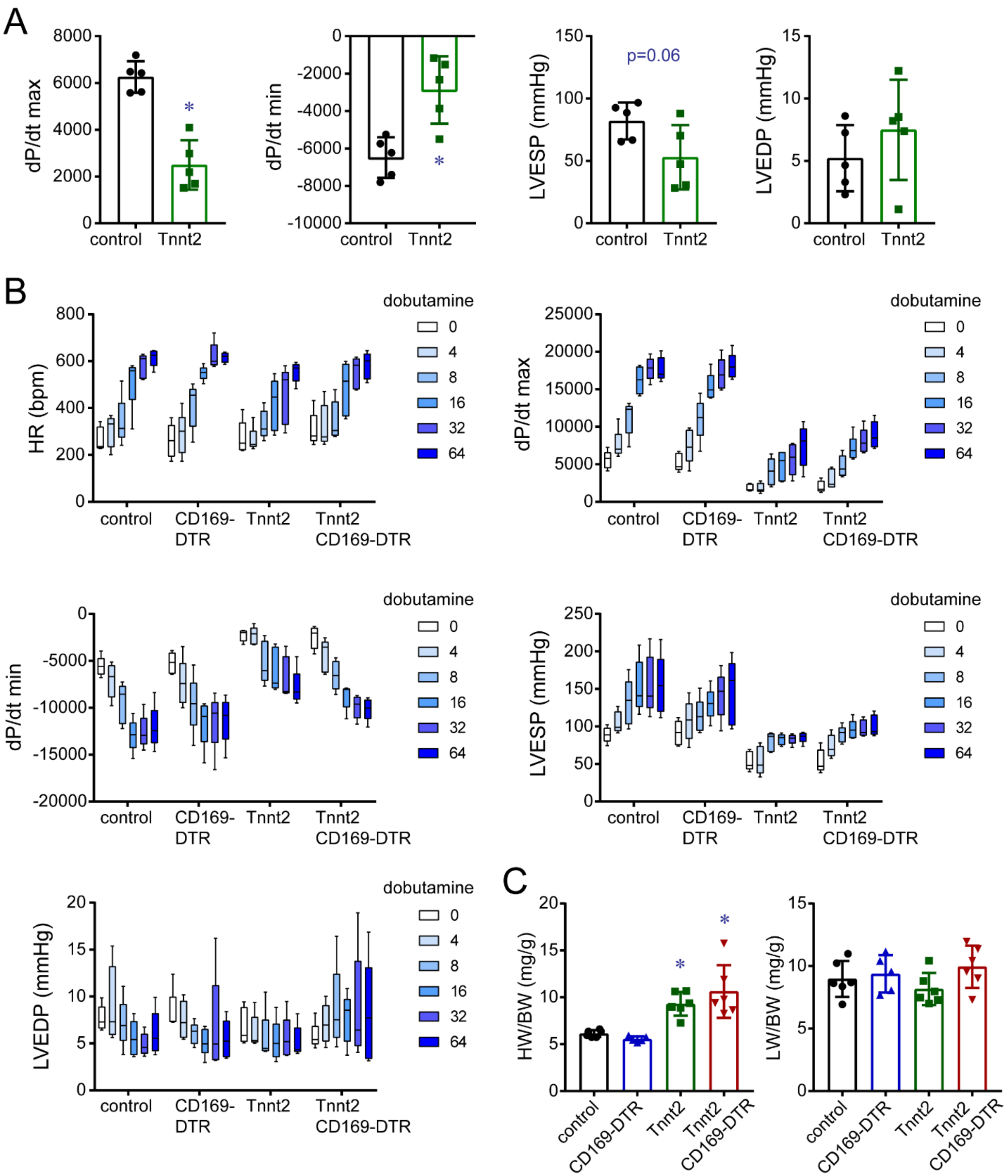
A
dP/dt max
control
Tnnt2
*
dP/dt min
*
LVESP (mmHg)
p=0.06
LVEDP (mmHg)
B
HR (bpm)
dobutamine
0
4
8
16
32
64
CD169-DTR
Tnnt2 CD169-DTR
dP/dt max
dP/dt min
LVESP (mmHg)
LVEDP (mmHg)
C
HW/BW (mg/g)
LW/BW (mg/g)

**Figure S7** (related to Figure 3). **Invasive hemodynamics**. **A**, LV pressure recordings of dP/dt max (mmHg/sec), dP/dt min, LV end systolic pressure (LVESP), and LV end diastolic pressure (LVEDP) in control and $Tnnt2^{\Delta K210/\Delta K210}$ mice at 4 weeks of age. * denotes p<0.05 compared to controls. Each data point represents individual animals. **B**, Measurements of heart rate (HR, beats per minute), dP/dt max, dP/dt min, LVESP, and LVEDP in 9 week old control, CD169-DTR, $Tnnt2^{\Delta K210/\Delta K210}$, and $Tnnt2^{\Delta K210/\Delta K210}$ CD169-DTR hearts after 3 weeks of DT treatment. Mice received escalating doses of dobutamine (ng/ml) via intravenous infusion. Data is presented as box and whisker plots. n=5-7 per experimental group. **C**, Measurements of heart weight (HW) relative to body weight (BW) and lung weight (LW) relative to body weight (BW) in 9 week old control, CD169-DTR, $Tnnt2^{\Delta K210/\Delta K210}$, and $Tnnt2^{\Delta K210/\Delta K210}$ CD169-DTR mice after 3 weeks of DT treatment. * denotes p<0.05 compared to controls. Each data point represents an individual animal. n=5-8 per experimental group.

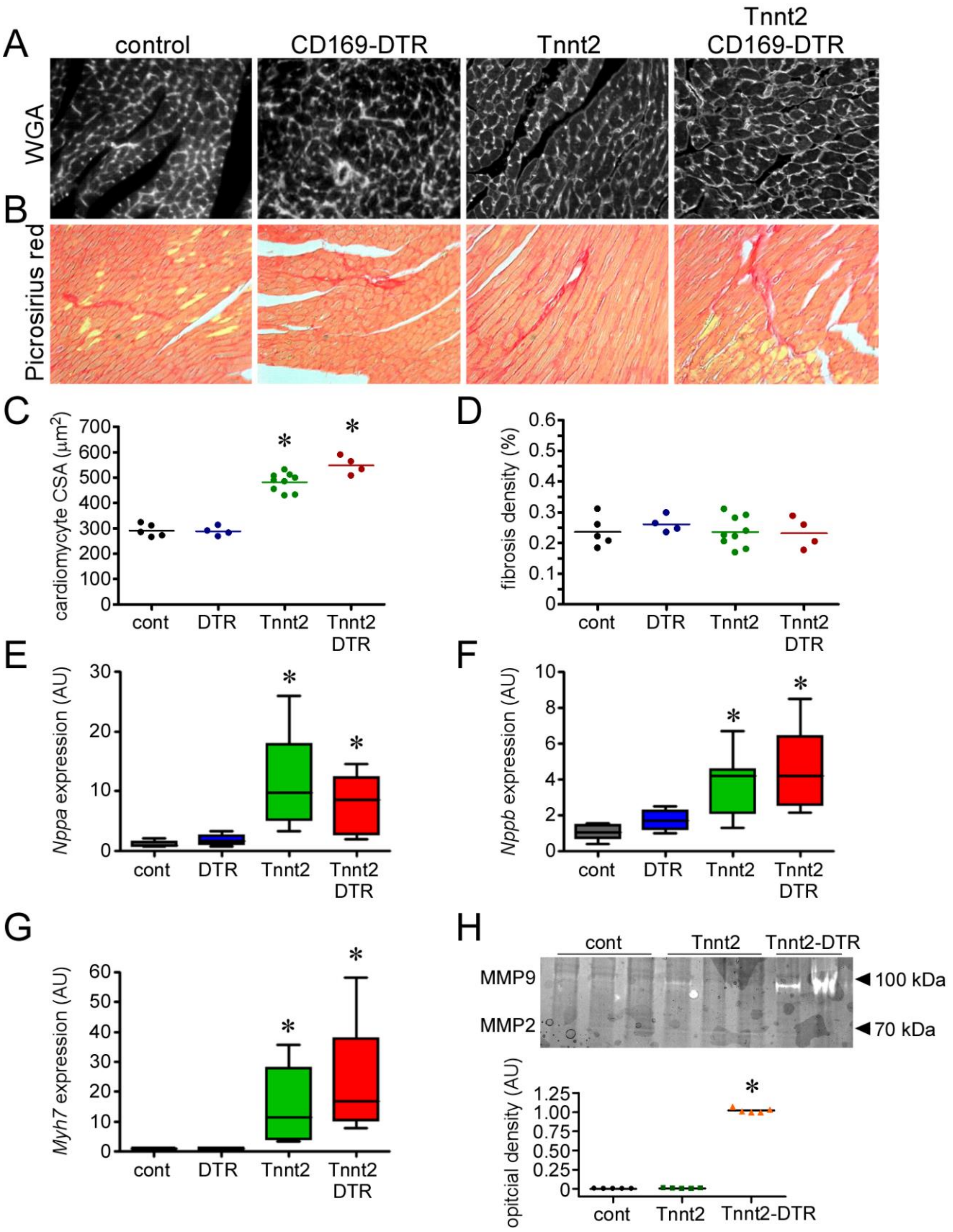

A
control
CD169-DTR
Tnnt2
Tnnt2 CD169-DTR
WGA
B
Picrosirius red
C
cardiomycyte CSA (μm2)
cont
DTR
Tnnt2
Tnnt2 DTR
D
fibrosis density (%)
E
Nppa expression (AU)
F
Nppb expression (AU)
G
Myh7 expression (AU)
H
cont
Tnnt2
Tnnt2-DTR
MMP9
MMP2
100 kDa
70 kDa
opitcial density (AU)

**Figure S8** (related to Figure 4). **CCR2- macrophages are not required for adverse LV remodeling. A-B**, Wheat germ agglutinin (WGA, A) and Picrosirius red (B) staining of control, CD169-DTR, Tnnt2$^{\Delta K210/\Delta K210}$, and Tnnt2$^{\Delta K210/\Delta K210}$ CD169-DTR hearts after 3 weeks of DT treatment. **C-D**, Quantification of cardiomyocyte cross-sectional area (C) and fibrotic density (D) in the hearts of control, CD169-DTR, Tnnt2$^{\Delta K210/\Delta K210}$, and Tnnt2$^{\Delta K210/\Delta K210}$ CD169-DTR mice after 3 weeks of DT treatment. * denotes p<0.05 compared to controls. Each data point represents a biologically independent replicate. **E-G**, Box and whisker s of Nppa, Nppb, and Myh7 mRNA expression in control, CD169-DTR, Tnnt2$^{\Delta K210/\Delta K210}$, and Tnnt2$^{\Delta K210/\Delta K210}$ CD169-DTR hearts after 3 weeks of DT treatment. * denotes p<0.05 compared to controls. n=5-6 per experimental group. **H**, Gelatinase activity assay showing minimal metallomatrix proteinase (MMP) activity in control and Tnnt2$^{\Delta K210/\Delta K210}$ hearts. Tnnt2-DTR mice (cardiomyocyte ablation) treated with DT were included as a positive control as they display marked increase in MMP activity. * denotes p<0.05 compared to controls. Each data point represents a biologically independent replicate. n=4-9 per experimental group.

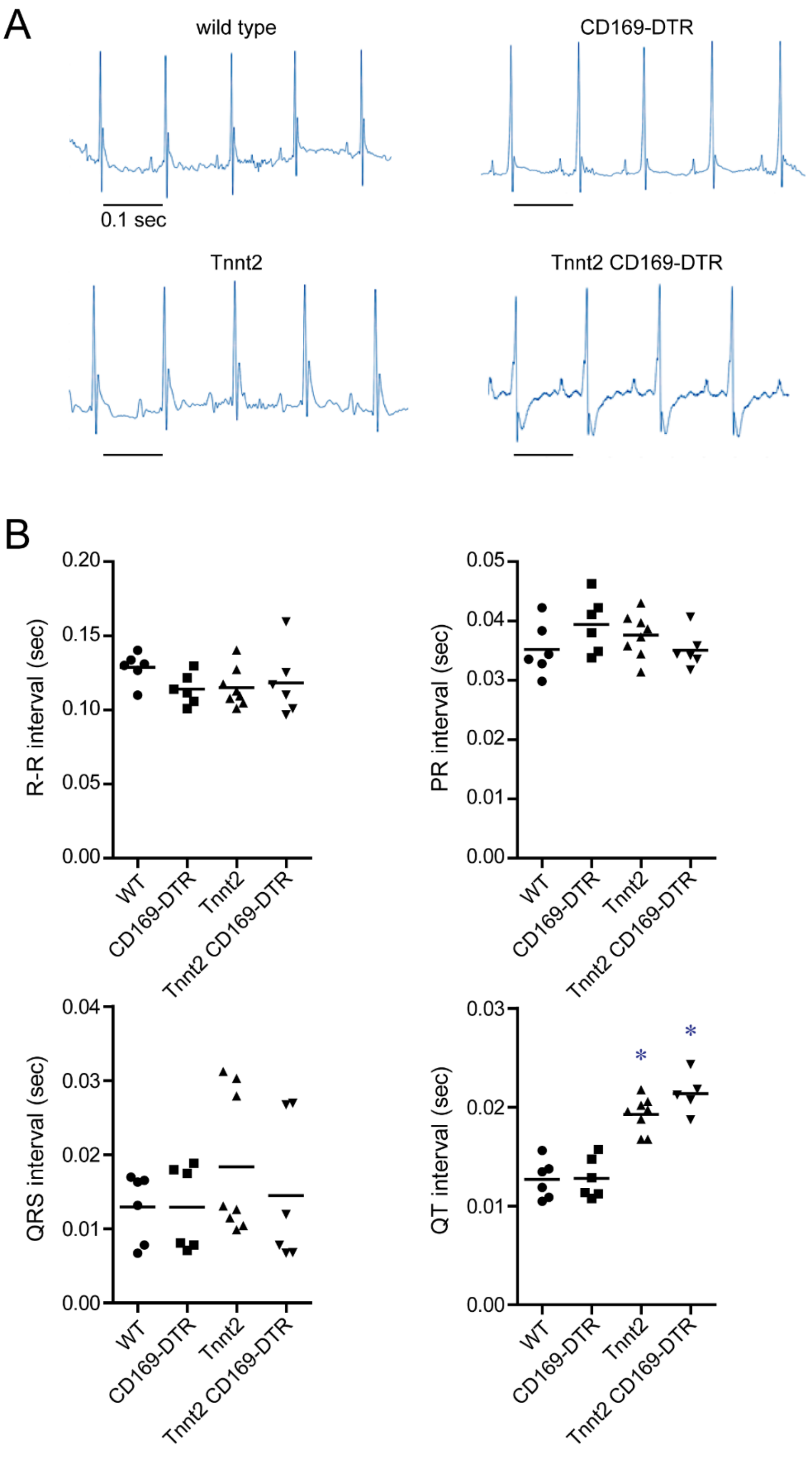

A
wild type
CD169-DTR
0.1 sec
Tnnt2
Tnnt2 CD169-DTR
B
R-R interval (sec)
0.00
0.05
0.10
0.15
0.20
WT
CD169-DTR
Tnnt2
Tnnt2 CD169-DTR
PR interval (sec)
0.00
0.01
0.02
0.03
0.04
0.05
QRS interval (sec)
0.00
0.01
0.02
0.03
0.04
QT interval (sec)
0.00
0.01
0.02
0.03
*
*

**Figure S9** (related to Figure 4). **ECG analysis does not reveal conduction delays in hearts depleted of CCR2- macrophages. A**, Representative ECG tracings of control (wild type), CD169-DTR, Tnnt2$^{\Delta K210/\Delta K210}$, and Tnnt2$^{\Delta K210/\Delta K210}$ CD169-DTR hearts after 3 weeks of DT treatment. **B**, Quantification of R-R, PR. QRS, and QT intervals in each experimental group. Prolongation of the QT interval was observed in Tnnt2$^{\Delta K210/\Delta K210}$, and Tnnt2$^{\Delta K210/\Delta K210}$ CD169-DTR hearts compared to controls. * denotes p<0.05 compared to controls. Each data point represents a biologically independent replicate. 6-8 mice per experimental group.

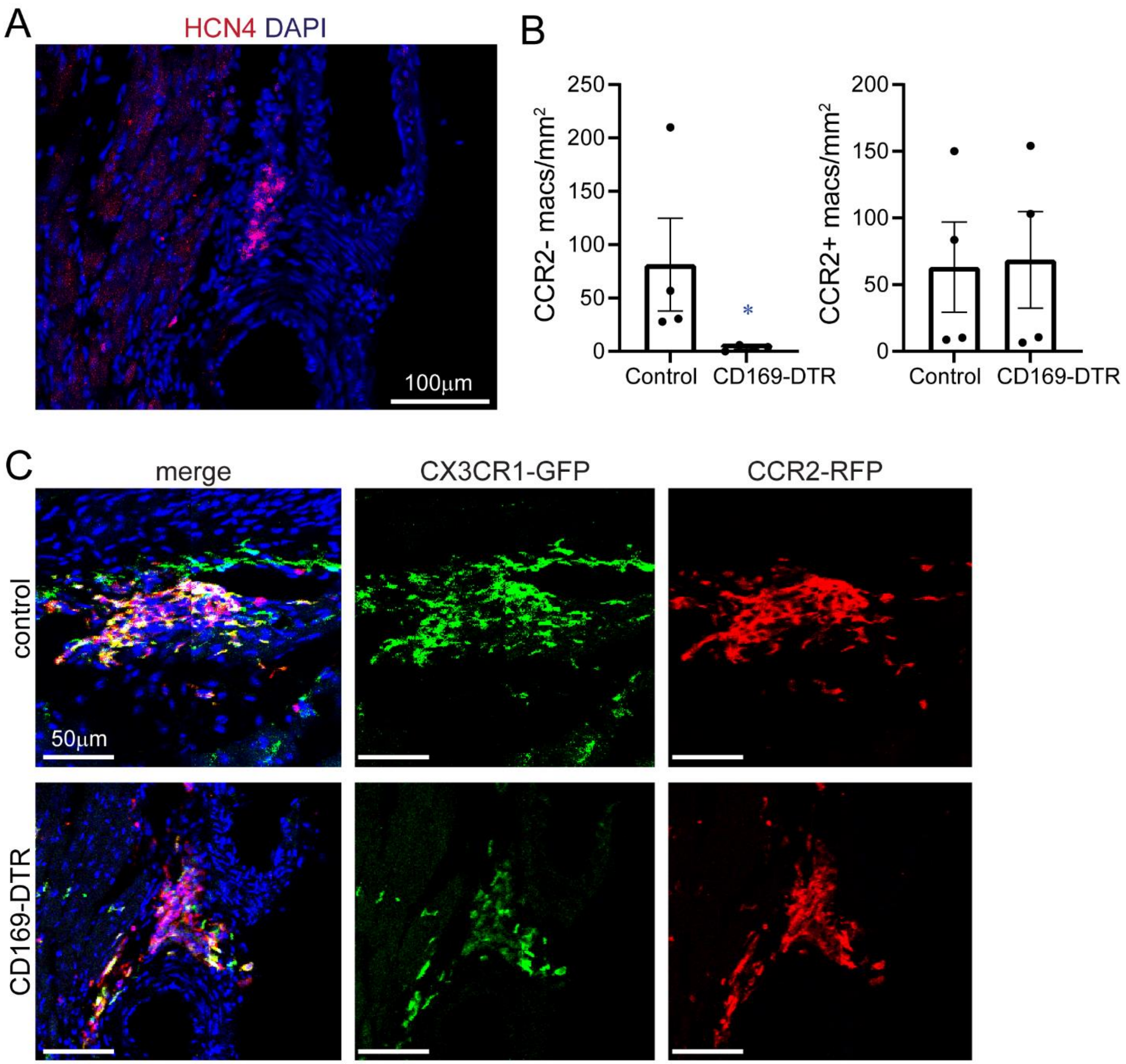


**Figure S10** (related to Figure 4). **Macrophage ablation in the atrioventricular node. A**, Representative image of HCN4 immunostaining (red) showing staining in the atrioventricular node. DAPI: blue. **B**, Quantification of CCR2- and CCR2+ macrophages in the atrioventricular node of control (CX3CR1-GFP/+ CCR2-RFP/+) and CD169-DTR (CD169-DTR CX3CR1-GFP/+ CCR2-RFP/+) mice. * denotes $p<0.05$ compared to controls. Each data point represents a biologically independent replicate (n=4). **C**, Immunostaining for GFP and RFP in the atrioventricular node of control (CX3CR1-GFP/+ CCR2-RFP/+) and CD169-DTR (CD169-DTR CX3CR1-GFP/+ CCR2-RFP/+) mice. The atrioventicular node region was identified in serial sections using HCN4 staining. All immunostaining studies were performed in 4 independent specimens.

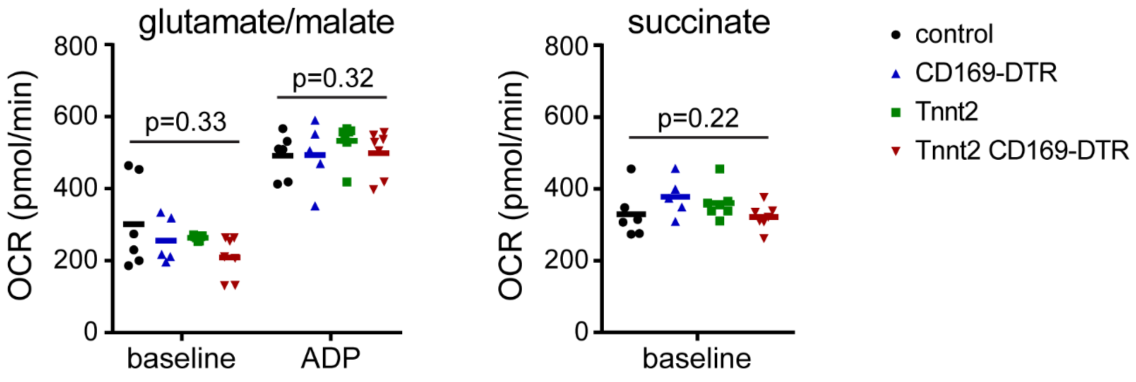


**Figure S11** (related to Figure 4). **Mitochondrial respiration.** Oxygen consumption rate (OCR) in mitochondria isolated from the hearts of control, CD169-DTR, Tnnt2$^{\Delta K210/\Delta K210}$, and Tnnt2$^{\Delta K210/\Delta K210}$ CD169-DTR 3 weeks of DT treatment. Mitochondria were provided either glutamate/malate or succinate substrates. ADP was administered to measure maximal OCR. Each data point represents mitochondria preparations from an individual hearts. n=5-6 mice per experimental group.

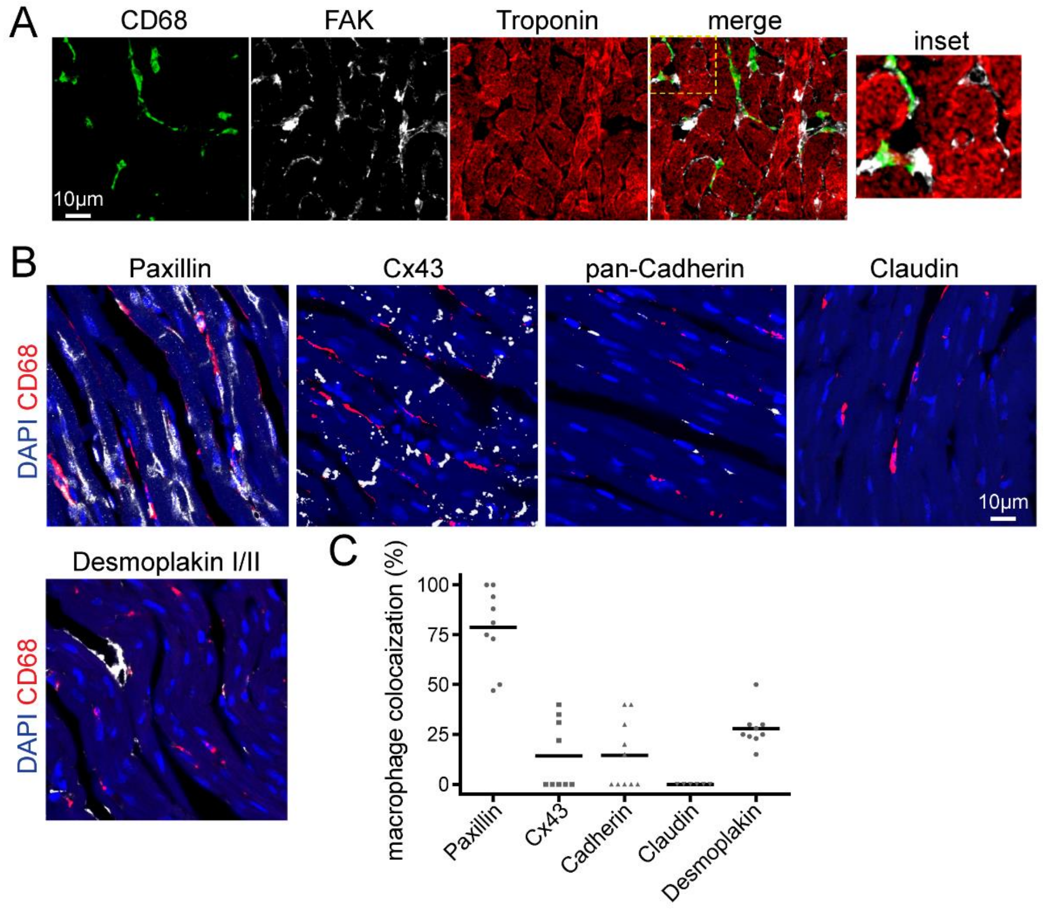


**Figure S12** (related to Figure 6). **Interactions between macrophages and cardiomyocytes. A**, Immunostaining for CD68 (green), FAK (white), Troponin (red) in Tnnt2$^{\Delta K210}$ hearts. **B,** Immunostaining of cell junction markers at sites of macrophage and cardiomyocyte interactions. Blue-DAPI, red-CD68, white-cell junction markers: Paxillin (focal adhesion complex) Cx43 (gap junction), pan-Cadherin (adherins junction), Claudin (tight junction), Desmoplakin I/II (desmosome). **C**, Quantification of the percent co-localization between macrophages and the indicated cell junction marker. Each data point represents an individual animal. n=7-9 mice per experimental group.

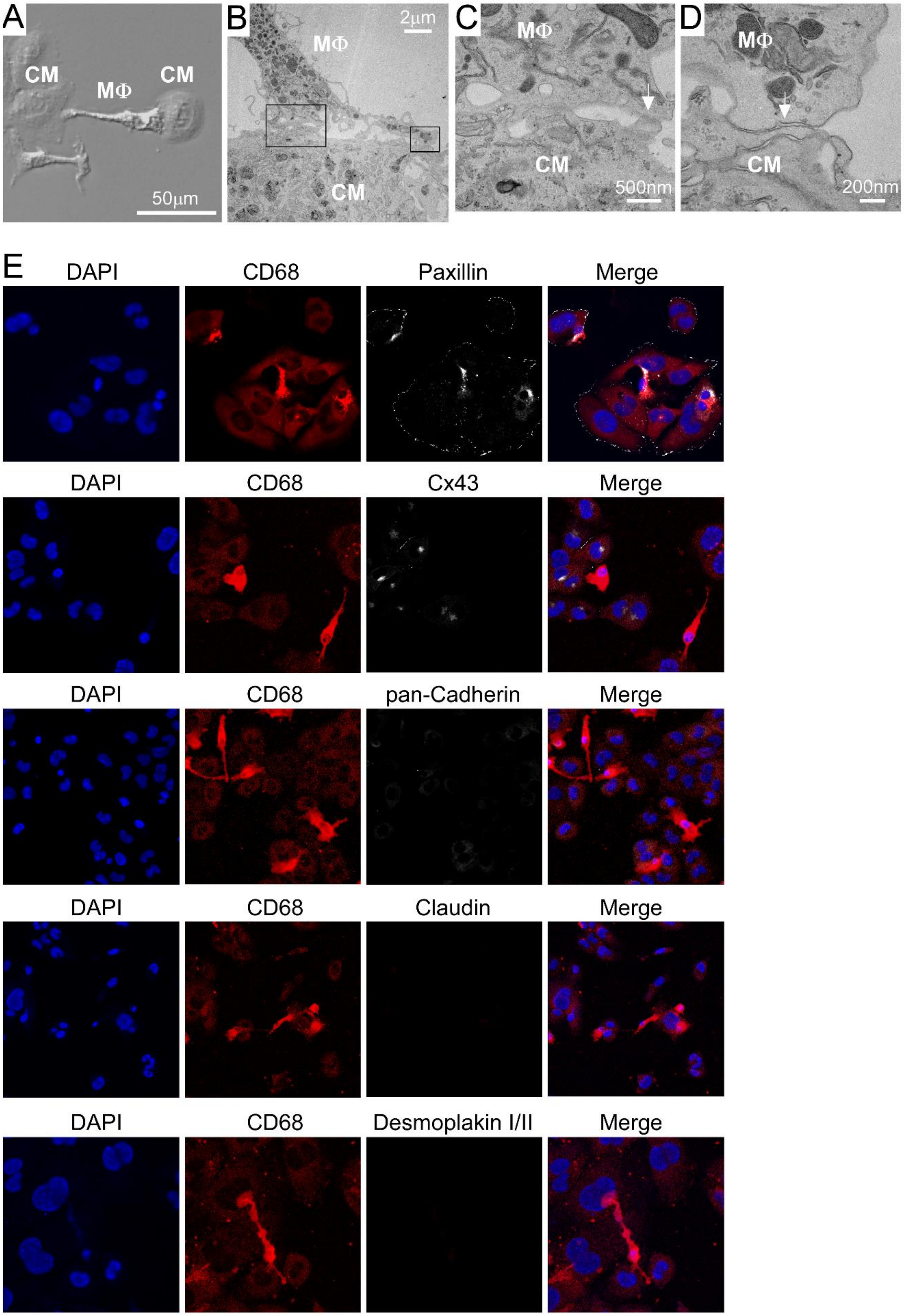
A
CM
MΦ
CM
50μm
B
2μm
MΦ
CM
C
MΦ
CM
500nm
D
MΦ
CM
200nm
E
DAPI
CD68
Paxillin
Merge
DAPI
CD68
Cx43
Merge
DAPI
CD68
pan-Cadherin
Merge
DAPI
CD68
Claudin
Merge
DAPI
CD68
Desmoplakin I/II
Merge

**Figure S13** (related to Figure 6). **Macrophages and cardiomyocytes spontaneously and physically interact in vitro. A-D**, Electron microscopy of bone marrow-derived macrophages (MΦ) and HL1 cardiomyocytes (CM) after 4 hours of co-culture demonstrating physical interactions (white arrows) between macrophages and cardiomyocytes. C-D: high magnification of boxed areas shown in B. **E**, Immunostaining of cell junction markers at sites of macrophage and cardiomyocyte interactions. Blue-DAPI, red-CD68, white-cell junction markers: Paxillin (focal adhesion complex) Cx43 (gap junction), pan-Cadherin (adherins junction), Claudin (tight junction), Desmoplakin I/II (desmosome). Representative images from 4 independent experiments.

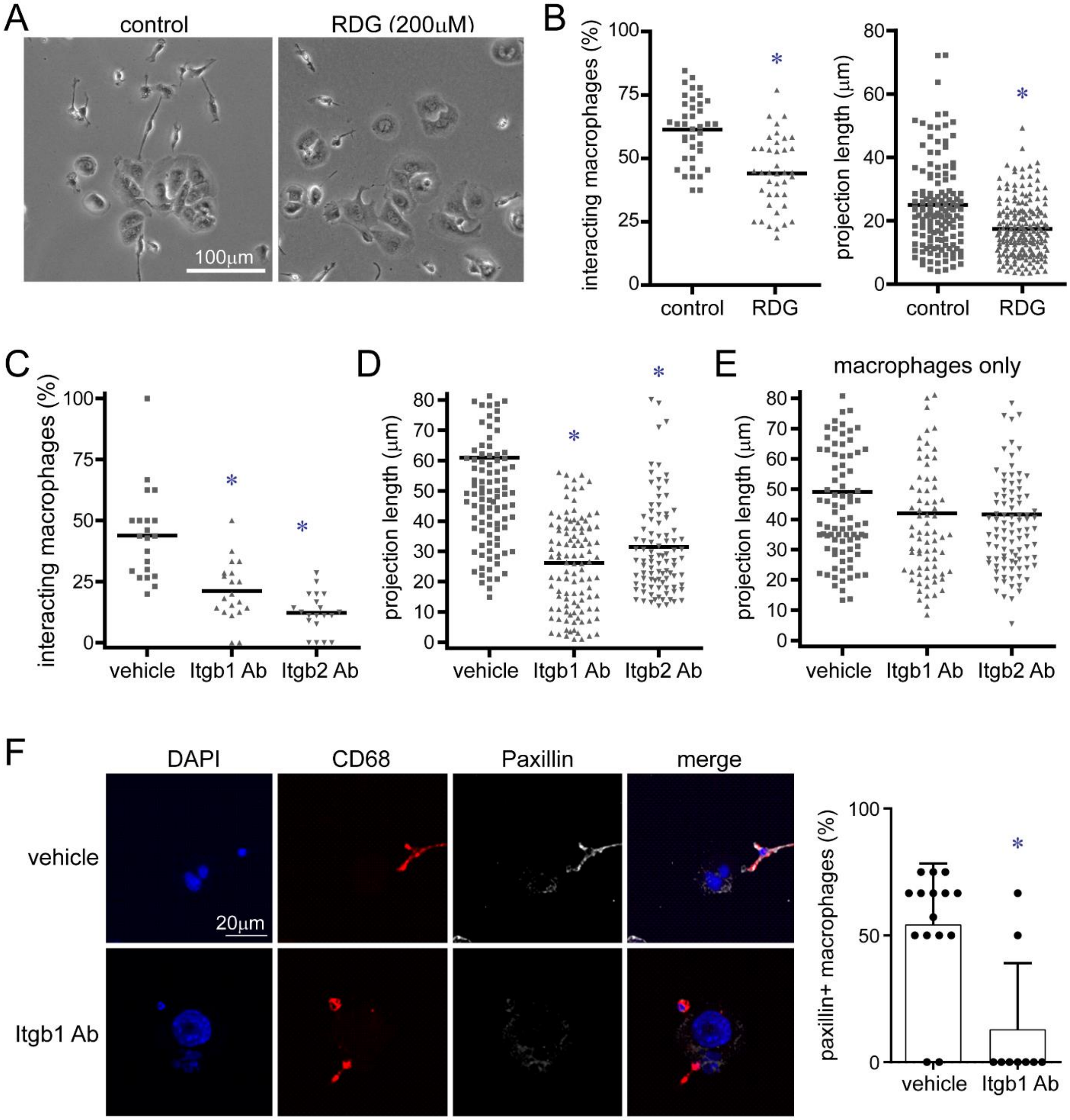

A
control
RDG (200μM)
100μm
B
interacting macrophages (%)
100
75
50
25
0
control
RDG
*
projection length (μm)
80
60
40
20
0
control
RDG
*
C
interacting macrophages (%)
100
75
50
25
0
vehicle
Itgb1 Ab
Itgb2 Ab
*
*
D
projection length (μm)
80
70
60
50
40
30
20
10
0
vehicle
Itgb1 Ab
Itgb2 Ab
*
*
E
macrophages only
projection length (μm)
80
70
60
50
40
30
20
10
0
vehicle
Itgb1 Ab
Itgb2 Ab
F
DAPI
CD68
Paxillin
merge
vehicle
Itgb1 Ab
20μm
paxillin+ macrophages (%)
100
50
0
vehicle
Itgb1 Ab
*

**Figure S14** (related to Figure 6). **β-integrins contribute to interactions between macrophages and cardiomyocytes in vitro. A**, Differential interference contrast (DIC) microscopy images of bone marrow-derived macrophages and HL1 cardiomyocytes co-cultured for 4 hours in the presence of vehicle to RDG peptides. **B**, Quantification of the percent of macrophages interacting with cardiomyocytes and macrophage projection length. * denotes $p<0.05$ compared to vehicle control (Mann-Whitney test). **C-D**, Quantification of the percent of macrophages interacting with cardiomyocytes and macrophage projection length in co-cultures treated with either isotype antibody (vehicle), Itgb1 neutralizing antibody, or Itgb2 neutralizing antibody. **E**, Quantification of projection length in bone marrow-derived macrophages cultured independently in the presence of isotype antibody (vehicle), Itgb1 neutralizing antibody, or Itgb2 neutralizing antibody. * denotes $p<0.05$ compared to control (AVOVA, post-hoc Tukey). **F**, CD68 (red) and Paxillin (white) immunostaining showing loss of Paxillin expression in cultured macrophages treated with the neutralizing Itgb1 antibody. Blue-DAPI. Representative images from 4 independent experiments. * denotes $p<0.05$ compared to vehicle control (Mann-Whitney test). Each data point in B-F represents information compiled from 4 experimental replicates. Experiments were repeated 4 times and all data was incorporated into the displayed graphs.

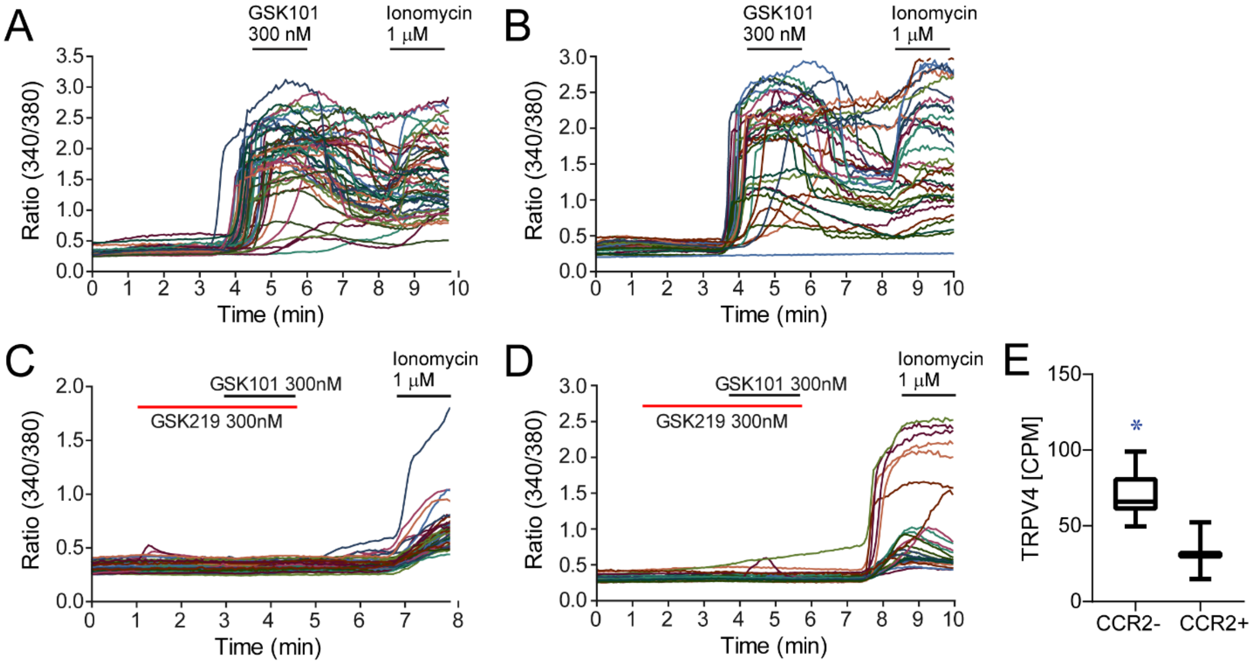


**Figure S15** (related to Figure 7). **TRPV4 channel activity in cardiac CCR2- and CCR2+ macrophages. A-B**, Ratiometric calcium assays showing that CCR2- (A) and CCR2+ (B) macrophages isolated from the adult mouse heart by flow cytometry express active TRPV4 channels. GSK101: TRPV4 agonist, Ionomycin: calcium ionophore. **C-D**, Ratiometric calcium assays showing that the TRPV4 antagonist (GSK219) blocks rises in intracellular calcium induced by GSK101 in cardiac CCR2- (C) and CCR2+ (D) macrophages. Each tracing represents an independently analyzed cell. Tracings were compiled from 3 independent experiments. **E**, TRPV4 mRNA expression in CCR2- and CCR2+ cardiac macrophages. RNAseq data derived from Immgen (Immunological Genome, 2020). CPM: counts per million. * denotes $p<0.05$ compared to CCR2+ macrophages (Mann-Whitney test). Data obtained from 5 biologically independent samples per experimental group.

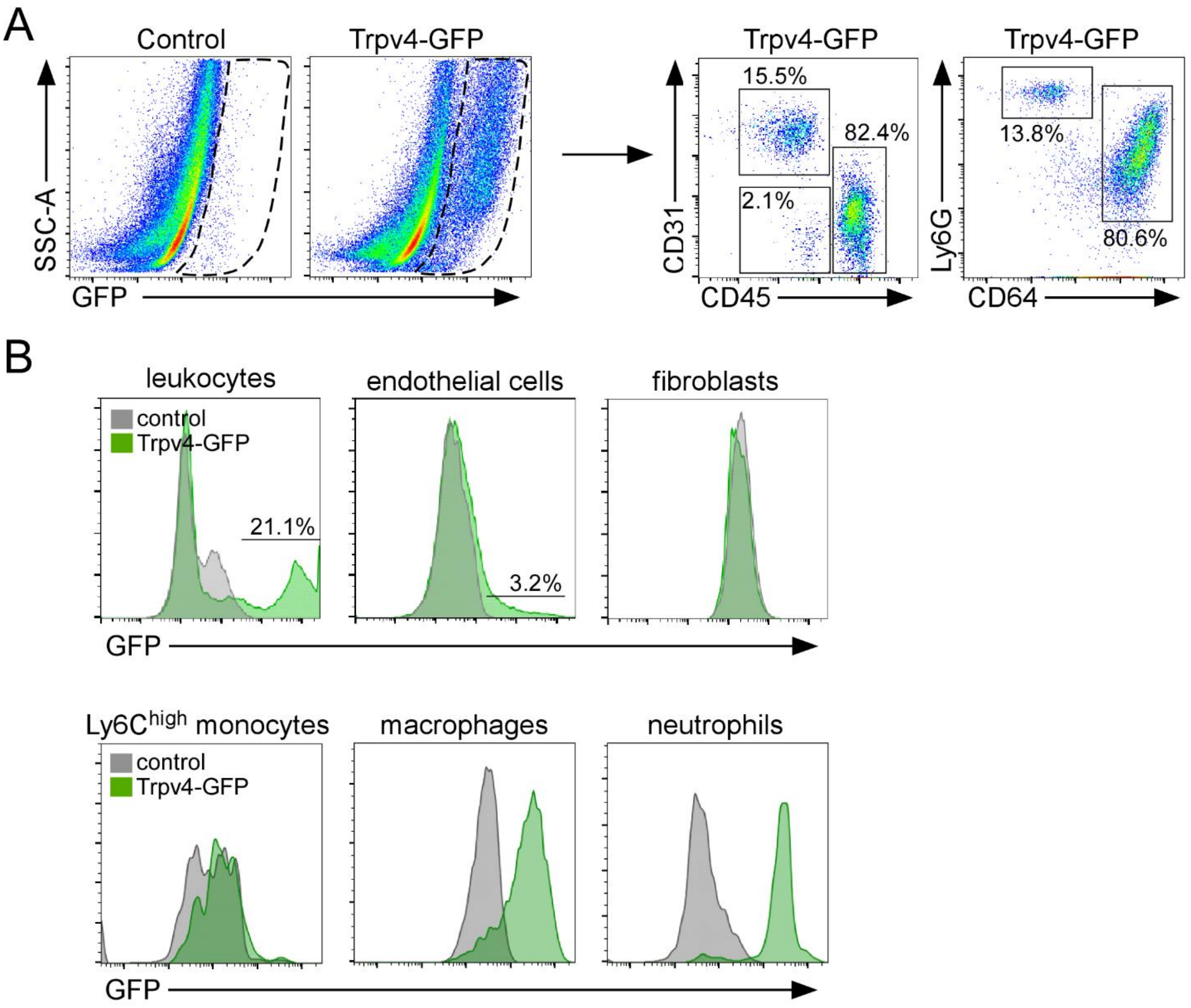


**Figure S16** (related to Figure 7). **TRPV4 expression in cardiac macrophages and neutrophils. A**, Flow cytometry of TRPV4-GFP BAC transgenic hearts showing GFP expression in leukocytes (CD45+), endothelial cells (CD31+CD45-), neutrophils (Ly6G+CD64-,) and macrophages (CD64+Ly6G-). Percentages of each cell type as a function of the number of GFP+ cells is shown. **B**, Histograms showing GFP mean florescent intensity of CD45+ leukocytes, CD31+CD45- endothelial cells, MEFSK4+CD45-CD31- fibroblasts, Ly6C$^{high}$ monocytes, CD64+Ly6C$^{low}$ macrophages, and Ly6G+ neutrophils isolated from littermate controls (grey) and TRPV4-GFP (green) hearts. Percentages of GFP+ leukocytes and endothelial cells are shown as a function of total number of leukocytes and endothelial cells, respectively. Data obtained from 3 independent experiments.

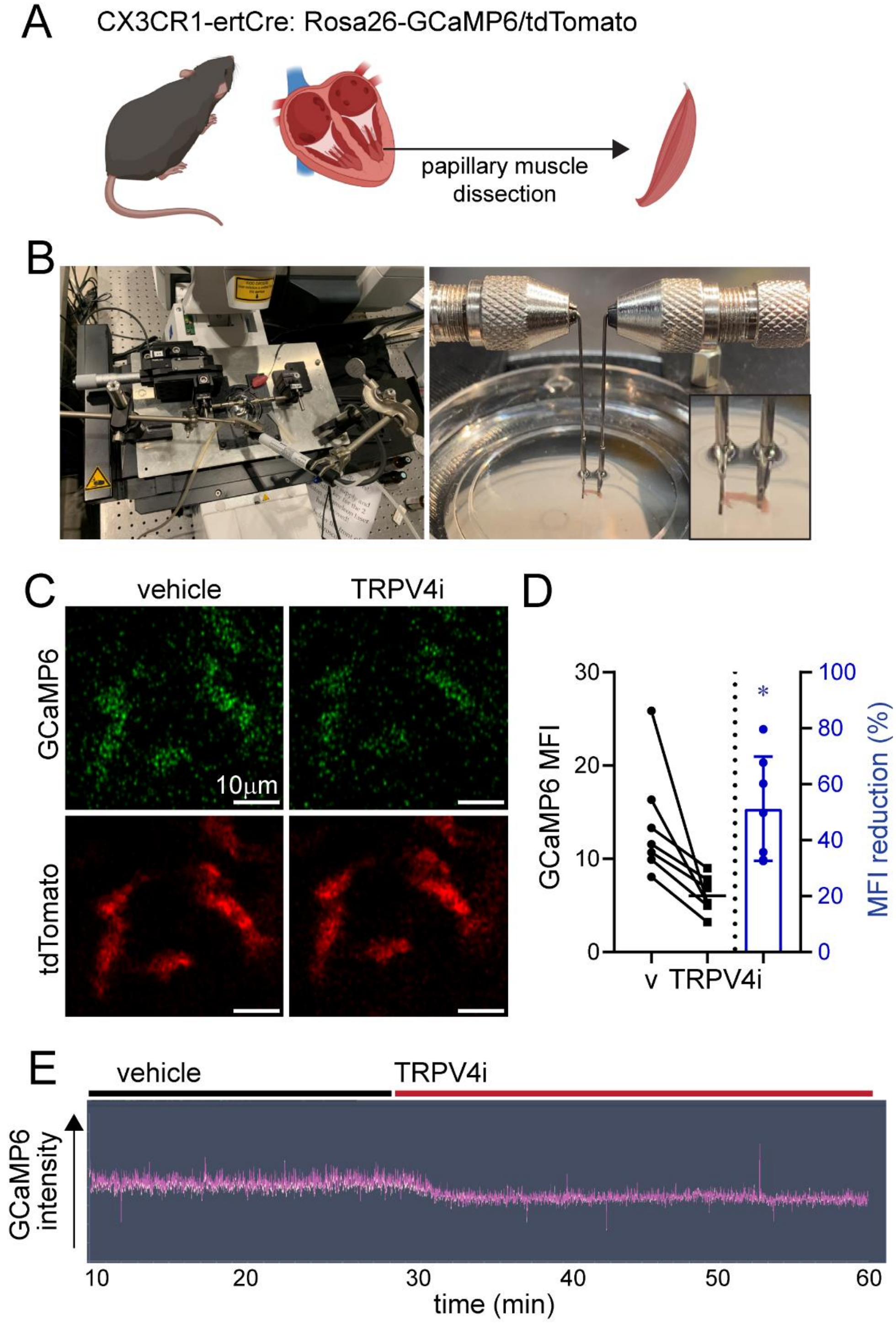
A
CX3CR1-ertCre: Rosa26-GCaMP6/tdTomato
papillary muscle
dissection
B
C
vehicle
TRPV4i
GCaMP6
tdTomato
10μm
D
GCaMP6 MFI
MFI reduction (%)
v
TRPV4i
E
vehicle
TRPV4i
GCaMP6 intensity
time (min)

**Figure S17** (related to Figure 7). **TRPV4 activity in cardiac macrophages *in situ*. A**, Schematic of mouse strain used (CX3CR1-ertCre; Rosa26-GCaMP6/tdTomato) and isolation of LV papillary muscles. **B**, Photograph of the 2-photon imaging stage and apparatus used to apply axial tension to papillary muscle preparations. **C**, 2-photon imaging of GFP (green) and tdTomato (red) in papillary muscle preparations harvested from CX3CR1-ertCre; Rosa26-GCaMP6/tdTomato mice treated with either vehicle or the TRPV4 inhibitor GSK219 (TRPV4i). **D**, Quantification of GCaMP6 signal. Each data point represent mean data from an individual experiment (n=6). * denotes $p<0.05$ compared to vehicle. **E**, Representative intensity graph of GCaMP6 signal over time (minutes) during infusion of vehicle (black bar) and TRPV4i (red bar) into the circulating media bath.

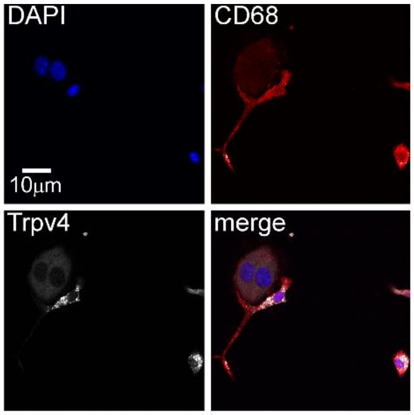


**Figure S18** (related to Figure 7). **TRPV4 expression in bone marrow-derived macrophages.** Immunostaining for CD68 (red) and TRPV4 (white) in bone marrow-derived macrophages co-cultured with HL1 cardiomyocytes. Blue: DAPI. Representative images from 4 independent experiments.

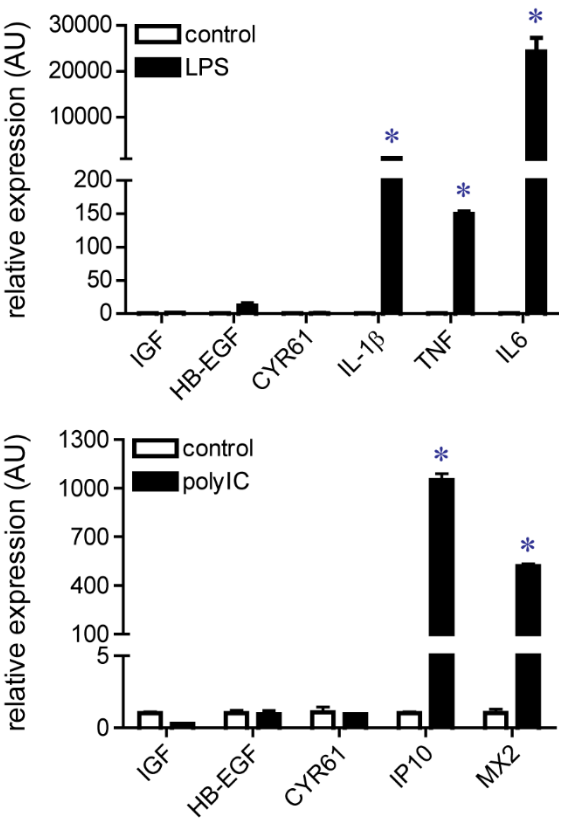


**Figure S19** (related to Figure 7). **Activators of MYD88 and TRIF signaling do not stimulate macrophage growth factor expression.** Quantitative RT-PCR assays of bone marrow-derived macrophages stimulated with vehicle control, LPS, or polyIC. * denote $p<0.05$. Error bars represent standard deviation. n=4 independent experiments.

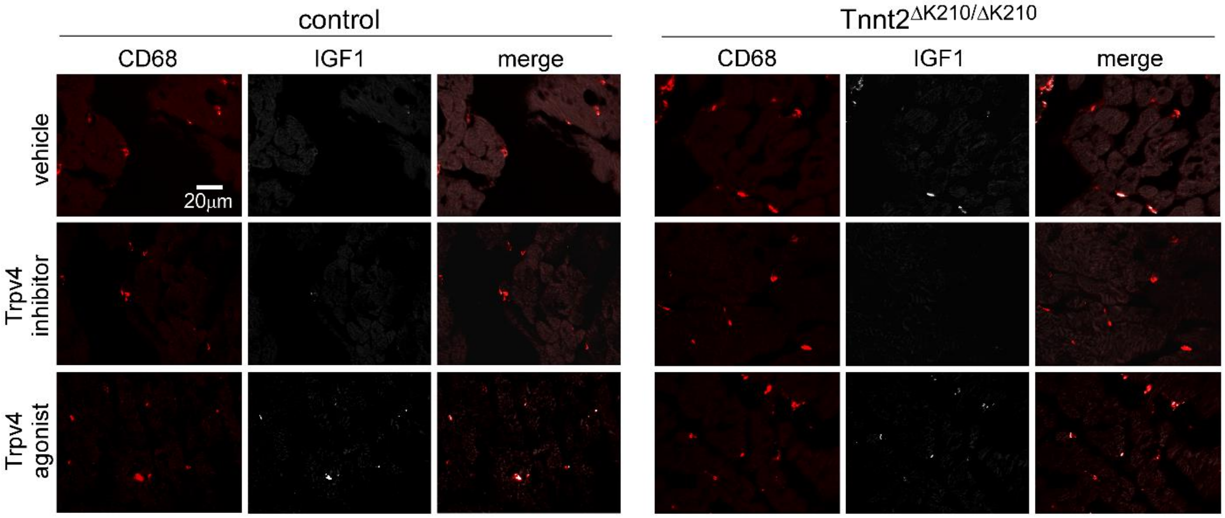


**Figure S20** (related to Figure 8). **TRPV4 channel activity regulates IGF1 expression in CCR2- macrophages.** Immunostaining for IGF1 (white) and CD68 (red) in the LV myocardium of control and Tnnt2$^{\Delta K210/\Delta K210}$ mice treated with either vehicle, TRPV4 inhibitor, or TRPV4 agonist demonstrating that TRPV4 channel activity regulates macrophage IGF1 protein expression in vivo. Representative images from 4 mice per experimental group.
